# The genetic basis of apple shape and size unraveled by digital phenotyping

**DOI:** 10.1101/2023.08.25.554767

**Authors:** Beat Keller, Michaela Jung, Simone Bühlmann-Schütz, Marius Hodel, Bruno Studer, Giovanni AL Broggini, Andrea Patocchi

## Abstract

Great diversity of shape, size, and skin color is observed among the fruits of different apple genotypes. These traits are critical for consumers and therefore interesting targets for breeding new apple varieties. However, they are difficult to phenotype and their genetic basis, especially for fruit shape and ground color, is largely unknown. We used the fruit FruitPhenoBox to digitally phenotype 506 genotypes of the apple reference population (apple REFPOP) genotyped for 303,148 single nucleotide polymorphism (SNP) markers. From the apple images, 573 highly heritable features describing fruit shape and size as well as 17 highly heritable features for fruit skin color were extracted to explore genotype-phenotype relationships. Out of these features, nine and four principal components (PCs) as well as 16 and eight uncorrelated features were chosen to carry out genome-wide association studies for fruit shape, size, and fruit skin color, respectively. In total, 69 SNPs scattered over all 17 apple chromosomes were significantly associated with round, conical, cylindrical, or symmetric fruit shapes and fruit size. Novel associations with major effect on round or conical fruit shapes and fruit size were identified on chromosomes 1 and 2. Additionally, 16 SNPs associated with PCs and uncorrelated features related to red over color as well as green and yellow ground color were found on eight chromosomes. The identified associations can be used to advance marker-assisted selection in apple fruit breeding to systematically select for desired fruit appearance.

## 1 Introduction

Apple (*Malus* × *domestica* Borkh.) cultivars inherited their great diversity of fruit qualities and sizes from its wild ancestors (Cornille et al. 2014). During domestication, the apple fruits have increased in size and gained other fruit qualities, increasingly reflecting human consumer preferences (Cornille et al. 2014; Duan et al. 2017). Today’s cultivated apple is the third most produced fruit crop in the world (http://www.fao.org/faostat). Its fruit appearance, that can be described as the external fruit quality including traits such as fruit shape, size, and skin color, has been found to be a primary purchase-driving criterion for consumers (Kays 1999; Musacchi and Serra 2018).

To satisfy consumer preferences for specific fruit size, information about genetic regions controlling the trait can benefit apple breeding. Genes potentially contributing to the increase in fruit size during apple speciation before its domestication colocalizing with a quantitative trait locus for fruit size on chromosome (Chr) 11 have been identified (Duan et al. 2017; Yao et al. 2015). Additionally, a locus on Chr 15 was associated with fruit vertical diameter of wild and cultivated *Malus* accessions (Liao et al. 2021), and two loci on Chr 8 and 15 were found to be significantly associated with fruit weight in *M.* × *domestica* (Devoghalaere et al. 2012). In the same two genetic regions, selective sweeps in *M.* × *domestica* compared with *M. sieversii* and *M. sylvestris* were reported (Duan et al. 2017). The genetic basis of fruit size in *M.* × *domestica* has been further studied on traits such as fruit weight, diameter, and length reporting many quantitative trait loci (QTL) across all apple chromosomes (Chang et al. 2014; Costa 2015; Devoghalaere et al. 2012; Kenis et al. 2008; Liu et al. 2016; Minamikawa et al. 2021). Despite the high abundance of known associations with traits related to fruit size, additional discovery, validation, and consolidation of the identified associations could facilitate future application in genomics-assisted breeding.

The quantitative inheritance of fruit shape in apple was postulated already in 1960 (Brown 1960). Fruit shape as a subjective score of fruit roundness appeared under strong genetic control (Hardner et al. 2016). Description of fruit shape as a ratio between fruit length and diameter resulted in discovery of QTL for length/diameter on multiple linkage groups (Chang et al. 2014; Sun et al. 2012). In tomato, various phenotypic measurements of fruit shape based on fruit boundaries (contours) automatically extracted using the Tomato Analyzer have resulted in the detection of several QTL for fruit shape (Brewer et al. 2006; Gonzalo et al. 2009). Similar digital phenotyping approach applied in apple may help further elucidate the genetic background of apple fruit shape.

The causal loci of red fruit skin in apples have been broadly studied in the past. Several transcription factors control anthocyanin production in the fruit skin during ripening (Espley et al. 2007; Takos et al. 2006). A major QTL for red over color has been repeatedly reported on Chr 9 at around 30 Mbp (Chagné et al. 2016; Duan et al. 2017; Moriya et al. 2017). However, the genomic control of green and yellow fruit skin color was rarely studied in apple. Visually assessed ground color, a gradient between green and yellow fruit skin color, was associated with QTL distributed over 13 chromosomes (Jung et al. 2022). Jung et al. (2022) further demonstrated that green color estimated digitally using an automated fruit sorting machine was an inverse of the digitally and visually estimated red over color, and all these skin color traits were associated with the same single nucleotide polymorphism (SNP) marker on Chr 9 at 33.8 Mbp. Furthermore, SNPs of minor effects associated with green color were identified on Chr 6, 10 and 17, none of the SNPs associated with green color overlapped with the SNPs reported for ground color, and no SNPs were reported for yellow color (Jung et al. 2022). Additional phenotypic information for green and yellow skin color may allow for a more detailed description and the advancement of biological understanding of these traits.

Digital phenotyping is increasingly used in fruit trees to ease quantification of traits (Huang et al. 2020). In apple trees, RGB images have been used to estimate tree architecture or count fruits on trees in the orchard (Zhang et al. 2023; Zine-El-Abidine et al. 2021). The recently developed phenotyping device FruitPhenoBox, an RGB camera system combined with automatized image analysis, showed as a low-cost tool suitable for extraction of fruit features (Kirchgessner et al. 2023). However, the application of fruit imaging for the assessment of fruit shape, size, and skin color with the aim of identifying genetic associations with these traits remained to be tested.

To dissect the genetic basis of apple fruit shape, size, and skin color, this study used the FruitPhenoBox (Kirchgessner et al. 2023) to digitally phenotype the apple REFPOP, a diverse apple reference population located at Agroscope in Waedenswil, Switzerland (Jung et al. 2020). We aimed to (i) extract new phenotypic features from images taken with the FruitPhenoBox, (ii) compare the features with apple fruit size and skin color traits obtained visually and using a commercial sorting machine, (iii) perform genome-wide associations studies (GWAS) on heritable features, and (iv) identify genetic markers and haplotypes that could be used for genomics-assisted breeding towards the improvement of apple fruit appearance.

## 2 Material and methods

### 2.1 Plant material and phenotyping

Fruits of the apple REFPOP, which was planted in 2016 at Waedenswil, Switzerland, were used to study fruit shape, size, and skin color. The 269 accessions and 265 progeny of the apple REFPOP were characterized by 303,148 SNP markers (Jung et al. 2020). The genotypes were replicated at least twice in a complete randomized block design. All fruits of each genotype replicate (tree) were manually harvested. Harvest date was defined for each tree separately as the date when more than 50% of all fruits reached their full physiological maturity (Jung et al. 2020). The FruitPhenoBox (Kirchgessner et al. 2023) was used to phenotype 175 and 506 genotypes harvested in 2019 and 2020, respectively. The phenotyping was performed on harvest date at the level of individual trees. When more than five fruits per tree were available, a random sample of five representative fruits was chosen for imaging. When five or fewer fruits were produced by a tree, all available fruits were imaged. Individual fruits with shape or size clearly different from the rest of the group, or diseased, deformed, or damaged fruits, were excluded before sampling and imaging.

To compare measurements obtained using the FruitPhenoBox with different measurement approaches, a set of visually scored traits and traits measured by a commercial sorting machine (GREEFA iQS4 v.1.0) in 2019 and 2020 available from Jung et al. (2022) were included in the study as supplementary traits. The visually scored supplementary traits ground color (labeled as Color_ground) and percentage of red over color (Color_over) were measured as described by Jung et al. (2022). The supplementary traits measured by the sorting machine were comprised of the percentage of green (labeled as Green), yellow (Yellow), and red fruit skin color (Red), fruit diameter (Diameter), length (Length), maximum size (maxSize), volume (Volume) and single fruit weight (Fruit_weight_single). For each of the supplementary traits measured by the sorting machine, the values were averaged across all fruits of a tree.

### 2.2 Imaging with the FruitPhenoBox

The FruitPhenoBox, its imaging procedure, and image segmentation were described by Kirchgessner et al. (2023). Briefly, the FruitPhenoBox consisted of five RGB cameras and a scale. Four cameras were placed in the corners of the FruitPhenoBox, and one camera was positioned above the fruit (Figure 1A). For imaging, a single fruit was placed on a platform on the scale. By a manual command, the weight was measured, and one image of the fruit was taken by each of the five cameras. The background of each image was identified and removed using image segmentation implemented in MATLAB (MATLAB 2010) as described by (Kirchgessner et al. 2023). The apple contour was derived in Cartesian and polar coordinates. The polar coordinates were expressed as radius and polar angle for 360 points, one point for each degree. The Cartesian coordinates were expressed as 360 corresponding x and y values. The color of the segmented fruit was derived as hue and saturation values ranging from 0 to 1 representing 36 averaged values among the color space over all segmented pixels.

**Figure 1:**
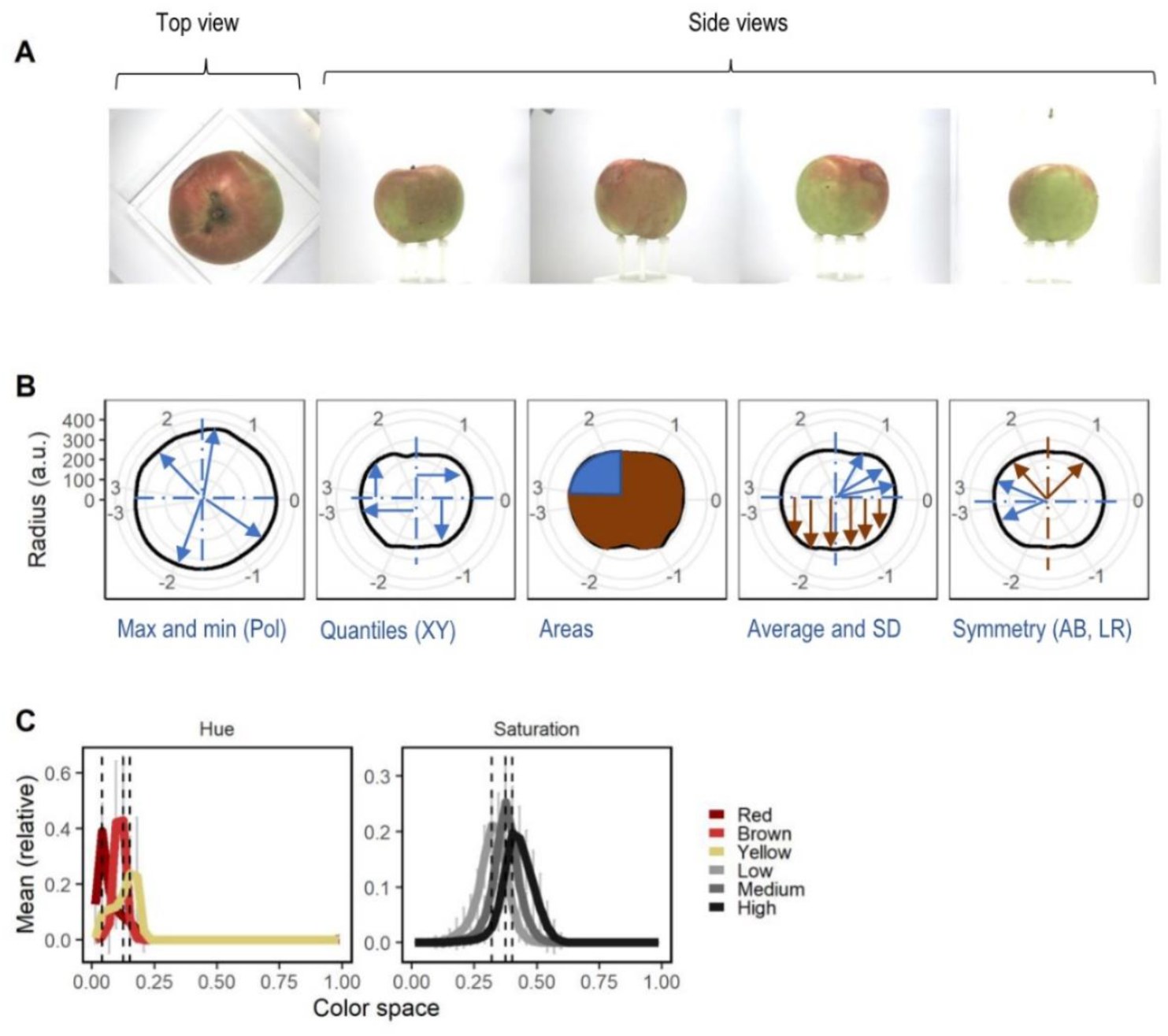
Example of feature definition with the FruitPhenoBox. A: One top and four side view RGB images were taken for each fruit. B: For shape and size feature definition, fruit contour was derived in polar (Pol) and Cartesian (XY) coordinates for each fruit and image, displayed here as radius in absolute values (a.u.). Features such as maximum and minimum coordinate values (Max and min), quantiles, areas, averages, and standard deviations (SD) as well as symmetries were extracted either from the full fruit contour or the top (A), bottom (B), left (L) or right (R) fruit sections or their combinations. C: For the color features, histograms were derived for hue and saturation color spaces. Color features were extracted at the positions of peaks obtained for hierarchically clustered genotypes.

Apple fruits whose dimension on the side-view images was smaller than 150 pixels were not further considered because they did not fully cover the opposite camera lens and caused the image segmentation to fail. To fully remove the apple stalk in the apple side-view images, data points above 1.5 times the interquartile range (the difference between the first and the third quartile) of the upper quarter of the contour data were removed.

### 2.3 Feature extraction for shape and size

From the apple fruit contours obtained from FruitPhenoBox, one-dimensional features for shape and size were systematically processed (Table 1, Figure 1B). For the points on the fruit contour defined in polar coordinates (Pol), the distance of the point from the center of the coordinate system, i.e., the radius, was used in feature extraction. The points in the Cartesian coordinate system were expressed as individual coordinates for both axes (X measuring fruit diameter, Y measuring fruit length). Different fruit sections were determined as the top (A), bottom (B), left (L) or right (R) half of the fruit, as well as intersections of the fruit halves, i.e., fruit quarters (A.L, A.R, B.L, B.R) and unions of the halves, i.e., the full fruit (AB, LR). Points from the fruit contour derived in Cartesian and polar coordinates within a chosen fruit section were used to calculate different types of features, namely the mean (Av), the standard deviation (SD), the minimum (Min) or maximum (Max) value, the 0.25, 0.50 or 0.75 quartile (Q25, Q50, Q75), the area, the ratio between Max and Min value (RatM), and the Max value relative to the Av value (MaxR). The area was calculated by summing up 1°-radius increments in the specified section. For each type of feature, the absolute value (Abs) was taken from the features extracted on different sections. For the opposite fruit halves, absolute values were first extracted and then the ratio (Rat) and the sum (Sum) were calculated across the fruit halves. The ratio or the sum of top vs. bottom halves was calculated as average (Rat.Av and Sum.Av) and SD (Rat.SD and Sum.SD). Hereby, the top and bottom halves were calculated from quarter values using the corresponding left-right fruit quarters (AB.LR). Additionally, symmetry between L and R as well as A and B was calculated as the Av or SD of the difference between corresponding absolute values of radius of each fruit half (Sym.abs, sections LR and AB). Similarly, the symmetry measure Sym.abs was obtained for individual apple sections A, B, L and R when comparing the opposite quarters within the fruit halves. Additionally, a relative symmetry (Sym.rel) was derived for the same apple sections as Sym.abs, but the measure Sym.rel was normalized for the area of the apple section.

**Table 1:** Feature definition key. Names of the extracted features represent combinations of the different row entries over the columns from left to right. For the shape and size feature definition, the FruitPhenoBox was used to extract fruit contours from images of individual apples. Mean (Av) or standard deviation (SD) was calculated over up to five apples per tree that were imaged with one camera from the top view (Top) and four side-view cameras. The four side-view images were either averaged (side average, SAv) or their SD was obtained (side SD, SSD). The apple contour was defined by points expressed in polar (Pol) or Cartesian coordinates. The points in the Cartesian coordinate system were defined as individual coordinates for two axes (X measuring fruit diameter, Y measuring fruit length). Different fruit sections were determined as the top (A), bottom (B), left (L) or right (R) half of the fruit, fruit quarters (A.L, A.R, B.L, B.R) and full fruits (AB, LR). Points from the fruit contour within a chosen fruit section were used to calculate different types of features, namely the mean (Av), the standard deviation (SD), the minimum (Min) or maximum (Max) value, the 0.25, 0.50 or 0.75 quartile (Q25, Q50, Q75), the area, the ratio between Max and Min value (RatM), the Max value relative to the Av value (MaxR) and the symmetry features (Sym.abs, Sym.res). To calculate each type of feature, the absolute value (Abs) was taken from the features extracted on different fruit sections, or the absolute values were first extracted for the opposite fruit halves, and then the ratio (Rat) and the sum (Sum) were calculated across the fruit halves. The ratio or the sum of top vs. bottom halves was calculated as average (Rat.Av, Sum.Av) and SD (Rat.SD, Sum.SD). Hereby, the top and bottom halves were calculated from quarter values using the corresponding left-right fruit quarters (AB.LR). For the color features, histograms for hue (Hue) and saturation (Sat) color spaces were used to define features for red, brown, and yellow hue as well as low, medium, and high saturation.

| Tree-level | Fruit-level | Data type | Calculation | Section | Feature type |
| --- | --- | --- | --- | --- | --- |
| Mean (Av) | Top | Pol | Abs | A.L | Av |
| SD | Side average (SAv) |  |  | A.R | SD |
|  | Side SD (SSD) |  |  | B.L | Max |
|  |  |  |  | B.R | Min |
|  |  |  |  |  | Q25 |
|  |  |  |  |  | Q50 |
|  |  |  |  |  | Q75 |
|  |  |  |  |  | Area |
|  |  |  |  |  | RatM |
| Mean (Av) | Top | Pol | Av | A | Av |
| SD | Side average (SAv) |  | SD | B | SD |
|  | Side SD (SSD) |  |  |  | Max |
|  |  |  |  |  | Min |
|  |  |  |  |  | Q25 |
|  |  |  |  |  | Q50 |
|  |  |  |  |  | Q75 |
|  |  |  |  |  | Area |
|  |  |  |  |  | RatM |
| Mean (Av) | Top | Pol | Abs | AB | Av |
| SD | Side average (SAv) |  | Sum.SD | AB.LR | SD |
|  | Side SD (SSD) |  | Sum.Av |  | Max |
|  |  |  | Rat.SD |  | Min |
|  |  |  | Rat.Av |  | Q25 |
|  |  |  |  |  | Q50 |
|  |  |  |  |  | Q75 |
|  |  |  |  |  | Area |
|  |  |  |  |  | RatM |
| Mean (Av) | Top | Pol | Av | LR | Sym.rel |
| SD | Side average (SAv) |  | SD | AB | Sym.abs |
|  | Side SD (SSD) |  |  |  |  |
| Mean (Av) | Top | Pol | Av | A | Sym.rel |
| SD | Side average (SAv) |  | SD | B | Sym.abs |
|  | Side SD (SSD) |  |  | L |  |
|  |  |  |  | R |  |
| Mean (Av) | Top | X | Abs | A | Av |
| SD | Side average (SAv) | Y |  | B | SD |
|  | Side SD (SSD) |  |  |  | Max |
|  |  | XY | Sum | A | Min |
|  |  |  | Rat | B | Q25 |
|  |  |  |  |  | Q50 |
|  |  |  |  |  | Q75 |
|  |  |  |  |  | RatM |
|  |  |  |  |  | MaxR |
| Mean (Av) | Side average (SAv) | Hue | Abs | AB | Red |
| SD | Side SD (SSD) |  |  |  | Yellow |
|  |  |  |  |  | Brown |
|  |  | Sat | Abs | AB | Low |
|  |  |  |  |  | Medium |
|  |  |  |  |  | High |

One image from top view (Top) and four images from side view (Side) were taken for each fruit. The features were corrected for camera effects in order to remove potential technical error due to the camera settings and position. Phenotypes (*y_ijkl_*) of the *i^th^* genotype, the *j^th^* year and the *k^th^* replicate (tree) taken by the *l^th^* camera were corrected with a fixed effect for the camera (*C_l_*_(*j*)_) nested in year using the model:

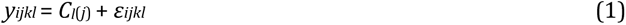

The residuals (*ε_ijkl_*) were then used as features (*y_ijk_*) for further analysis. For the four side-view images of a fruit, the feature values were averaged (SAv) or their standard deviation (SSD) was calculated. Mean values (Av) and standard deviations (SD) were calculated across up to five fruits harvested from each tree (replicate of a genotype). For each feature, the feature extraction finally resulted in one value per tree and year, i.e., one value for each combination of genotype, replicate and year.

### 2.4 Feature extraction for skin color

From the color histogram of each image with its background removed, features for color were extracted (Table 1). Initially, a histogram for either hue or saturation color space was obtained for each genotype by averaging all images from side view taken for the genotype. The average histograms per genotype were hierarchically clustered and three clusters of genotypes were defined for each color space. Average histogram was plotted for each of the three clusters of genotypes and the two color spaces, and the maximum values of the histograms (peaks) were identified (Figure 1C). Next, the feature values at the peaks (peak +/− 0.05 of the color space) were extracted for each fruit image separately. They were categorized into red, brown, and yellow hue as well as low, medium, and high saturation. The green colored apple fruits were not sufficiently represented to form an own peak.

The features were corrected for the camera effect using model 1. For the four side-view images of a fruit, the feature values were averaged (SAv) or their standard deviation (SSD) was calculated. Then, the mean values (Av) and standard deviations (SD) across up to five fruits harvested from each tree (replicate of a genotype) were calculated. Color features were always based on the full fruit (AB). For each feature, the feature extraction finally resulted in one value for each combination of genotype, replicate and year.

### 2.5 Statistical analysis

For the extracted features, outlier data points were removed when they were 15 times larger or lower than the size of the interquartile range. For every feature, adjusted means of each genotype (*G_i_*) were modeled for each extracted feature value per tree (*y_ijk_*) as the following:

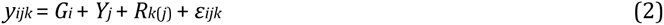

where *Y_j_* was a fixed effect for the year, *R_k_*_(*j*)_ was a fixed effect for the replicated genotype nested in year and *ε_ijk_* was the error term. For these adjusted features, clonal mean heritability (*H*^2^) was calculated as the genotypic variance (*σ_g_*) divided by the sum of genotype and the error variance (*σ_ε_*) adjusted for the average number of replicates over both years (*n̅*):

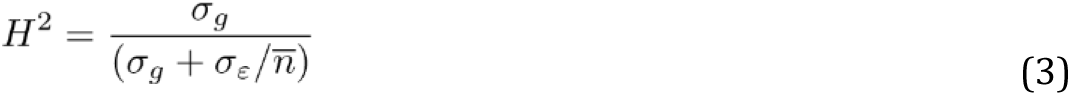

To reduce dimensions of the set of adjusted features, principal component analysis (PCA) was carried out. The PCA was performed separately for (*i*) the fruit shape and size features as well as for (*ii*) fruit skin color features using the R package FactoMineR (Lê et al. 2008). The adjusted features showing an *H*^2^ lower than 0.4 were removed prior to PCA to focus the analysis on highly heritable features. From the highly heritable features with an *H*^2^ higher than 0.6, a set of uncorrelated features with a Pearson correlation lower than 0.75 between each other was chosen using the *findCorrelation* command of the R package caret (Kuhn 2008). The uncorrelated features and supplementary traits measured by the conventional methods (sorting machine and visual scoring) were correlated to the principal components (PCs) and overlayed on a biplot.

The GWAS was carried out using the Bayesian-information and Linkage-disequilibrium Iteratively Nested Keyway (BLINK) algorithm (Huang et al. 2018). The GWAS was performed for the first nine PCs (PC1 to PC9) for fruit shape and size, first four PCs for fruit skin color, and for the set of uncorrelated features for fruit shape, size, and color. For GWAS, the imputed SNP matrix for the apple REFPOP genotypes was obtained from Jung et al. (2020). The position of the SNP markers was determined according to the reference genome GDDH13 (v1.1) (Daccord et al. 2017). The first three PCs of the SNP matrix were used to correct for population structure in GWAS, mainly to adjust for the genetic differences between progeny and accessions. The *p-*values resulting from GWAS were Bonferroni-corrected with the significance threshold set to the 5% level, or (where indicated) to the 1% level for in-depth analysis.

To assess the additive effect of all SNPs associated with one shape and size PC, or all associations with PCs and uncorrelated traits, the SNPs ordered by a decreasing −*log*_10_(*p*) value were included in a linear regression model, and the regression sum of squares explained by each SNP was estimated. All SNPs associated with a shape and size PC were collapsed into haplotypes to assess their combined effect on the PC. The haplotypes were further classified into hierarchical clusters. Hierarchical cluster analysis was performed based on the dissimilarities of the haplotypes using the *hclust* command from the R package stats with the method set as “complete”. To assess the combined effect of physically linked SNPs associated with PCs and uncorrelated traits for shape and size, haplotypes were constructed by collapsing all SNP values within 0.1 Mbp distance (further denoted as haplotypes based on groups of physically linked SNPs). For each haplotype cluster and every haplotype based on groups of physically linked SNPs, the average side-view fruit contour was plotted in absolute and relative values (mean=0 and standard deviation=1).

Major effects of the SNPs associated with fruit shape and size PCs and uncorrelated features were assessed from the visualization of the average side-view fruit contour for each allele dosage (0, 1 or 2 alternative alleles). The associated SNPs were classified as affecting fruit size (size), leading to lengthened fruits (conical shape) or very long fruits (cylindrical shape). The basic fruit shape for all classes was round, with the opposite effect of one of the alleles in a SNP resulting in conical or cylindrical shape (designated as favorable allele). Following these principles, the major effects were also visually assessed for the average side-view fruit contours for haplotype clusters and haplotypes based on groups of physically linked SNPs associated with PCs and uncorrelated traits for fruit shape and size. For the fruit skin color PCs and uncorrelated features, the major effects of allele dosage (0, 1 or 2 alternative alleles) were visually assessed from feature distributions along the hue and saturation spectra. Allele associated with a peak in the red part of the hue spectrum was assumed as the baseline, with the opposite allele classified as being associated with light green or yellow hue. Alleles associated with light green or yellow hue were designated favorable.

All statistical analyses and data formatting in this article were performed with R (R Core Team 2014).

### 2.6 Comparison with previously published QTL

Earlier reports on QTL mapping and GWAS in apple that were extensively reviewed and reported by Jung et al. (2022) were taken for comparison with the associations reported in this study. From the review of 41 studies, the trait group named fruit size was considered for the comparison with the SNPs associated with fruit shape and size PCs and uncorrelated features, and the trait groups fruit ground color and over color were considered for the comparison with the associations with the fruit color PCs and uncorrelated features. Jung et al. (2022) visually assigned the positions of the reviewed associations within respective chromosomes to the three chromosome segments, i.e., top, center and bottom. To compare chromosome-segment combinations found for loci in our and the earlier studies, the SNPs on each of the 17 chromosomes in our dataset were categorized into three equally sized chromosome segments, while the length of each chromosome was defined by its last marker. Following these principles, QTL obtained for the ratio between fruit length and diameter found by Sun et al. (2012) and Chang et al. (2014), and results of the study of fruit size by Liao et al. (2021), which were not included in the review of Jung et al. (2022), were added to the comparison with shape and size PCs and uncorrelated features.

Associations with fruit shape and size as well as color traits reported recently by Duan et al. (2017), Minamikawa et al. (2021), Liao et al. (2021), Jung et al. (2022) and Dujak et al. (in prep.) were compared with the results obtained in our study. Position of DNA sequences containing selective sweeps associated with fruit size published by Duan et al. (2017) were estimated by the alignment of the sequences to the reference genome of the ‘Golden Delicious’ doubled-haploid line GDDH13 (v1.1) (Daccord et al. 2017) using the Genome Database for Rosaceae (https://www.rosaceae.org). Previously published molecular markers and significant SNPs identified in the current study were considered as co-localized when they were less than 0.1 Mbp distant from each other.

## 3 Results

### 3.1 Shape and size features

A total of 1,248 adjusted features were filtered for heritability values above 0.4, which resulted in 573 highly heritable features for fruit shape and size. Selection for uncorrelated features for shape and size resulted in 16 features showing various distributions (Figure S1A). Among the uncorrelated features, one was derived from images of the top view of fruits, ten accounted for standard deviation within the individual fruit shape or size of the different side views and five represented the average of the fruit shape or size measures from the side views. A comparison of principal components (PCs) derived from the 573 highly heritable features with the 16 uncorrelated features and the supplementary traits measured by the sorting machine showed that the PC1 was positively correlated with the sorting machine traits fruit length, volume, and single fruit weight (Figure 2A). The PC2 was strongly correlated with the symmetry features (e.g., Av_SSD_Pol_SD_R_Sym.rel) and the shape heterogeneity features defined as standard deviation between measures of different fruit parts derived from polar data of the fruit side views (e.g., Av_SSD_Pol_SD_B.LR_Max). For the highly heritable features, the PC1 and PC2 explained 48.2% and 11.9% of the variance, respectively. The PC3 was strongly positively correlated with the feature Av_Top_Pol_Av_A.LR_RatM, which is a parameter describing fruit roundness from top view and negatively correlated with the sorting machine traits related to fruit size (Figure 2B). The PC4 was strongly correlated with a symmetry feature (Av_Sav_Pol_SD_B_Sym.rel) and the feature Av_SSD_Pol_Rat.Av_AB.LR_SD describing the symmetry and homogeneity of the fruit. The PC3 and PC4 explained 8.9 and 4% of the variance, respectively. The PC5 to 8 explained less than 3% of the variance each (Figure S2).

**Figure 2:**
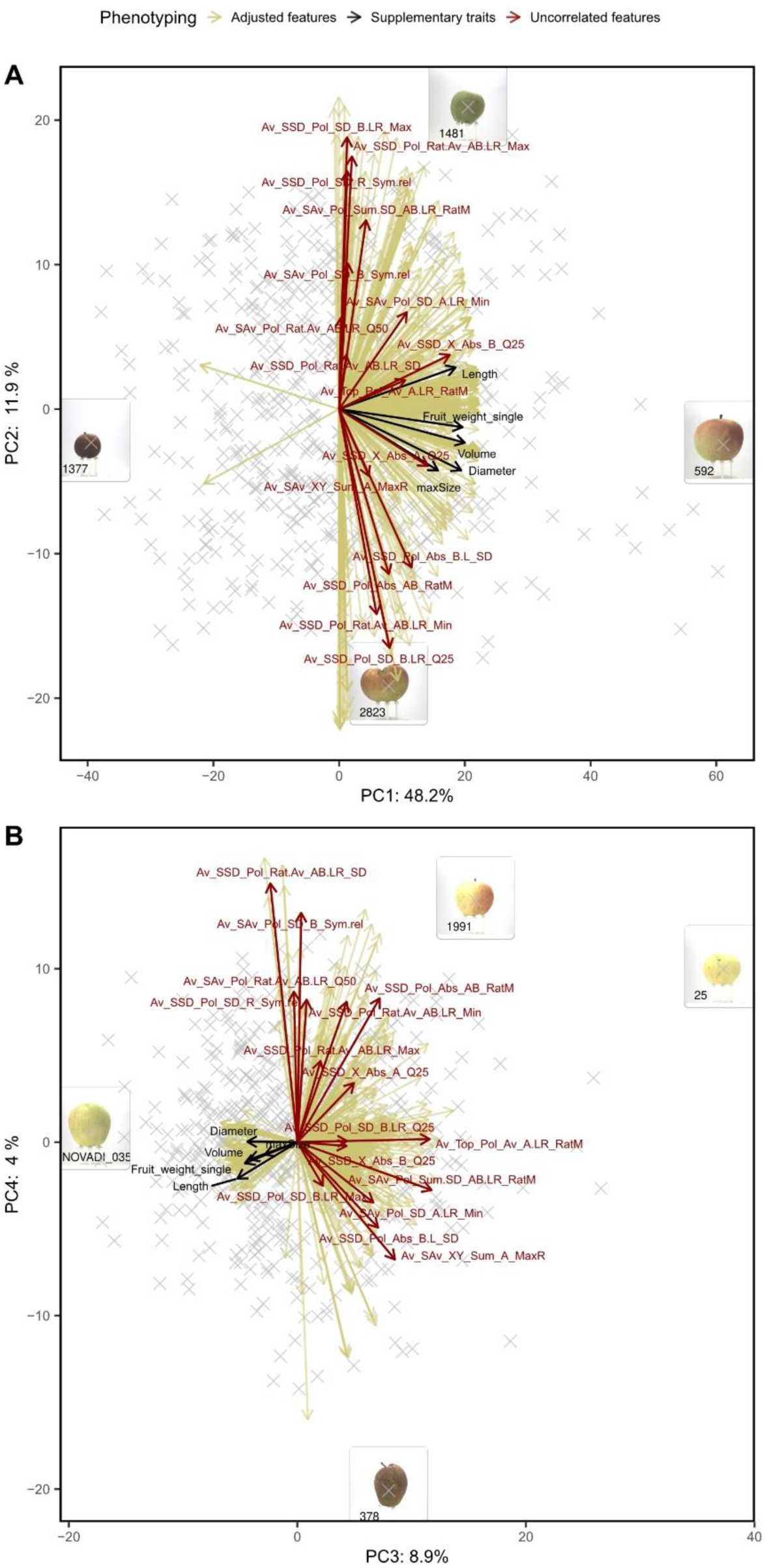
Principal component (PC) analysis of highly heritable features for fruit shape and size. Uncorrelated (r < 0.75) fruit shape and size features (red arrows) and supplementary features measured using the sorting machine (black arrows) were overlayed in a biplot. Images of extreme genotypes of the apple REFPOP were shown in boxes and labeled with their respective genotype codes. A: PC1 and PC2, B: PC3 and PC4.

### 3.2 Skin color features

Filtering 20 adjusted features for heritability (*H*^2^ *>* 0.4) resulted in 17 highly heritable fruit skin color features. From the highly heritable features, eight uncorrelated features were identified, and they represented all color feature groups (chosen were the low, medium, and high saturation, as well as red, brown, and yellow hue features). The uncorrelated color features showed various distributions (Figure S1B). In a PCA of the highly heritable features, the features were well represented by PC1 and PC 2 that accounted for 54.8% of the variance in the data (Figure 3A). When PCs derived from the highly heritable features were compared with the uncorrelated features and the supplementary traits measured by the sorting machine or scored visually, the biplot showed that PC1 was strongly positively correlated with the green color assessed by the sorting machine and the uncorrelated features for yellow hue (Figure 3A). Furthermore, the PC1 was strongly negatively correlated with the red color scored visually and by the sorting machine as well as the uncorrelated feature representing red hue. The PC2 was strongly positively correlated with the uncorrelated feature for low saturation and negatively correlated with the visually scored ground color (discriminating between green and yellow ground color). The PC3 was strongly correlated with the uncorrelated color saturation features (i.e., Av_SAv_Sat_Abs_AB_Medium and Av_SAv_Sat_Abs_AB_High) (Figure 3B). The PC4 was correlated with the sorting machine trait green and the uncorrelated features related to low saturation and yellow hue (e.g., Av_SSD_Hue_Abs_AB_Yellow). The uncorrelated features for brown hue (Av_SSD_Hue_Abs_AB_Brown and Av_SAv_Hue_Abs_AB_Brown) were well represented by the plane of PC3 and PC4 as shown by the long arrows, but these features did not strongly correlate with either of the PCs.

**Figure 3:**
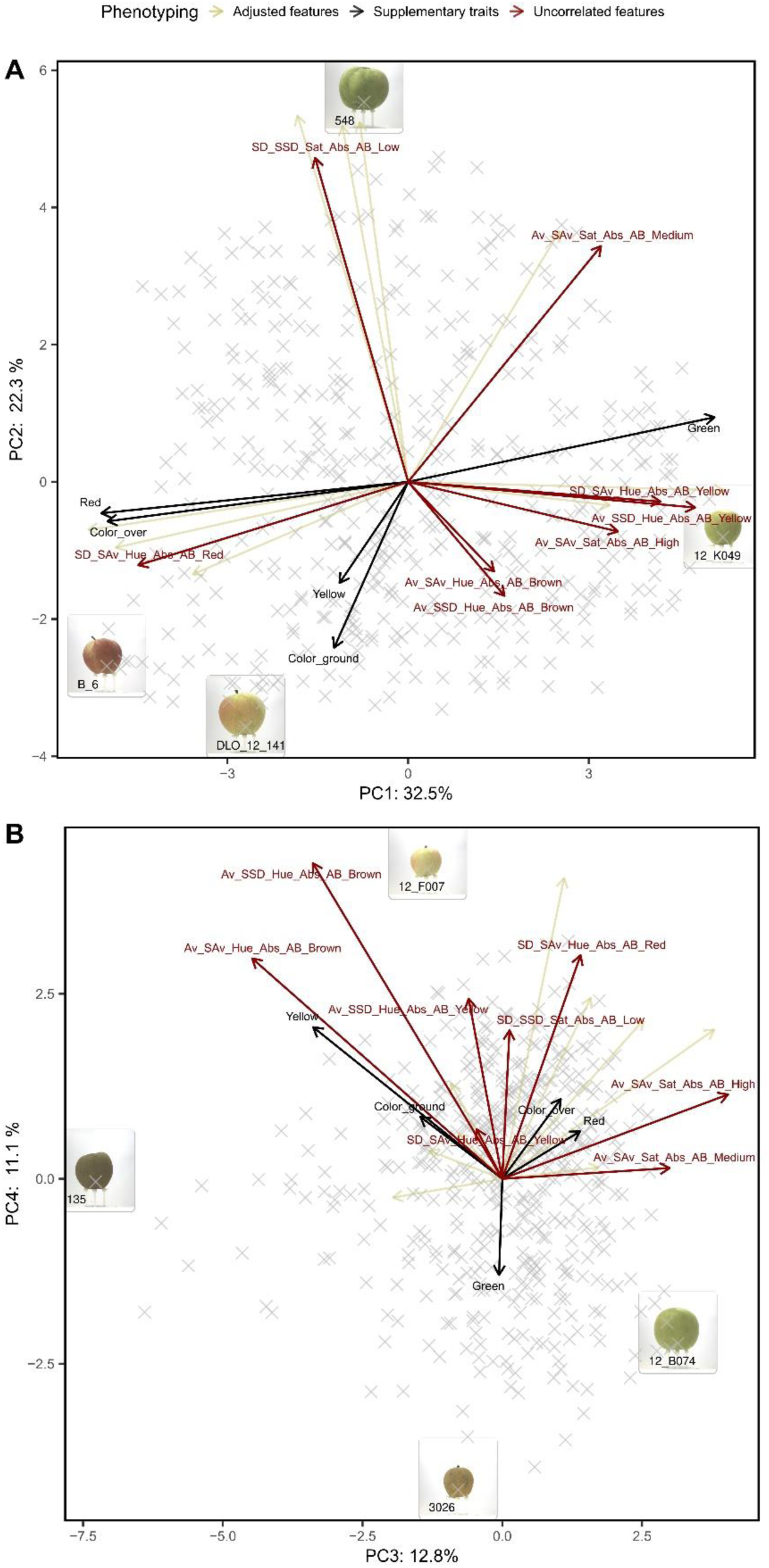
Principal component (PC) analysis of highly heritable features for color. Uncorrelated (r < 0.75) fruit color features (red arrows) and supplementary features measured visually or using the sorting machine (black arrows) were overlayed in a biplot. Images of extreme genotypes of the apple REFPOP were shown in boxes and labeled with their respective genotype codes. A: PC1 and PC2, B: PC3 and PC4.

### 3.3 Genome-wide association studies for shape and size

In total, 21 SNPs were significantly associated with PC1 to PC5 (Figure 4, Figure 5A, Table S1). The plot of expected and observed −*log*_10_(*p*) values showed no apparent *p-*value inflation and therefore a good control of population structure (Figure 5B). Out of the SNPs significantly associated with PC1 to PC5, 13 SNPs were observed at 1% significance level and showed mainly additive effects of the allele dosage (0, 1 or 2 alternative alleles) on the phenotype (Figure 5C). Different proportions of phenotypic variance were explained by the SNPs associated with PCs (Table 2). Seven SNPs associated with PC1 explained together 35.7% of the phenotypic variance with additive effects for each SNP explaining between 2.8% and 9.2% of the variance. The five SNPs significantly associated with PC2 explained 21.1% of the variance with additive effects for each SNP explaining between 0.3% and 6.4% of the variance. The SNPs associated with PC3 to PC5 explained 18.5%, 18.4% and 3.9% of the phenotypic variance, respectively. The PC6 to PC9 did not show any significant associations with SNP markers (Figure S3).

**Figure 4:**
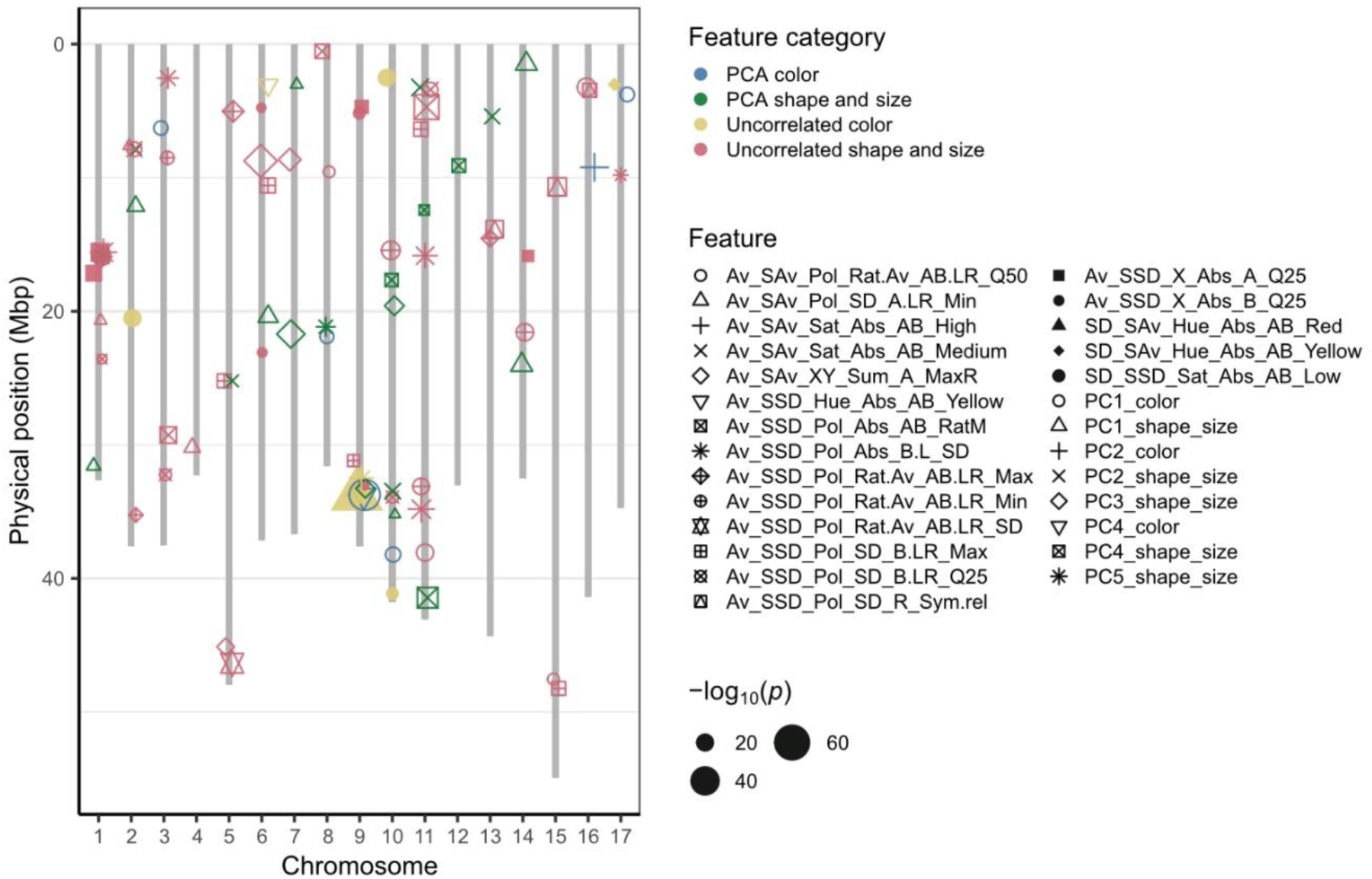
Distribution of the significant associations of markers with PCs and uncorrelated traits on the apple genome. The colors of the points represent feature categories, shapes of points stand for the individual features and the point size represents the −*log*_10_(*p*) value.

**Figure 5:**
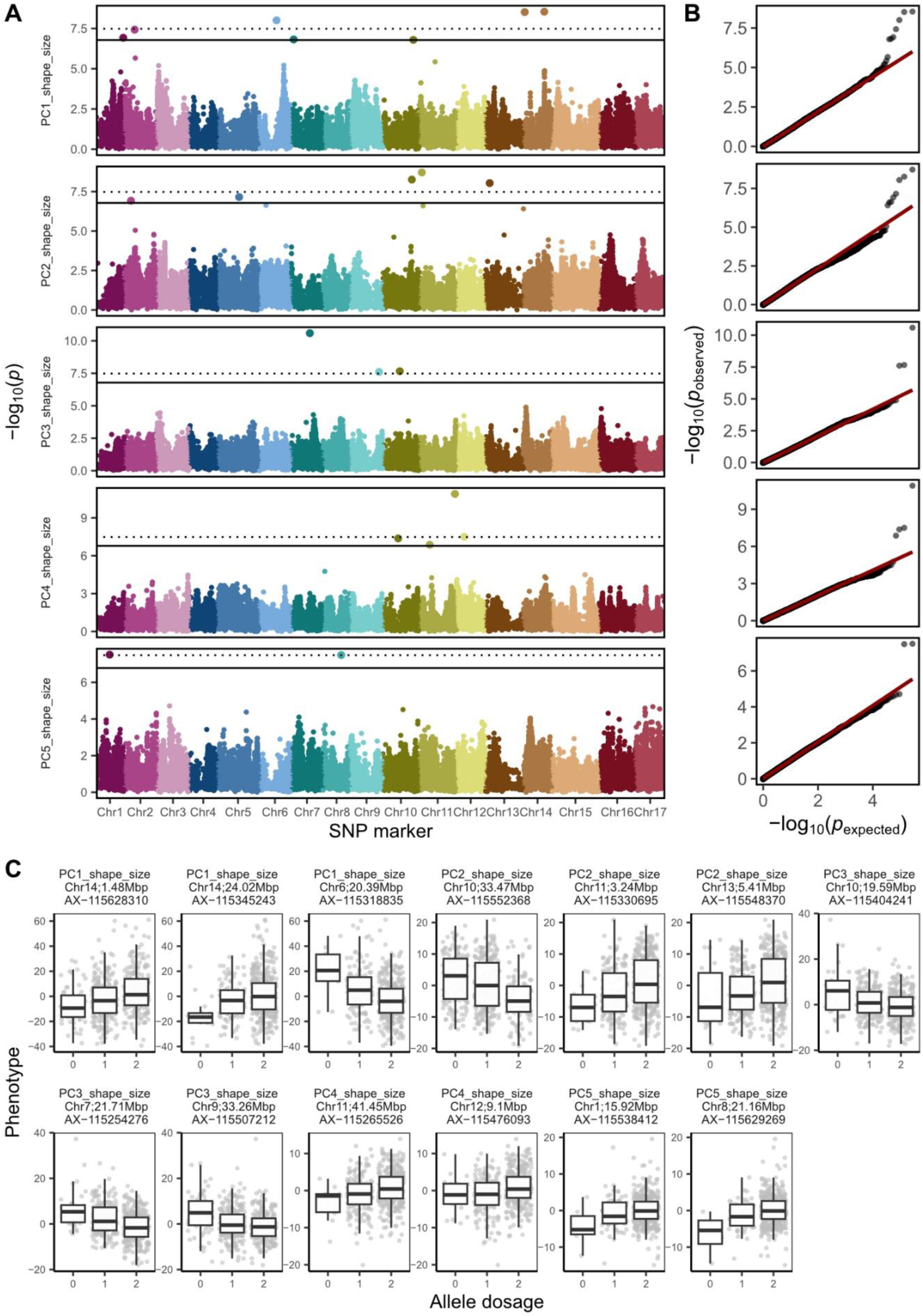
Results of the genome-wide association studies for the first five principal components (PCs) build from the highly heritable fruit shape and size features. A: Manhattan plots with the −*log*_10_(*p*) value for every SNP marker and PC. The SNP markers are displayed according to their physical position on the genome and chromosome (Chr). The Bonferroni-corrected significance level is indicated at 5% and 1% by the solid and dotted line, respectively. B: The expected and observed *p*-values for SNP markers and PCs. The 1:1 line for non-significant *p-*values is shown in red. C: The boxplots for allele dosage effect (0, 1 or 2 alternative alleles) for SNPs significantly associated with PCs at 1% significance level.

**Table 2:** Significant marker associations with principal component (PC) 1 to 5 for shape and size. Marker name, chromosome (Chr), position (Pos) in base pairs, *p*-value, and the respective sum of squares and explained variance for each marker and the residual are shown.

| PC | Marker | Chr | Pos (bp) | *p*-value | Sum of squares | Explained variance (%) |
| --- | --- | --- | --- | --- | --- | --- |
| PC1 | AX-115460130 | 7 | 3024174 | 1.52E-07 | 3934.81 | 2.8 |
| PC1 | AX-115345243 | 14 | 24020772 | 2.87E-09 | 4295.028 | 3.1 |
| PC1 | AX-115227580 | 10 | 35210847 | 1.62E-07 | 5824.678 | 4.2 |
| PC1 | AX-115534322 | 2 | 12130252 | 3.75E-08 | 6050.312 | 4.3 |
| PC1 | AX-115628310 | 14 | 1478099 | 3.03E-09 | 8193.44 | 5.9 |
| PC1 | AX-115318835 | 6 | 20393412 | 9.86E-09 | 8516.722 | 6.1 |
| PC1 | AX-115380250 | 1 | 31575453 | 1.19E-07 | 12866.27 | 9.2 |
| PC1 | Residual |  |  |  | 89512.3 | 64.3 |
| PC2 | AX-115490525 | 2 | 7904059 | 1.20E-07 | 112.5119 | 0.3 |
| PC2 | AX-115548370 | 13 | 5411038 | 9.11E-09 | 915.1389 | 2.8 |
| PC2 | AX-115429431 | 5 | 25209696 | 2.13E-08 | 1888.73 | 5.8 |
| PC2 | AX-115330695 | 11 | 3241944 | 1.93E-09 | 1921.867 | 5.9 |
| PC2 | AX-115552368 | 10 | 33469975 | 5.50E-09 | 2090.653 | 6.4 |
| PC2 | Residual |  |  |  | 25874.43 | 78.9 |
| PC3 | AX-115404241 | 10 | 19586733 | 2.20E-08 | 767.3217 | 3 |
| PC3 | AX-115507212 | 9 | 33255077 | 2.48E-08 | 948.2588 | 3.8 |
| PC3 | AX-115254276 | 7 | 21705520 | 2.58E-11 | 2957.387 | 11.7 |
| PC3 | Residual |  |  |  | 20563.65 | 81.5 |
| PC4 | AX-115659654 | 11 | 12416234 | 1.35E-07 | 434.6513 | 3.8 |
| PC4 | AX-115476093 | 12 | 9098303 | 3.16E-08 | 441.0631 | 3.9 |
| PC4 | AX-115419379 | 10 | 17661953 | 4.24E-08 | 591.8564 | 5.2 |
| PC4 | AX-115265526 | 11 | 41448132 | 1.26E-11 | 633.1262 | 5.6 |
| PC4 | Residual |  |  |  | 9294.302 | 81.6 |
| PC5 | AX-115538412 | 1 | 15918343 | 3.09E-08 | 124.1923 | 1.5 |
| PC5 | AX-115629269 | 8 | 21161287 | 3.25E-08 | 197.7971 | 2.4 |
| PC5 | Residual |  |  |  | 7864.159 | 96.1 |

Visual assessment of the effect of allele dosage on the side-view fruit contour for each associated SNP and PC showed major effects of individual SNPs on PCs (Figure S4, Table S1). SNPs significantly associated with PC1 affected fruit size but showed only a minor effect on fruit shape. The SNPs associated with PC2 and PC3 showed a major effect on fruit shape, with different allele dosage resulting in round or lengthened (conical) fruit shapes. Two SNPs associated with PC4 affected fruit size, one SNP resulted in contour of round or conical fruit shape and one SNP (AX−115419379) was associated with round or very long (cylindrical) fruit contour. The associations with PC5 influenced both the fruit size (AX−115629269) and conical shape (AX−115538412). For haplotypes assembled from SNPs associated with each of the PCs, additive effects of haplotypes on PCs were found, and visual differences in shape and/or size were observed for images of genotypes from the extremes of the PC distribution (Figure S5-S8).

When haplotypes built from the 21 SNPs associated with PC1 to PC5 were hierarchically clustered, a pronounced effect of the individual clusters for each PC on fruit shape and size was observed (Figure 6, Table S5). Although all PCs affected both fruit shape and size, the effect on fruits size of the haplotype clusters for PC1 and especially for PC5 was particularly strong. The haplotype clusters for PC2, PC3 and PC4 were mostly responsible for round or conical fruit shape. The haplotype cluster ‘2’ suggested an effect of PC4 on cylindrical fruit shape.

**Figure 6:**
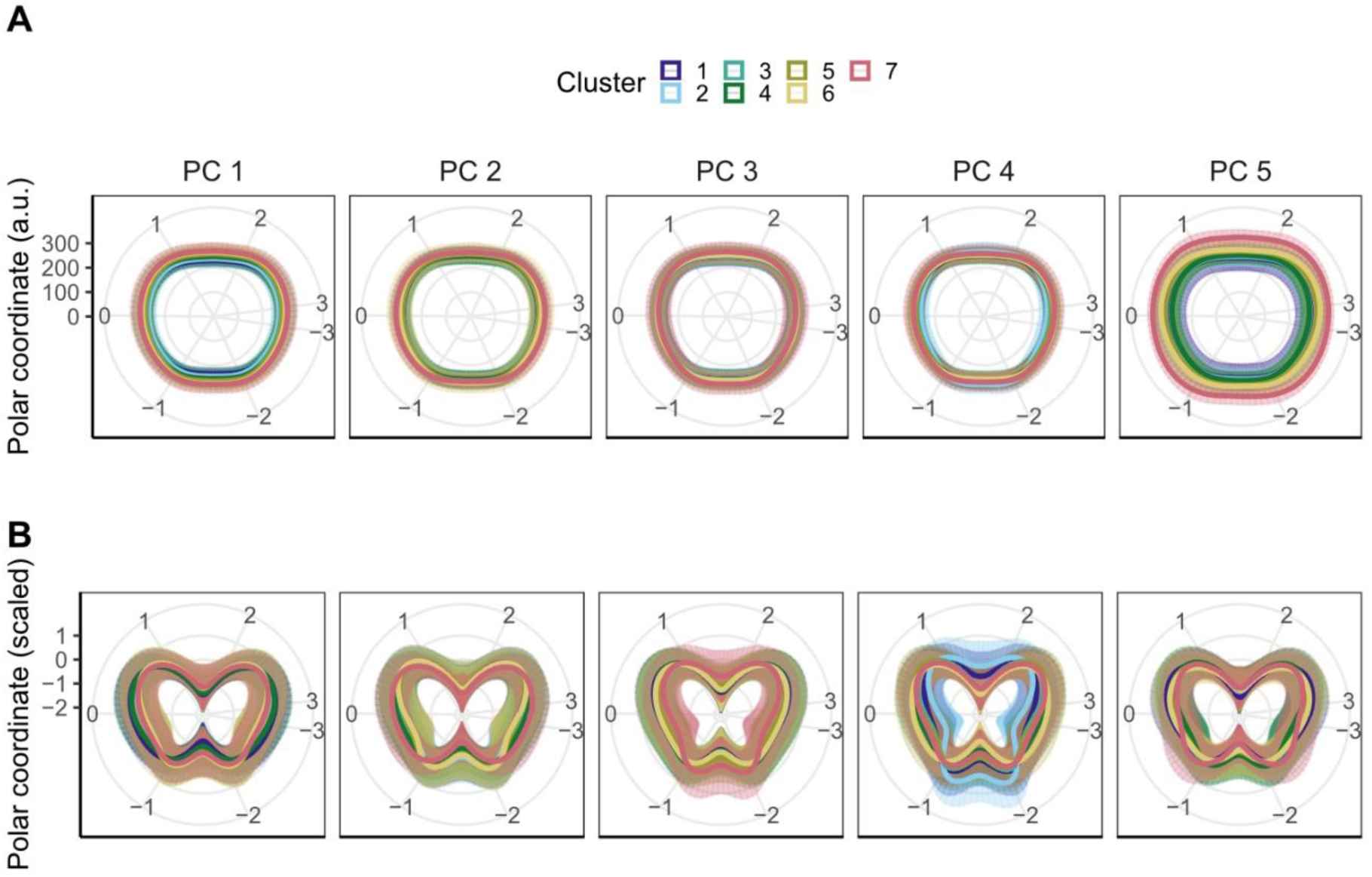
Effect of haplotype clusters based on the SNPs associated with the principal component (PC) 1, 2, 3, 4 and 5 for shape and size. The effect of each cluster on the averaged fruit contour is shown in A: absolute values and B: relative values. The relative values were scaled (mean=0 and standard deviation=1) for every individual apple fruit with the resulting minimum value being the center of the graph, i.e., the visualized diameter size is arbitrary, and the shape effect is enhanced. The error bars show the standard deviation of all genotypes included in the haplotype cluster (visible as semitransparent area around the fruit contour). The Bonferroni-corrected significance level of the considered associations was 5%.

For 15 out of 16 uncorrelated fruit shape and size features, 48 associations of markers with features were identified, and they affected fruits conical shape, cylindrical shape, or size (Figure 4, Figure S10, Table S1). All associations with the uncorrelated features referred to the side view of the cameras and contours. No significant associations with the uncorrelated feature for the top camera view were found, and therefore the visual inspection of the top-view contours was omitted. For haplotypes assembled from SNPs associated with the uncorrelated symmetry feature Av_SSD_Pol_SD_R_Sym.rel, additive effects of haplotypes on the feature were found, and visual differences in fruit symmetry were observed for images of genotypes from the extremes of the feature distribution (Figure S9).

Ten of the SNPs associated with PCs or uncorrelated features occurred physically close to each other (within 100,000 bp between pairs of SNPs) in four genomic regions with two or three SNPs located in each region. After these closely located SNPs were grouped into haplotypes for each of the genomic regions (Figure 7), the three SNPs on Chr 1 and 2 resulted in nine haplotypes each, the two SNPs on Chr 11 and 16 in seven and eight haplotypes, respectively. Although different haplotypes of the four genomic regions showed an effect on both fruit shape and size, the differences in fruit size between haplotypes of the genomic region on Chr 1 at 15.9 Mbp were particularly strong. In this region, the haplotype ‘220’ was associated with fruit contour of a notably small size. In the genomic region on Chr 2 at 7.9 Mbp, the haplotypes ‘220’, ‘210’ and ‘212’ resulted in decidedly conical shapes. The haplotype ‘10’ on Chr 11 at 3.3 Mbp showed an effect towards large fruit size. The haplotypes on Chr 16 at 3.4 Mbp showed no pronounced effect on fruit shape or size.

**Figure 7:**
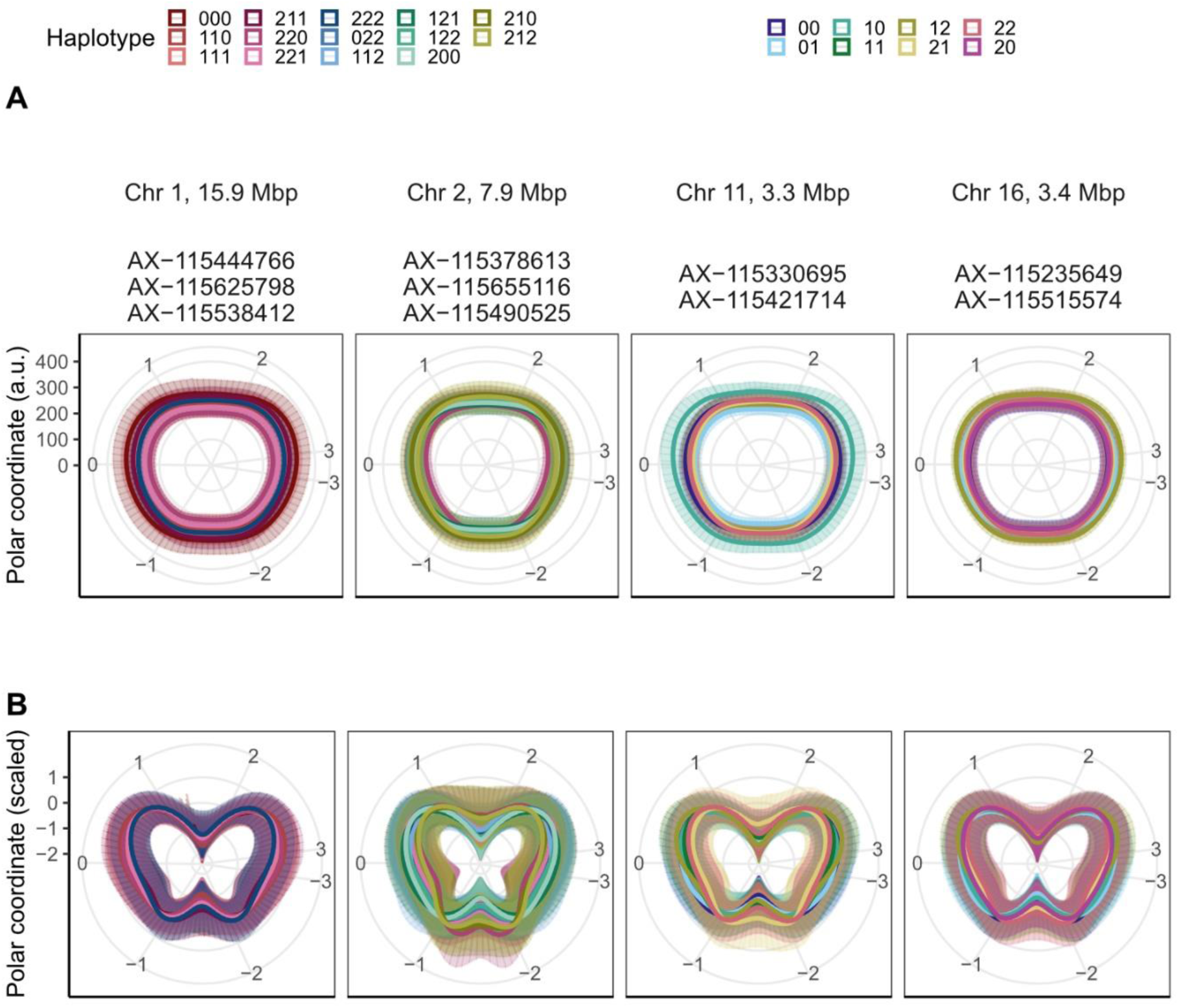
Effect of haplotypes based on groups of physically linked SNPs for shape and size found in four genomic regions. The effect of each haplotype on the averaged fruit contour is shown in A: absolute values and B: relative values. The relative values were scaled (mean=0 and standard deviation=1) for every individual apple fruit with the resulting minimum value being the center of the graph, i.e., the visualized diameter size is arbitrary, and the shape effect is enhanced. The error bars show the standard deviation of all genotypes included in the haplotype cluster (visible as semitransparent area around the fruit contour). The Bonferroni-corrected significance level of the considered associations was 5%.

### 3.4 Genome-wide association studies for skin color

In total, 16 SNPs were significantly associated with three PCs and six out of the eight uncorrelated features for the apple fruit skin color (Table S1). Seven SNPs were significantly associated with PC1, PC2 and PC4, while no significant association was observed for PC3 (Figure 8A). No *p-*value inflation was visible in the Q-Q plot (Figure 8B). All seven SNPs were identified above the 1% significance level and showed mainly additive effects on the color PCs (Figure 8C). Upon visual inspection of major effects of allele dosage (0, 1 or 2 alternative alleles) that were assessed from feature distributions along the hue and saturation spectra (Figure 9, Figure S12), alleles associated with red and light green hue or red and yellow hue were identified (Table S1). Following this principle, five SNPs associated with PC1, PC4, medium saturation, high saturation, red and yellow hue in the genomic region on Chr 9 at 32.5–33.8 Mbp showed two peaks for red hue and one prominent peak for light green hue (Figure 9, Figure S12). For saturation, all three allele dosages of the SNPs in the genomic region on Chr 9 showed distributions resembling bell curves of similar mean with an increased kurtosis for the allele associated with light green hue (Figure 9, Figure S12). Additionally, similarities between distributions along the hue and saturation spectra were found for three SNPs associated with PC2 on Chr 16 at 9.23 Mbp and a low saturation uncorrelated feature (SD_SSD_Sat_Abs_AB_Low) on Chr 2 at 20.51 Mbp and Chr 10 at 2.51 Mbp (Figure 9, Figure S12). For these three SNPs, a peak was found in the yellow part of the hue spectrum and shifts of the distributions for the different allele dosages on the saturation spectrum towards low and high saturation were observed. The remaining seven SNPs associated with the color features showed only minor (undefinable) effects on the hue and saturation spectra (Figure 9, Figure S12). For haplotypes assembled from SNPs significantly associated with the uncorrelated color feature SD_SAv_Hue_Abs_AB_Yellow, visual differences in fruit color were observed for images of genotypes from the extremes of the feature distribution (Figure S11).

**Figure 8:**
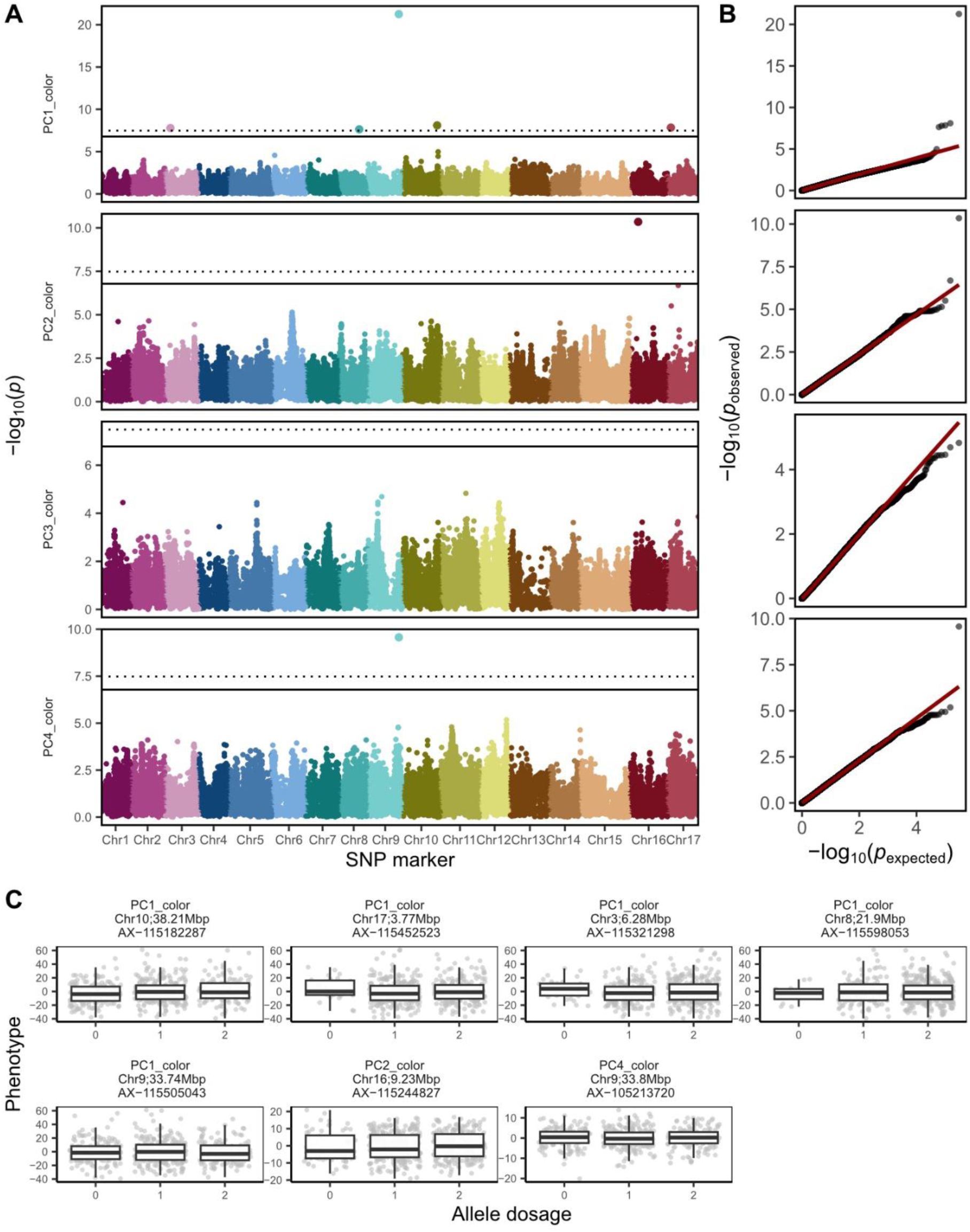
Results of the genome-wide association studies for the first four principal components (PCs) build from the highly heritable fruit color features. A: Manhattan plots with the −*log*_10_(*p*) value for every SNP marker and PC. The SNP markers are displayed according to their physical position on the genome and chromosome (Chr). The Bonferroni-corrected significance level is indicated at 5% and 1% by the solid and dotted line, respectively. B: The expected and observed *p-*values for SNP markers and PCs. The 1:1 line for non-significant *p-*values is shown in red. C: The boxplots for allele dosage effect (0, 1 or 2 alternative alleles) for SNPs significantly associated with PCs at 1% significance level.

**Figure 9:**
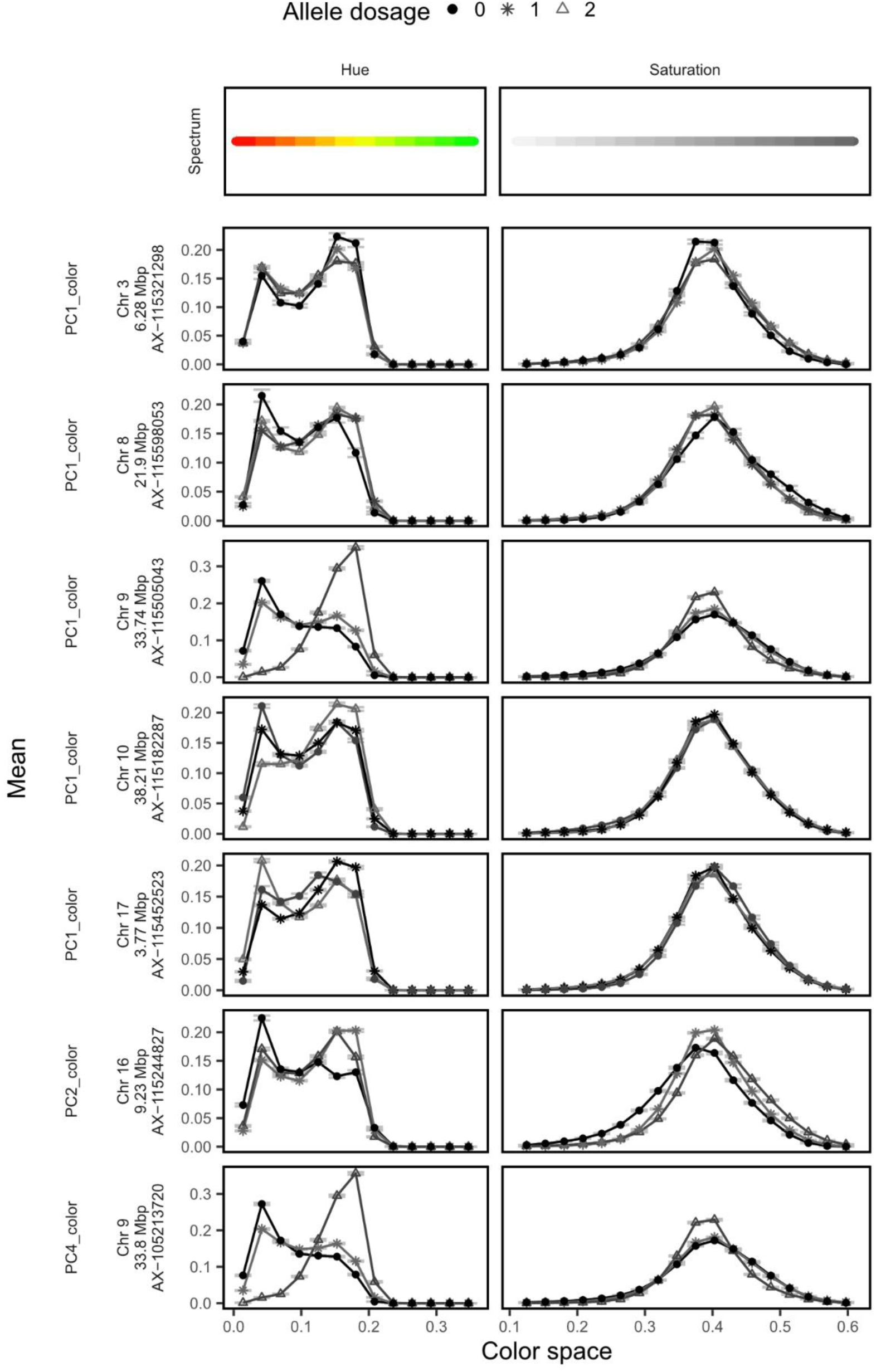
Allele dosage effect (0, 1 or 2 alternative alleles) of the SNPs significantly associated with the principal component (PC) 1, 2 and 4 on the hue and saturation color space. The error bars show the standard deviation of all genotypes per allele dosage. The Bonferroni-corrected significance level of the considered associations was 5%.

### 3.5 New and previously published associations

Out of 69 associations of markers with PCs and uncorrelated features for fruit shape and size and 16 associations of markers with PCs and uncorrelated features for fruit skin color that were identified in this study, 19 SNPs co-localized in genomic regions (genetic distance *<* 0.1 Mbp) with 35 associations that have been reported in previous publications (Table S2). Two markers (AX-105213720, AX-115452523) identified in this study were identical as reported by Jung et al. (2022) and associated with green color and red over color. Additionally, 119 potential overlaps (corresponding to 48 SNP markers) were found between our and the previously reported loci for fruit shape and size (Table S3). No overlapping positions with QTL reported in previous studies were found for 19 and three SNPs associated with shape/size and color features, respectively (Table S4).

## 4 Discussion

### 4.1 Fruit shape and size

For over a decade, numerous studies have discovered QTL associated with apple fruit quality traits such as fruit shape and size (Devoghalaere et al. 2012; Kenis et al. 2008; Sun et al. 2012). The extensive collection of published fruit quality loci was recently reviewed by Jung et al. (2022). Making use of this review, we found several groups of three to four associated SNPs from our study that matched chromosome segments found in former studies and potentially pinpointed loci of broader interest (Table S3). Such groups of SNPs associated with the uncorrelated features and PCs for fruit shape and size were located on Chr 9 at 31.2–33.3 Mbp, Chr 10 at 15.4-19.6 Mbp and Chr 11 at 3.2-6.4 Mbp, and various QTL for fruit size were previously reported in the proximity of these loci (Chang et al. 2014; Kenis et al. 2008; Liao et al. 2021; Liu et al. 2016; Minamikawa et al. 2021; Sun et al. 2012). Not only the traits characterizing fruit size, but also fruit shape described as ratio between fruit length and diameter has been previously associated with QTL that colocalized with the associated genomic regions we identified on Chr 9 at 31.2–33.3 Mbp, Chr 10 at 15.4-19.6 Mbp and Chr 10 at 33.5-35.2 Mbp (Chang et al. 2014; Sun et al. 2012). The numerous overlaps between the known and the presently reported loci proved that FruitPhenoBox is a suitable tool for high-throughput phenotyping of fruit shape and size, and that the obtained features can be successfully linked with genomic information using GWAS.

Out of all associations reported here for the uncorrelated features and PCs for fruit shape and size, 19 SNPs located across seven different chromosomes did not colocalize with QTL from the literature reviewed for fruit shape and size traits (Table S4). Among the novel loci, a group of five SNPs on Chr 1 at 15.6–20.7 Mbp was associated with fruit size (four SNP markers) and conical shape (one SNP marker). Three of these SNPs occurred physically close to each other (within 100,000 bp), and the haplotypes built with them showed a pronounced effect on fruit size (Figure 7). The different haplotypes provide with novel information about the genetic architecture of fruit size, and they could be used for marker-assisted selection for fruits of different sizes. To our knowledge and despite the numerous QTL known for size-related traits in apple (Costa 2015; Devoghalaere et al. 2012; Duan et al. 2017; Kenis et al. 2008), marker-assisted selection for fruit size has yet to be applied in apple. Markers selecting for fruit size may be especially helpful when breeding new varieties bearing resistances to various pests and diseases originating from small-fruit wild apple accessions (e.g., *M. floribunda* 821, the ornamental ‘Evereste’ or *Malus* × *robusta* 5). Selecting for haplotypes associated with larger fruit size using marker-assisted selection could improve efficiency of breeding such resistant varieties.

For fruit shape, the novel associations with conical and cylindrical fruit shapes can assist in breeding for or against exotic fruit shapes in the future and offer a possibility to explore fruit shape beyond the traditionally studied fruit length/diameter ratio (Chang et al. 2014; Sun et al. 2012). Because of their association with round and prolonged shapes that were visible from the average fruit contours, the individual SNPs and haplotypes built of SNPs of the newly discovered genomic region on Chr 2 at 7.6–7.9 Mbp can offer the opportunity for breeders to address round or conical fruit shapes (Figure 7).

Despite the distinct effect of the newly discovered group of loci on Chr 2 at 7.6–7.9 Mbp on fruit shape, a minor effect of the haplotypes on fruit size was observed from the averaged fruit contours (Figure 7). Similar combination of effects on both fruit shape and size was visible for the novel group of loci on Chr 1 at 15.6–20.7 Mbp that was primarily associated with fruit size, as well as for the groups of loci on Chr 11 and 16 (Figure 7). Loci associated simultaneously with shape and size have been shown in other crops such as cucurbits, tomatoes, or pear (Pan et al. 2020; Sierra-Orozco et al. 2021; Zhang et al. 2013). Although it was proposed earlier that size and shape are under independent genetic control in apple (Chang et al. 2014), the simultaneous effects of the loci described here on both fruit shape and size may suggest a biological interdependence between the shape and size of apple fruits.

### 4.2 Fruit skin color

Many studies have explored the genetic mechanisms behind coloring of apple fruit, resulting in reports of QTL mostly located on Chr 9 (Bianco et al. 2014; Chagné et al. 2016; Duan et al. 2017; Moriya et al. 2017). For the associations with the color PCs and uncorrelated features found here (Table S1), numerous overlaps in the bottom segment of Chr 9 at 32.5–33.8 Mbp were found between our associations and the associations found in the review of Jung et al. (2022) (Table S3). These associations are all likely related to the degree of red over color, and they confirmed the role of the locus on Chr 9 in genetic regulation of red skin color in apples.

Although the locus on Chr 9 is known to explain the majority of the phenotypic variance in the red skin color of apples, additional minor loci on other chromosomes have been discovered for this trait (Duan et al. 2017; Jung et al. 2022). Our comparison with literature showed ten overlaps with associations for the uncorrelated color features and PCs located on Chr 2 at 20.5 Mbp, Chr 6 at 3.1 Mbp, Chr 8 at 21.9 Mbp, Chr 10 at 2.5 Mbp, Chr 16 at 9.2 Mbp and Chr 17 between 3.1 and 3.8 Mbp (Table S3). The SNP on Chr 17 at 3.8 Mbp for PC1 colocalized exactly with a SNP reported for a visually screened red over color by Jung et al. (2022) (Table S2). This supports the ability of the digital methods based on FruitPhenoBox to precisely localize QTL associated not only with shape and size features, but also with color features. Furthermore, the FruitPhenoBox resulted in three associations with color features found on Chr 3 and 10 that did not colocalize with QTL from the reviewed literature (Table S4), and therefore contributed novel insights into genetic architecture of color traits in apple.

Due to the peaks in the red part of the hue spectrum, which were observed for all rediscovered and novel associations with the color PCs and uncorrelated features (Figure 9, Figure S12), it can be assumed that every reported SNP was associated with red skin color. At the same time, most of the SNPs were associated with similar patterns along the saturation spectrum. However, three SNPs associated with a low saturation uncorrelated feature and PC2 located on Chr 2, 10 and 16 showed shifts in the distributions for the different allele dosages on the saturation spectrum towards higher or lower saturation (Figure 9, Figure S12). Furthermore, PC2 was negatively correlated with ground color scored visually and yellow color scored by the sorting machine in Jung et al. (2022). PC2 was also positively correlated with the low saturation uncorrelated feature associated with the SNPs on Chr 2 and 10 (Figure 3), and a peak was found in the yellow part of the hue spectrum for the SNPs on Chr 2, 10 and 16 (Figure 9, Figure S12). Based on this evidence, the three SNPs may affect the variation in ground color, a trait that has been studied using visual assessment of the gradient between green and yellow skin color in the past (Costa 2015; Jung et al. 2022). The associations with digitally estimated low saturation uncorrelated feature and PC2 contributed further understanding of the trait architecture towards the implementation of marker-based breeding technologies for ground color.

## 5 Conclusion

The FruitPhenoBox provided accurate and detailed fruit phenotypes of a diverse reference population, which were used to discover associations of markers with principal components and uncorrelated features by genome-wide association studies. Digital phenotypes obtained with FruitPhenoBox showed as a powerful solution and efficient alternative to visual phenotyping while providing with novel and rediscovered loci for fruit shape, size, and color, which enhanced our understanding of the variation and the genetics of the studied traits. Using marker-assisted selection, SNPs associated with fruit size may improve the efficiency of breeding resistant varieties with resistances originating from small-fruited genotypes by reducing the number of pseudo-backcrosses needed to produce sufficiently large apples. Loci related to cylindrical or conical fruit shape can assist with breeding for or against exotic fruit shapes. Combined effects of discovered loci on both shape and size suggested genetic interdependence between these traits in apple. The minor loci associated with fruit color contributed novel understanding of trait architecture for the ground color of apple fruits, and they may complement the existing DNA tests for skin color of apples. The results of this study can support apple breeders when improving the process of selecting future apple varieties for their appearance by marker-assisted selection.

## Data availability

Raw imaging data used in this study are available at (TBA). All SNP genotypic data have been deposited at https://doi.org/10.15454/1ERHGX and https://doi.org/10.15454/IOPGYF. The raw supplementary phenotypic data are available at https://doi.org/10.15454/VARJYJ.

## Acknowledgments

The authors thank the field technicians of Agroscope, Waedenswil, Switzerland for maintenance of the orchard and phenotypic data collection.

## References

Bianco L, Cestaro A, Sargent DJ, Banchi E, Derdak S, Di Guardo M, Salvi S, Jansen J, Viola R, Gut I et al. 2014. Development and validation of a 20K single nucleotide polymorphism (SNP) whole genome genotyping array for apple (*Malus* × *domestica* Borkh). PLOS ONE. 9(10):e110377.

Brewer MT, Lang L, Fujimura K, Dujmovic N, Gray S, van der Knaap E. 2006. Development of a controlled vocabulary and software application to analyze fruit shape variation in tomato and other plant species. Plant Physiol. 141(1):15–25.

Brown AG. 1960. The inheritance of shape, size and season of ripening in progenies of the cultivated apple. Euphytica. 9:327–337.

Chagné D, Kirk C, How N, Whitworth C, Fontic C, Reig G, Sawyer G, Rouse S, Poles L, Gardiner SE et al. 2016. A functional genetic marker for apple red skin coloration across different environments. Tree Genet Genom. 12(4):67.

Chang Y, Sun R, Sun H, Zhao Y, Han Y, Chen D, Wang Y, Zhang X, Han Z. 2014. Mapping of quantitative trait loci corroborates independent genetic control of apple size and shape. Scientia Horticulturae. 174:126–132.

Cornille A, Giraud T, Smulders MJM, Roldán-Ruiz I, Gladieux P. 2014. The domestication and evolutionary ecology of apples. Trends Genet. 30(2):57–65.

Costa F. 2015. MetaQTL analysis provides a compendium of genomic loci controlling fruit quality traits in apple. Tree Genet Genom. 11(1):819.

Daccord N, Celton J-M, Linsmith G, Becker C, Choisne N, Schijlen E, van de Geest H, Bianco L, Micheletti D, Velasco R et al. 2017. High-quality de novo assembly of the apple genome and methylome dynamics of early fruit development. Nat Genet. 49(7):1099–1106.

Devoghalaere F, Doucen T, Guitton B, Keeling J, Payne W, Ling TJ, Ross JJ, Hallett IC, Gunaseelan K, Dayatilake GA et al. 2012. A genomics approach to understanding the role of auxin in apple (*Malus* x *domestica*) fruit size control. BMC Plant Biol. 12(1):7.

Duan N, Bai Y, Sun H, Wang N, Ma Y, Li M, Wang X, Jiao C, Legall N, Mao L et al. 2017. Genome re-sequencing reveals the history of apple and supports a two-stage model for fruit enlargement. Nature Communications. 8(1):249.

Espley RV, Hellens RP, Putterill J, Stevenson DE, Kutty-Amma S, Allan AC. 2007. Red colouration in apple fruit is due to the activity of the MYB transcription factor, MdMYB10. Plant J. 49(3):414–427.

Gonzalo MJ, Brewer MT, Anderson C, Sullivan D, Gray S, van der Knaap E. 2009. Tomato fruit shape analysis using morphometric and morphology attributes implemented in Tomato Analyzer software program. J Am Soc Hort Sci. 134(1):77–87.

Hardner CM, Evans K, Brien C, Bliss F, Peace C. 2016. Genetic architecture of apple fruit quality traits following storage and implications for genetic improvement. Tree Genet Genom. 12(2):20.

Huang M, Liu X, Zhou Y, Summers RM, Zhang Z. 2018. BLINK: A package for the next level of genome-wide association studies with both individuals and markers in the millions. GigaScience. 8(2).

Huang Y, Ren Z, Li D, Liu X. 2020. Phenotypic techniques and applications in fruit trees: A review. Plant Methods. 16(1):107.

Jung M, Keller B, Roth M, Aranzana MJ, Auwerkerken A, Guerra W, Al-Rifaï M, Lewandowski M, Sanin N, Rymenants M et al. 2022. Genetic architecture and genomic predictive ability of apple quantitative traits across environments. Hortic Res. 9.

Jung M, Roth M, Aranzana MJ, Auwerkerken A, Bink M, Denancé C, Dujak C, Durel C-E, Font i Forcada C, Cantin CM, et al. 2020. The apple REFPOP—a reference population for genomics-assisted breeding in apple. Hortic Res. 7(1):189.

Kays SJ. 1999. Preharvest factors affecting appearance. Postharvest Biol Technol. 15(3):233–247.

Kenis K, Keulemans J, Davey MW. 2008. Identification and stability of QTLs for fruit quality traits in apple. Tree Genet Genom. 4(4):647–661.

Kirchgessner N, Hodel M, Studer B, Patocchi A, Broggini GA. 2023. FruitPhenoBox – a device for rapid and automated fruit phenotyping of small sample sizes. PREPRINT (Version 1) available at Research Square.

Kuhn M. 2008. Building predictive models in R using the caret package. J Stat Softw. 28(5):1–26.

Lê S, Josse J, Husson F. 2008. FactoMineR: An R package for multivariate analysis. J Stat Softw. 25(1):1–18.

Liao L, Zhang W, Zhang B, Fang T, Wang X-F, Cai Y, Ogutu C, Gao L, Chen G, Nie X et al. 2021. Unraveling a genetic roadmap for improved taste in the domesticated apple. Molecular Plant. 14(9):1454–1471.

Liu Z, Bao D, Liu D, Zhang Y, Ashraf MA, Chen X. 2016. Construction of a genetic linkage map and QTL analysis of fruit-related traits in an F1 Red Fuji x Hongrou apple hybrid. Open Life Sciences. 11(1):487–497.

MATLAB. 2010. MATLAB: 2010. The MathWorks Inc.

Minamikawa MF, Kunihisa M, Noshita K, Moriya S, Abe K, Hayashi T, Katayose Y, Matsumoto T, Nishitani C, Terakami S et al. 2021. Tracing founder haplotypes of Japanese apple varieties: Application in genomic prediction and genome-wide association study. Hortic Res. 8(1):49.

Moriya S, Kunihisa M, Okada K, Shimizu T, Honda C, Yamamoto T, Muranty H, Denancé C, Katayose Y, Iwata H et al. 2017. Allelic composition of MdMYB1 drives red skin color intensity in apple (*Malus* × *domestica* Borkh.) and its application to breeding. Euphytica. 213(4):78.

Musacchi S, Serra S. 2018. Apple fruit quality: Overview on pre-harvest factors. Scientia Horticulturae. 234:409–430.

Pan Y, Wang Y, McGregor C, Liu S, Luan F, Gao M, Weng Y. 2020. Genetic architecture of fruit size and shape variation in cucurbits: A comparative perspective. Theor Appl Genet. 133(1):1–21.

R Core Team. 2014. R: A language and environment for statistical computing. Vienna, Austria: R Foundation for Statistical Computing.

Sierra-Orozco E, Shekasteband R, Illa-Berenguer E, Snouffer A, van der Knaap E, Lee TG, Hutton SF. 2021. Identification and characterization of GLOBE, a major gene controlling fruit shape and impacting fruit size and marketability in tomato. Hortic Res. 8.

Sun HH, Zhao YB, Li CM, Chen DM, Wang Y, Zhang XZ, Han ZH. 2012. Identification of markers linked to major gene loci involved in determination of fruit shape index of apples (*Malus domestica*). Euphytica. 185(2):185–193.

Takos AM, Jaffé FW, Jacob SR, Bogs J, Robinson SP, Walker AR. 2006. Light-induced expression of a MYB gene regulates anthocyanin biosynthesis in red apples. Plant Physiol. 142(3):1216–1232.

Yao J-L, Xu J, Cornille A, Tomes S, Karunairetnam S, Luo Z, Bassett H, Whitworth C, Rees-George J, Ranatunga C et al. 2015. A microRNA allele that emerged prior to apple domestication may underlie fruit size evolution. Plant J. 84(2):417–427.

Zhang C, Serra S, Quirós-Vargas J, Sangjan W, Musacchi S, Sankaran S. 2023. Non-invasive sensing techniques to phenotype multiple apple tree architectures. Information Processing in Agriculture. 10(1):136–147.

Zhang R-p, Wu J, Li X-g, Khan MA, Chen H, Korban SS, Zhang S-l. 2013. An AFLP, SRAP, and SSR genetic linkage map and identification of QTLs for fruit traits in pear (Pyrus L.). Plant Molecular Biology Reporter. 31(3):678–687.

Zine-El-Abidine M, Dutagaci H, Galopin G, Rousseau D. 2021. Assigning apples to individual trees in dense orchards using 3D colour point clouds. Biosys Eng. 209:30–52.

