## Supplemental Materials for "The genetic basis of apple shape and size unraveled by digital phenotyping"

### 1 Supplemental tables

Table S1: Significant SNP associations with principal components (PCs) and uncorrelated features for shape/size and color. Chromosome (Chr), position, $p$ value, minor allelle frequency (MAF), allele dosage, favorable (fav) allele are shown. The explained (expl) variance is based on associated sum of squares regressing all SNPs of a trait (without correction for population structure).

| Trait | Marker | Chr | Position | P value | MAF | Major effect | Allele dosage | Fav allele | Expl variance |
|---|---|---|---|---|---|---|---|---|---|
| Av_SAv_Pol_Rat.Av_AB.LR_Q50 | AX-115235649 | 16 | 3230048 | $1.9 \times 10^{-10}$ | 0.39 | size | 2 | A | 0.06 |
| Av_SAv_Pol_Rat.Av_AB.LR_Q50 | AX-115300506 | 11 | 38069491 | $9.7 \times 10^{-10}$ | 0.17 | conical shape | 0 | G | 0.05 |
| Av_SAv_Pol_Rat.Av_AB.LR_Q50 | AX-115445494 | 8 | 9562443 | $7.7 \times 10^{-8}$ | 0.07 | conical shape | 1 | T | 0.04 |
| Av_SAv_Pol_Rat.Av_AB.LR_Q50 | AX-115631329 | 15 | 47545121 | $7.8 \times 10^{-8}$ | 0.09 | conical shape | 2 | T | 0.03 |
| Av_SAv_Pol_SD_A.LR_Min | AX-115341839 | 1 | 20696409 | $1.5 \times 10^{-7}$ | 0.11 | size | 2 | T | 0.01 |
| Av_SAv_Pol_SD_A.LR_Min | AX-115378613 | 2 | 7626325 | $1.5 \times 10^{-7}$ | 0.14 | size | 2 | T | 0.04 |
| Av_SAv_Pol_SD_A.LR_Min | AX-115506351 | 4 | 30194434 | $6 \times 10^{-8}$ | 0.07 | size | 1 | | 0.07 |
| Av_SAv_Sat_Abs_AB_High | AX-115619261 | 9 | 32475582 | $1 \times 10^{-9}$ | 0.22 | green | 0 | G | 0.06 |
| Av_SAv_Sat_Abs_AB_Medium | AX-115425415 | 9 | 32473327 | $3.4 \times 10^{-8}$ | 0.46 | green | 2 | G | 0.09 |
| Av_SAv_XY_Sum_A_MaxR | AX-115254491 | 7 | 8648220 | $3.7 \times 10^{-9}$ | 0.35 | size | 1 | G | 0.01 |
| Av_SAv_XY_Sum_A_MaxR | AX-115257677 | 5 | 45121659 | $5.1 \times 10^{-8}$ | 0.39 | conical shape | 0 | G | 0.08 |
| Av_SAv_XY_Sum_A_MaxR | AX-115375194 | 6 | 8743643 | $6.4 \times 10^{-14}$ | 0.13 | size | 0 | T | 0.16 |
| Av_SSD_Hue_Abs_AB_Yellow | AX-115210619 | 6 | 3069025 | $1.2 \times 10^{-8}$ | 0.47 | undefinable | | | 0.11 |
| Av_SSD_Pol_Abs_AB_RatM | AX-115327911 | 11 | 4706860 | $4.2 \times 10^{-15}$ | 0.39 | size | 0 | G | 0.07 |
| Av_SSD_Pol_Abs_AB_RatM | AX-115451037 | 3 | 29264257 | $2.6 \times 10^{-9}$ | 0.49 | undefinable | | | 0.01 |
| Av_SSD_Pol_Abs_AB_RatM | AX-115457086 | 8 | 537872 | $1.8 \times 10^{-8}$ | 0.27 | conical shape | 0 | G | 0.04 |
| Av_SSD_Pol_Abs_B.L_SD | AX-115251897 | 3 | 2544800 | $1.3 \times 10^{-8}$ | 0.33 | size | 2 | A | 0.01 |
| Av_SSD_Pol_Abs_B.L_SD | AX-115375737 | 17 | 9806448 | $1.2 \times 10^{-7}$ | 0.48 | size | 2 | A | 0.06 |
| Av_SSD_Pol_Abs_B.L_SD | AX-115436422 | 11 | 15841357 | $9.3 \times 10^{-10}$ | 0.21 | conical shape | 0 | G | 0.03 |
| Av_SSD_Pol_Abs_B.L_SD | AX-115440676 | 11 | 34816193 | $2.2 \times 10^{-10}$ | 0.05 | size | 0 | C | 0.03 |
| Av_SSD_Pol_Abs_B.L_SD | AX-115444766 | 1 | 15580877 | $1.9 \times 10^{-10}$ | 0.05 | size | 0 | T | 0.08 |
| Av_SSD_Pol_Rat.Av_AB.LR_Max | AX-115287563 | 13 | 14542354 | $6.2 \times 10^{-8}$ | 0.24 | conical shape | 1 | T | 0.05 |
| Av_SSD_Pol_Rat.Av_AB.LR_Max | AX-115459245 | 5 | 5031575 | $7.6 \times 10^{-9}$ | 0.41 | conical shape | 0 | G | 0.07 |
| Av_SSD_Pol_Rat.Av_AB.LR_Max | AX-115629265 | 2 | 35251284 | $1.3 \times 10^{-7}$ | 0.06 | conical shape | 0 | C | 0.03 |
| Av_SSD_Pol_Rat.Av_AB.LR_Min | AX-115278645 | 3 | 8512354 | $3.2 \times 10^{-8}$ | 0.26 | size | 2 | T | 0.04 |
| Av_SSD_Pol_Rat.Av_AB.LR_Min | AX-115311704 | 14 | 21571993 | $7.6 \times 10^{-10}$ | 0.15 | size | 0 | C | 0.06 |
| Av_SSD_Pol_Rat.Av_AB.LR_Min | AX-115349165 | 10 | 15436508 | $1.6 \times 10^{-10}$ | 0.11 | cylindrical shape | 0 | T | 0.06 |
| Av_SSD_Pol_Rat.Av_AB.LR_Min | AX-115482171 | 11 | 33111593 | $1.8 \times 10^{-9}$ | 0.39 | conical shape | 0 | A | 0.05 |
| Av_SSD_Pol_Rat.Av_AB.LR_SD | AX-115644112 | 5 | 46356956 | $1.3 \times 10^{-9}$ | 0.11 | conical shape | 0 | T | 0.07 |
| Av_SSD_Pol_SD_B.LR_Max | AX-115283778 | 15 | 48233842 | $2.4 \times 10^{-8}$ | 0.11 | conical shape | 0 | C | 0.06 |
| Av_SSD_Pol_SD_B.LR_Max | AX-115391247 | 9 | 31184244 | $7.4 \times 10^{-8}$ | 0.33 | conical shape | 0 | G | 0.06 |
| Av_SSD_Pol_SD_B.LR_Max | AX-115429431 | 5 | 25209696 | $2.1 \times 10^{-8}$ | 0.41 | conical shape | 0 | A | 0.07 |
| Av_SSD_Pol_SD_B.LR_Max | AX-115449727 | 11 | 6367847 | $1.8 \times 10^{-8}$ | 0.25 | conical shape | 2 | A | 0.04 |
| Av_SSD_Pol_SD_B.LR_Max | AX-115492426 | 6 | 10584742 | $9.9 \times 10^{-9}$ | 0.40 | conical shape | 2 | T | 0.04 |
| Av_SSD_Pol_SD_B.LR_Q25 | AX-115183496 | 1 | 23578695 | $1.5 \times 10^{-7}$ | 0.08 | cylindrical shape | 0 | C | 0.02 |
| Av_SSD_Pol_SD_B.LR_Q25 | AX-115184639 | 10 | 33907766 | $6.4 \times 10^{-8}$ | 0.37 | conical shape | 2 | C | 0.05 |
| Av_SSD_Pol_SD_B.LR_Q25 | AX-115421714 | 11 | 3408716 | $1.1 \times 10^{-8}$ | 0.18 | conical shape | 2 | A | 0.04 |
| Av_SSD_Pol_SD_B.LR_Q25 | AX-115520624 | 3 | 32236169 | $7.4 \times 10^{-8}$ | 0.37 | size | 2 | A | 0.04 |
| Av_SSD_Pol_SD_B.LR_Q25 | AX-115655116 | 2 | 7872781 | $1.1 \times 10^{-8}$ | 0.15 | size | 2 | A | 0.00 |
| Av_SSD_Pol_SD_R_Sym.rel | AX-115389297 | 13 | 13849729 | $9.6 \times 10^{-10}$ | 0.18 | symmetric shape | | | 0.06 |
| Av_SSD_Pol_SD_R_Sym.rel | AX-115461813 | 15 | 10680808 | $3.3 \times 10^{-10}$ | 0.32 | symmetric shape | | | 0.07 |
| Av_SSD_Pol_SD_R_Sym.rel | AX-115515574 | 16 | 3452384 | $2.3 \times 10^{-8}$ | 0.46 | symmetric shape | | | 0.06 |
| Av_SSD_X_Abs_A_Q25 | AX-115221494 | 1 | 17152361 | $9.6 \times 10^{-10}$ | 0.07 | size | 2 | T | 0.04 |
| Av_SSD_X_Abs_A_Q25 | AX-115363036 | 14 | 15862103 | $4.5 \times 10^{-8}$ | 0.17 | size | 0 | T | 0.10 |
| Av_SSD_X_Abs_A_Q25 | AX-115442384 | 9 | 4698417 | $1.2 \times 10^{-8}$ | 0.08 | size | 2 | G | 0.02 |
| Av_SSD_X_Abs_A_Q25 | AX-115444766 | 1 | 15580877 | $7.4 \times 10^{-11}$ | 0.05 | size | 0 | T | 0.09 |
| Av_SSD_X_Abs_B_Q25 | AX-115319264 | 6 | 4761377 | $1.1 \times 10^{-7}$ | 0.05 | size | 0 | A | 0.04 |
| Av_SSD_X_Abs_B_Q25 | AX-115473970 | 6 | 23085308 | $9 \times 10^{-8}$ | 0.14 | size | 0 | A | 0.06 |
| Av_SSD_X_Abs_B_Q25 | AX-115500353 | 9 | 33000755 | $6.8 \times 10^{-8}$ | 0.29 | conical shape | 0 | G | 0.03 |
| Av_SSD_X_Abs_B_Q25 | AX-115521252 | 9 | 5170431 | $2.8 \times 10^{-8}$ | 0.45 | size | 1 | G | 0.04 |
| Av_SSD_X_Abs_B_Q25 | AX-115625798 | 1 | 15868916 | $3.2 \times 10^{-12}$ | 0.06 | size | 0 | C | 0.10 |
| PC1_color | AX-115182287 | 10 | 38213067 | $7.9 \times 10^{-9}$ | 0.47 | undefinable | | | 0.01 |
| PC1_color | AX-115321298 | 3 | 6282194 | $1.6 \times 10^{-8}$ | 0.26 | undefinable | | | 0.05 |
| PC1_color | AX-115452523 | 17 | 3769755 | $1.5 \times 10^{-8}$ | 0.31 | undefinable | | | 0.02 |
| PC1_color | AX-115505043 | 9 | 33744382 | $5.6 \times 10^{-22}$ | 0.46 | green | 2 | G | 0.33 |
| PC1_color | AX-115598053 | 8 | 21899808 | $2.3 \times 10^{-8}$ | 0.14 | undefinable | | | 0.03 |
| PC1_shape_size | AX-115227580 | 10 | 35210847 | $1.6 \times 10^{-7}$ | 0.30 | size | 0 | G | 0.04 |
| PC1_shape_size | AX-115318835 | 6 | 20393412 | $9.9 \times 10^{-9}$ | 0.15 | size | 0 | T | 0.09 |
| PC1_shape_size | AX-115345243 | 14 | 24020772 | $2.9 \times 10^{-9}$ | 0.14 | size | 2 | C | 0.03 |
| PC1_shape_size | AX-115380250 | 1 | 31575453 | $1.2 \times 10^{-7}$ | 0.25 | size | 0 | C | 0.07 |
| PC1_shape_size | AX-115460130 | 7 | 3024174 | $1.5 \times 10^{-7}$ | 0.16 | size | 2 | C | 0.04 |
| PC1_shape_size | AX-115534322 | 2 | 12130252 | $3.7 \times 10^{-8}$ | 0.43 | size | 0 | A | 0.03 |
| PC1_shape_size | AX-115628310 | 14 | 1478099 | $3 \times 10^{-9}$ | 0.34 | size | 2 | G | 0.05 |
| PC2_color | AX-115244827 | 16 | 9227750 | $4.5 \times 10^{-11}$ | 0.37 | yellow | 2 | T | 0.08 |
| PC2_shape_size | AX-115330695 | 11 | 3241944 | $1.9 \times 10^{-9}$ | 0.18 | conical shape | 2 | G | 0.05 |
| PC2_shape_size | AX-115429431 | 5 | 25209696 | $7 \times 10^{-8}$ | 0.41 | conical shape | 0 | A | 0.03 |
| PC2_shape_size | AX-115490525 | 2 | 7904059 | $1.2 \times 10^{-7}$ | 0.18 | conical shape | 0 | A | 0.00 |
| PC2_shape_size | AX-115548370 | 13 | 5411038 | $9.1 \times 10^{-9}$ | 0.22 | conical shape | 2 | G | 0.03 |
| PC2_shape_size | AX-115552368 | 10 | 33469975 | $5.5 \times 10^{-9}$ | 0.47 | conical shape | 0 | G | 0.09 |
| PC3_shape_size | AX-115254276 | 7 | 21705520 | $2.6 \times 10^{-11}$ | 0.23 | conical shape | 2 | T | 0.12 |
| PC3_shape_size | AX-115404241 | 10 | 19586733 | $2.2 \times 10^{-8}$ | 0.25 | conical shape | 0 | C | 0.03 |
| PC3_shape_size | AX-115507212 | 9 | 33255077 | $2.5 \times 10^{-8}$ | 0.35 | conical shape | 0 | T | 0.04 |
| PC4_color | AX-105213720 | 9 | 33801013 | $2.7 \times 10^{-10}$ | 0.48 | green | 2 | C | 0.10 |
| PC4_shape_size | AX-115265526 | 11 | 41448132 | $1.3 \times 10^{-11}$ | 0.19 | conical shape | 0 | A | 0.04 |
| PC4_shape_size | AX-115419379 | 10 | 17661953 | $4.2 \times 10^{-8}$ | 0.15 | cylindrical shape | 0 | C | 0.05 |
| PC4_shape_size | AX-115476093 | 12 | 9098303 | $3.2 \times 10^{-8}$ | 0.20 | size | 0 | T | 0.06 |
| PC4_shape_size | AX-115659654 | 11 | 12416234 | $1.4 \times 10^{-7}$ | 0.19 | conical shape | 0 | G | 0.04 |
| PC5_shape_size | AX-115538412 | 1 | 15918343 | $3.1 \times 10^{-8}$ | 0.08 | conical shape | 2 | T | 0.01 |
| PC5_shape_size | AX-115629269 | 8 | 21161287 | $3.2 \times 10^{-8}$ | 0.06 | size | 0 | A | 0.02 |
| SD_SAv_Hue_Abs_AB_Red | AX-105213720 | 9 | 33801013 | $9.9 \times 10^{-35}$ | 0.49 | green | 2 | C | 0.42 |
| SD_SAv_Hue_Abs_AB_Yellow | AX-115332892 | 9 | 33729410 | $1.3 \times 10^{-16}$ | 0.49 | green | 2 | C | 0.15 |
| SD_SAv_Hue_Abs_AB_Yellow | AX-115394842 | 17 | 3024389 | $1.5 \times 10^{-8}$ | 0.25 | undefinable | | | 0.03 |
| SD_SSD_Sat_Abs_AB_Low | AX-115228698 | 10 | 2507193 | $2.2 \times 10^{-9}$ | 0.41 | yellow | 2 | C | 0.05 |
| SD_SSD_Sat_Abs_AB_Low | AX-115316726 | 2 | 20512335 | $1.5 \times 10^{-9}$ | 0.30 | yellow | 0 | A | 0.09 |
| SD_SSD_Sat_Abs_AB_Low | AX-115536962 | 10 | 41138070 | $9.3 \times 10^{-8}$ | 0.34 | undefinable | | | 0.01 |

Table S2: Overlap between QTL identified in the current and previous studies. The physical difference between the positions is given in bp. Only differences below 100'000 bp were considered.

| Chr | Position | Marker | Position_ref | Marker_ref | Trait | Publication | Difference | Trait_ref | Remark |
|---|---|---|---|---|---|---|---|---|---|
| 1 | 15868916 | AX-115625798 | 15867063 | Chr01_15867063 | Av_SSD_X_Abs_B_Q25 | Liao et al. (2021) | 1853 | Oxalate | 2 years, China |
| 1 | 15918343 | AX-115538412 | 15906474 | Chr01_15906474 | PC5_shape_size | Liao et al. (2021) | 11869 | Oxalate | 2 years, China |
| 1 | 23578695 | AX-115183496 | 23552731 | AX-115322263 | Av_SSD_Pol_SD_B.LR_Q25 | Jung et al. (2022) | 25964 | Number of fruits | location-specific CHE |
| 1 | 23578695 | AX-115183496 | 23578926 | SNP_FB_0431170 | Av_SSD_Pol_SD_B.LR_Q25 | Minamikawa et al. (2021) | -231 | Firm | Across years, Japan |
| 2 | 7626325 | AX-115378613 | 7721047 | MDP0000843352 | Av_SAv_Pol_SD_A.LR_Min | Duan et al. (2017) | -94722 | Fruit development | Peroxidase |
| 2 | 7626325 | AX-115378613 | 7697242 | MDP0000293312 | Av_SAv_Pol_SD_A.LR_Min | Duan et al. (2017) | -70917 | Fruit development | Phosphatidylserine decarboxylase proenzyme 2 |
| 2 | 7872781 | AX-115655116 | 7881043 | MDP0000327940 | Av_SSD_Pol_SD_B.LR_Q25 | Duan et al. (2017) | -8262 | Fruit development | Chlorophyll a-b binding protein, chloroplastic |
| 2 | 7872781 | AX-115655116 | 7869905 | MDP0000232805 | Av_SSD_Pol_SD_B.LR_Q25 | Duan et al. (2017) | 2876 | Fruit development | Lipid transfer protein |
| 2 | 7904059 | AX-115490525 | 7883091 | MDP0000232809 | PC2_shape_size | Duan et al. (2017) | 20968 | Fruit development | 3-hydroxyacyl-[acyl-carrier-protein] dehydratase FabZ |
| 3 | 32236169 | AX-115520624 | 32328637 | AX-115621973 | Av_SSD_Pol_SD_B.LR_Q25 | Jung et al. (2022) | -92468 | Floral emergence | location-specific BEL |
| 4 | 30194434 | AX-115506351 | 30124476 | SNP_FB_0591255 | Av_SAv_Pol_SD_A.LR_Min | Minamikawa et al. (2021) | 69958 | Mealiness | Across years, Japan |
| 5 | 5031575 | AX-115459245 | 4949273 | AX-115392975 | Av_SSD_Pol_Rat.Av_AB.LR_Max | Jung et al. (2022) | 82302 | Fruit firmness | location-specific CHE |
| 5 | 25209696 | AX-115429431 | 25236549 | AX-115413471 | Av_SSD_Pol_SD_B.LR_Max | Jung et al. (2022) | -26853 | Flowering intensity | across-location |
| 6 | 4761377 | AX-115319264 | 4815618 | AX-115220470 | Av_SSD_X_Abs_B_Q25 | Jung et al. (2022) | -54241 | Russet freq. - overall | location-specific FRA |
| 7 | 21705520 | AX-115254276 | 21735820 | MDP0000336563 | PC3_shape_size | Duan et al. (2017) | -30300 | Fruit development | Synovial sarcoma associated ss18 protein, putative |
| 7 | 21705520 | AX-115254276 | 21723408 | MDP0000189381 | PC3_shape_size | Duan et al. (2017) | -17888 | Fruit development | Putative calmodulin-like protein |
| 8 | 21161287 | AX-115629269 | 21158378 | MDP0000231982 | PC5_shape_size | Duan et al. (2017) | 2909 | Fruit development | Pentatricopeptide repeat-containing protein |
| 8 | 21161287 | AX-115629269 | 21161135 | MDP0000507227 | PC5_shape_size | Duan et al. (2017) | 152 | Fruit development | UDP-glucose 4-epimerase, putative |
| 10 | 15436508 | AX-115349165 | 15373507 | MDP0000161978 | Av_SSD_Pol_Rat.Av_AB.LR_Min | Duan et al. (2017) | 63001 | Fruit development | Fatty acid desaturase, putative |
| 16 | 3230048 | AX-115235649 | 3220684 | AX-115602744 | Av_SAv_Pol_Rat.Av_AB.LR_Q50 | Jung et al. (2022) | 9364 | Titratable acidity | location-specific CHE |
| 16 | 3230048 | AX-115235649 | 3229804 | Chr16_3229804 | Av_SAv_Pol_Rat.Av_AB.LR_Q50 | Liao et al. (2021) | 244 | Malate-1 | 2 years, China |
| 16 | 3452384 | AX-115515574 | 3479443 | SNP_FB_0336036 | Av_SSD_Pol_SD_R_Sym.rel | Minamikawa et al. (2021) | -27059 | CraTop | Across years, Japan |
| 16 | 3452384 | AX-115515574 | 3452384 | Chr16_3452384 | Av_SSD_Pol_SD_R_Sym.rel | Liao et al. (2021) | 0 | Malate-1 | 2 years, China |
| 6 | 3069025 | AX-115210619 | 3070483 | AX-115210618 | Av_SSD_Hue_Abs_AB_Yellow | Jung et al. (2022) | -1458 | Red over color | location-specific ESP |
| 9 | 33729410 | AX-115332892 | 33801319 | s9_33801319 | SD_SAv_Hue_Abs_AB_Yellow | Duan et al. (2017) | -71909 | Skin color | |
| 9 | 33729410 | AX-115332892 | 33799120 | AX-115279459 | SD_SAv_Hue_Abs_AB_Yellow | Jung et al. (2022) | -69710 | Red over color | location-specific ITA |
| 9 | 33801013 | AX-105213720 | 33801013 | AX-105213720 | PC4_color | Jung et al. (2022) | 0 | Red over color | across-location |
| 9 | 33801013 | AX-105213720 | 33801013 | AX-105213720 | PC4_color | Jung et al. (2022) | 0 | Green color* | location-specific CHE |
| 10 | 38213067 | AX-115182287 | 38218534 | AX-115263961 | PC1_color | Jung et al. (2022) | -5467 | Trunk diameter | location-specific FRA |
| 10 | 38213067 | AX-115182287 | 38217999 | AX-115263958 | PC1_color | Jung et al. (2022) | -4932 | Green color* | location-specific CHE |
| 10 | 41138070 | AX-115536962 | 41129257 | Chr10_41129257 | SD_SSD_Sat_Abs_AB_Low | Liao et al. (2021) | 8813 | Citrate | 2 years, China |
| 16 | 9227750 | AX-115244827 | 9244325 | AX-115244807 | PC2_color | Jung et al. (2022) | -16575 | Ground color | across-location |
| 16 | 9227750 | AX-115244827 | 9240764 | AX-115244813 | PC2_color | Jung et al. (2022) | -13014 | Harvest date | across-location |
| 17 | 3024389 | AX-115394842 | 3119571 | AX-115231505 | SD_SAv_Hue_Abs_AB_Yellow | Jung et al. (2022) | -95182 | Water core grade* | location-specific ITA |
| 17 | 3024389 | AX-115394842 | 3096111 | AX-115407463 | SD_SAv_Hue_Abs_AB_Yellow | Jung et al. (2022) | -71722 | Russet cover | location-specific ESP |
| 17 | 3769755 | AX-115452523 | 3823904 | AX-115394824 | PC1_color | Jung et al. (2022) | -54149 | Red over color | location-specific BEL |
| 17 | 3769755 | AX-115452523 | 3769755 | AX-115452523 | PC1_color | Jung et al. (2022) | 0 | Red over color | location-specific ESP |

Table S3: Identified markers with potentially overlapping positions reported in previous studies.

(continuous on next page)

| Chr | Position | Segment | Marker | Feature | Publication | Trait in reference |
|---|---|---|---|---|---|---|
| 1 | 23552731 | bottom | AX-115183496 | Av_SSD_Pol_SD_B.LR_Q25 | Jung et al. (2022) | Number of fruits |
| 1 | 23578695 | bottom | AX-115183496 | Av_SSD_Pol_SD_B.LR_Q25 | Costa (2015) | Fruit height |
| 1 | 31575453 | bottom | AX-115380250 | PC1_shape | Costa (2015) | Fruit height |
| 2 | 35251284 | bottom | AX-115629265 | Av_SSD_Pol_Rat.Av_AB.LR_Max | Chang et al. (2014) | Fruit length, diameter or size |
| 3 | 2544800 | top | AX-115251897 | Av_SSD_Pol_Abs_B.L_SD | Sun et al. (2015) | Fruit weight |
| 3 | 8512354 | top | AX-115278645 | Av_SSD_Pol_Rat.Av_AB.LR_Min | Sun et al. (2015) | Fruit weight |
| 3 | 29264257 | bottom | AX-115451037 | Av_SSD_Pol_Abs_AB_RatM | Chang et al. (2014) | Fruit length, diameter or size |
| 3 | 29264257 | bottom | AX-115451037 | Av_SSD_Pol_Abs_AB_RatM | Potts et al. (2014) | Fruit diameter |
| 3 | 29264257 | bottom | AX-115451037 | Av_SSD_Pol_Abs_AB_RatM | Potts et al. (2014) | Weight |
| 3 | 29264257 | bottom | AX-115451037 | Av_SSD_Pol_Abs_AB_RatM | Potts et al. (2014) | Length |
| 3 | 32236169 | bottom | AX-115520624 | Av_SSD_Pol_SD_B.LR_Q25 | Chang et al. (2014) | Fruit length, diameter or size |
| 3 | 32236169 | bottom | AX-115520624 | Av_SSD_Pol_SD_B.LR_Q25 | Potts et al. (2014) | Fruit diameter |
| 3 | 32236169 | bottom | AX-115520624 | Av_SSD_Pol_SD_B.LR_Q25 | Potts et al. (2014) | Weight |
| 3 | 32236169 | bottom | AX-115520624 | Av_SSD_Pol_SD_B.LR_Q25 | Potts et al. (2014) | Length |
| 5 | 5031575 | top | AX-115459245 | Av_SSD_Pol_Rat.Av_AB.LR_Max | Chang et al. (2014) | Ratio between height and diameter |
| 5 | 5031575 | top | AX-115459245 | Av_SSD_Pol_Rat.Av_AB.LR_Max | Minamikawa et al. (2021) | Fruit weight in grams |
| 5 | 25209696 | center | AX-115429431 | Av_SSD_Pol_SD_B.LR_Max | Potts et al. (2014) | Length |
| 5 | 25209696 | center | AX-115429431 | PC2_shape | Potts et al. (2014) | Length |
| 5 | 25209696 | center | AX-115429431 | Av_SSD_Pol_SD_B.LR_Max | Sun et al. (2015) | Fruit weight |
| 5 | 25209696 | center | AX-115429431 | PC2_shape | Sun et al. (2015) | Fruit weight |
| 5 | 25209696 | center | AX-115429431 | Av_SSD_Pol_SD_B.LR_Max | Potts et al. (2014) | Weight |
| 5 | 25209696 | center | AX-115429431 | PC2_shape | Potts et al. (2014) | Weight |
| 5 | 25209696 | center | AX-115429431 | Av_SSD_Pol_SD_B.LR_Max | Minamikawa et al. (2021) | Fruit weight in grams |
| 5 | 25209696 | center | AX-115429431 | PC2_shape | Minamikawa et al. (2021) | Fruit weight in grams |
| 5 | 25209696 | center | AX-115429431 | Av_SSD_Pol_SD_B.LR_Max | Potts et al. (2014) | Fruit diameter |
| 5 | 25209696 | center | AX-115429431 | PC2_shape | Potts et al. (2014) | Fruit diameter |
| 5 | 25209696 | center | AX-115429431 | Av_SSD_Pol_SD_B.LR_Max | Potts et al. (2014) | Length |
| 5 | 25209696 | center | AX-115429431 | PC2_shape | Potts et al. (2014) | Length |
| 5 | 25209696 | center | AX-115429431 | Av_SSD_Pol_SD_B.LR_Max | Sun et al. (2015) | Fruit weight |
| 5 | 25209696 | center | AX-115429431 | PC2_shape | Sun et al. (2015) | Fruit weight |
| 5 | 25209696 | center | AX-115429431 | Av_SSD_Pol_SD_B.LR_Max | Potts et al. (2014) | Weight |
| 5 | 25209696 | center | AX-115429431 | PC2_shape | Potts et al. (2014) | Weight |
| 5 | 25209696 | center | AX-115429431 | Av_SSD_Pol_SD_B.LR_Max | Minamikawa et al. (2021) | Fruit weight in grams |
| 5 | 25209696 | center | AX-115429431 | PC2_shape | Minamikawa et al. (2021) | Fruit weight in grams |
| 5 | 25209696 | center | AX-115429431 | Av_SSD_Pol_SD_B.LR_Max | Potts et al. (2014) | Fruit diameter |
| 5 | 25209696 | center | AX-115429431 | PC2_shape | Potts et al. (2014) | Fruit diameter |
| 5 | 45121659 | bottom | AX-115257677 | Av_SAv_XY_Sum_A_MaxR | Chang et al. (2014) | Fruit length, diameter or size |
| 5 | 46356956 | bottom | AX-115644112 | Av_SSD_Pol_Rat.Av_AB.LR_SD | Chang et al. (2014) | Fruit length, diameter or size |
| 6 | 20393412 | center | AX-115318835 | PC1_shape | Liu et al. (2016) | Fruit weight |
| 6 | 20393412 | center | AX-115318835 | PC1_shape | Liu et al. (2016) | Fruit diameter |
| 6 | 23085308 | center | AX-115473970 | Av_SSD_X_Abs_B_Q25 | Liu et al. (2016) | Fruit weight |
| 6 | 23085308 | center | AX-115473970 | Av_SSD_X_Abs_B_Q25 | Liu et al. (2016) | Fruit diameter |
| 7 | 3024174 | top | AX-115460130 | PC1_shape | Costa (2015) | Fruit diameter |
| 7 | 3024174 | top | AX-115460130 | PC1_shape | Costa (2015) | Fruit size (1-5 scale) |
| 7 | 3024174 | top | AX-115460130 | PC1_shape | Costa (2015) | Weight |
| 7 | 3024174 | top | AX-115460130 | PC1_shape | Costa (2015) | Fruit height |
| 7 | 3024174 | top | AX-115460130 | PC1_shape | Chang et al. (2014) | Ratio between height and diameter |
| 7 | 8648220 | top | AX-115254491 | Av_SAv_XY_Sum_A_MaxR | Costa (2015) | Fruit diameter |
| 7 | 8648220 | top | AX-115254491 | Av_SAv_XY_Sum_A_MaxR | Costa (2015) | Fruit size (1-5 scale) |
| 7 | 8648220 | top | AX-115254491 | Av_SAv_XY_Sum_A_MaxR | Costa (2015) | Weight |
| 7 | 8648220 | top | AX-115254491 | Av_SAv_XY_Sum_A_MaxR | Costa (2015) | Fruit height |
| 7 | 8648220 | top | AX-115254491 | Av_SAv_XY_Sum_A_MaxR | Chang et al. (2014) | Ratio between height and diameter |
| 8 | 537872 | top | AX-115457086 | Av_SSD_Pol_Abs_AB_RatM | Chang et al. (2014) | Fruit length, diameter or size |
| 8 | 9562443 | top | AX-115445494 | Av_SAv_Pol_Rat.Av_AB.LR_Q50 | Chang et al. (2014) | Fruit length, diameter or size |
| 8 | 21161287 | bottom | AX-115629269 | PC5_shape | Chang et al. (2014) | Fruit length, diameter or size |
| 9 | 31184244 | bottom | AX-115391247 | Av_SSD_Pol_SD_B.LR_Max | Chang et al. (2014) | Ratio between height and diameter |
| 9 | 31184244 | bottom | AX-115391247 | Av_SSD_Pol_SD_B.LR_Max | Chang et al. (2014) | Fruit length, diameter or size |
| 9 | 33000755 | bottom | AX-115500353 | Av_SSD_X_Abs_B_Q25 | Chang et al. (2014) | Ratio between height and diameter |
| 9 | 33000755 | bottom | AX-115500353 | Av_SSD_X_Abs_B_Q25 | Chang et al. (2014) | Fruit length, diameter or size |
| 9 | 33255077 | bottom | AX-115507212 | PC3_shape | Chang et al. (2014) | Ratio between height and diameter |
| 9 | 33255077 | bottom | AX-115507212 | PC3_shape | Chang et al. (2014) | Fruit length, diameter or size |
| 10 | 15436508 | center | AX-115349165 | Av_SSD_Pol_Rat.Av_AB.LR_Min | Kenis et al. (2008) | Mean fruit fresh weight |
| 10 | 15436508 | center | AX-115349165 | Av_SSD_Pol_Rat.Av_AB.LR_Min | Sun et al. (2012) | Ratio between height and diameter |
| 10 | 15436508 | center | AX-115349165 | Av_SSD_Pol_Rat.Av_AB.LR_Min | Kenis et al. (2008) | Mean fruit diameter |
| 10 | 17661953 | center | AX-115419379 | PC4_shape | Kenis et al. (2008) | Mean fruit fresh weight |
| 10 | 17661953 | center | AX-115419379 | PC4_shape | Sun et al. (2012) | Ratio between height and diameter |
| 10 | 17661953 | center | AX-115419379 | PC4_shape | Kenis et al. (2008) | Mean fruit diameter |
| 10 | 19586733 | center | AX-115404241 | PC3_shape | Kenis et al. (2008) | Mean fruit fresh weight |
| 10 | 19586733 | center | AX-115404241 | PC3_shape | Sun et al. (2012) | Ratio between height and diameter |
| 10 | 19586733 | center | AX-115404241 | PC3_shape | Kenis et al. (2008) | Mean fruit diameter |
| 10 | 33469975 | bottom | AX-115552368 | PC2_shape | Sun et al. (2012) | Ratio between height and diameter |
| 10 | 33907766 | bottom | AX-115184639 | Av_SSD_Pol_SD_B.LR_Q25 | Sun et al. (2012) | Ratio between height and diameter |
| 10 | 35210847 | bottom | AX-115227580 | PC1_shape | Sun et al. (2012) | Ratio between height and diameter |
| 11 | 3241944 | top | AX-115330695 | PC2_shape | Costa (2015) | Fruit size (1-5 scale) |
| 11 | 3241944 | top | AX-115330695 | PC2_shape | Chang et al. (2014) | Ratio between height and diameter |
| 11 | 3241944 | top | AX-115330695 | PC2_shape | Chang et al. (2014) | Fruit length, diameter or size |
| 11 | 3241944 | top | AX-115330695 | PC2_shape | Sun et al. (2012) | Ratio between height and diameter |
| 11 | 3408716 | top | AX-115421714 | Av_SSD_Pol_SD_B.LR_Q25 | Costa (2015) | Fruit size (1-5 scale) |
| 11 | 3408716 | top | AX-115421714 | Av_SSD_Pol_SD_B.LR_Q25 | Chang et al. (2014) | Ratio between height and diameter |
| 11 | 3408716 | top | AX-115421714 | Av_SSD_Pol_SD_B.LR_Q25 | Chang et al. (2014) | Fruit length, diameter or size |
| 11 | 3408716 | top | AX-115421714 | Av_SSD_Pol_SD_B.LR_Q25 | Sun et al. (2012) | Ratio between height and diameter |
| 11 | 4706860 | top | AX-115327911 | Av_SSD_Pol_Abs_AB_RatM | Costa (2015) | Fruit size (1-5 scale) |
| 11 | 4706860 | top | AX-115327911 | Av_SSD_Pol_Abs_AB_RatM | Chang et al. (2014) | Ratio between height and diameter |
| 11 | 4706860 | top | AX-115327911 | Av_SSD_Pol_Abs_AB_RatM | Chang et al. (2014) | Fruit length, diameter or size |
| 11 | 4706860 | top | AX-115327911 | Av_SSD_Pol_Abs_AB_RatM | Sun et al. (2012) | Ratio between height and diameter |
| 11 | 6367847 | top | AX-115449727 | Av_SSD_Pol_SD_B.LR_Max | Costa (2015) | Fruit size (1-5 scale) |
| 11 | 6367847 | top | AX-115449727 | Av_SSD_Pol_SD_B.LR_Max | Chang et al. (2014) | Ratio between height and diameter |
| 11 | 6367847 | top | AX-115449727 | Av_SSD_Pol_SD_B.LR_Max | Chang et al. (2014) | Fruit length, diameter or size |
| 11 | 6367847 | top | AX-115449727 | Av_SSD_Pol_SD_B.LR_Max | Sun et al. (2012) | Ratio between height and diameter |
| 11 | 12416234 | top | AX-115659654 | PC4_shape | Costa (2015) | Fruit size (1-5 scale) |
| 11 | 12416234 | top | AX-115659654 | PC4_shape | Chang et al. (2014) | Ratio between height and diameter |
| 11 | 12416234 | top | AX-115659654 | PC4_shape | Chang et al. (2014) | Fruit length, diameter or size |
| 11 | 12416234 | top | AX-115659654 | PC4_shape | Sun et al. (2012) | Ratio between height and diameter |
| 11 | 15841357 | center | AX-115436422 | Av_SSD_Pol_Abs_B.L_SD | Minamikawa et al. (2021) | Fruit weight in grams |
| 11 | 15841357 | center | AX-115436422 | Av_SSD_Pol_Abs_B.L_SD | Sun et al. (2012) | Ratio between height and diameter |
| 11 | 33111593 | bottom | AX-115482171 | Av_SSD_Pol_Rat.Av_AB.LR_Min | Sun et al. (2012) | Ratio between height and diameter |
| 11 | 34816193 | bottom | AX-115440676 | Av_SSD_Pol_Abs_B.L_SD | Sun et al. (2012) | Ratio between height and diameter |
| 11 | 38069491 | bottom | AX-115300506 | Av_SAv_Pol_Rat.Av_AB.LR_Q50 | Sun et al. (2012) | Ratio between height and diameter |
| 11 | 41448132 | bottom | AX-115265526 | PC4_shape | Sun et al. (2012) | Ratio between height and diameter |
| 12 | 9098303 | top | AX-115476093 | PC4_shape | Sun et al. (2012) | Ratio between height and diameter |
| 13 | 5411038 | top | AX-115548370 | PC2_shape | Minamikawa et al. (2021) | Fruit weight in grams |
| 13 | 5411038 | top | AX-115548370 | PC2_shape | Sun et al. (2012) | Ratio between height and diameter |
| 13 | 13849729 | top | AX-115389297 | Av_SSD_Pol_SD_R_Sym.rel | Minamikawa et al. (2021) | Fruit weight in grams |
| 13 | 13849729 | top | AX-115389297 | Av_SSD_Pol_SD_R_Sym.rel | Sun et al. (2012) | Ratio between height and diameter |
| 13 | 14542354 | top | AX-115287563 | Av_SSD_Pol_Rat.Av_AB.LR_Max | Minamikawa et al. (2021) | Fruit weight in grams |
| 13 | 14542354 | top | AX-115287563 | Av_SSD_Pol_Rat.Av_AB.LR_Max | Sun et al. (2012) | Ratio between height and diameter |

| Chr | Position | Segment | Marker | Feature | Publication | Trait in reference |
|---|---|---|---|---|---|---|
| 14 | 24020772 | bottom | AX-115345243 | PC1_shape | Chang et al. (2014) | Fruit length, diameter or size |
| 15 | 10680808 | top | AX-115461813 | Av_SSD_Pol_SD_R_Sym.rel | Minamikawa et al. (2021) | Fruit weight in grams |
| 15 | 10680808 | top | AX-115461813 | Av_SSD_Pol_SD_R_Sym.rel | Chang et al. (2014) | Ratio between height and diameter |
| 15 | 10680808 | top | AX-115461813 | Av_SSD_Pol_SD_R_Sym.rel | Chang et al. (2014) | Fruit length, diameter or size |
| 15 | 47545121 | bottom | AX-115631329 | Av_SAv_Pol_Rat.Av_AB.LR_Q50 | Devoghalaere et al. (2012) | Fruit size - single fruit weight |
| 15 | 48233842 | bottom | AX-115283778 | Av_SSD_Pol_SD_B.LR_Max | Devoghalaere et al. (2012) | Fruit size - single fruit weight |
| 16 | 3230048 | top | AX-115235649 | Av_SAv_Pol_Rat.Av_AB.LR_Q50 | Minamikawa et al. (2021) | Fruit weight in grams |
| 16 | 3452384 | top | AX-115515574 | Av_SSD_Pol_SD_R_Sym.rel | Minamikawa et al. (2021) | Fruit weight in grams |
| 17 | 9806448 | top | AX-115375737 | Av_SSD_Pol_Abs_B.L_SD | Costa (2015) | Fruit size (1-5 scale) |
| 17 | 9806448 | top | AX-115375737 | Av_SSD_Pol_Abs_B.L_SD | Chang et al. (2014) | Ratio between height and diameter |
| 17 | 9806448 | top | AX-115375737 | Av_SSD_Pol_Abs_B.L_SD | Costa (2015) | Fruit height |
| 17 | 9806448 | top | AX-115375737 | Av_SSD_Pol_Abs_B.L_SD | Chang et al. (2014) | Fruit length, diameter or size |
| 2 | 20512335 | center | AX-115316726 | SD_SSD_Sat_Abs_AB_Low | Duan et al. (2017) | Six levels of fruit skin color |
| 6 | 3070483 | top | AX-115210619 | Av_SSD_Hue_Abs_AB_Yellow | Jung et al. (2022) | Red over color |
| 8 | 21899808 | bottom | AX-115598053 | PC1_color | Minamikawa et al. (2021) | Degree of skin coloration in percent (1-5 scale) |
| 9 | 32473327 | bottom | AX-115425415 | Av_SAv_Sat_Abs_AB_Medium | Costa (2015) | Overcolor % |
| 9 | 32473327 | bottom | AX-115425415 | Av_SAv_Sat_Abs_AB_Medium | Amyotte et al. (2017) | Presence or absence of red skin colour |
| 9 | 32473327 | bottom | AX-115425415 | Av_SAv_Sat_Abs_AB_Medium | Kunihisa et al. (2014) | “Depth” of the red coloration |
| 9 | 32473327 | bottom | AX-115425415 | Av_SAv_Sat_Abs_AB_Medium | Migicovsky et al. (2021) | Red versus green |
| 9 | 32473327 | bottom | AX-115425415 | Av_SAv_Sat_Abs_AB_Medium | Duan et al. (2017) | Six levels of fruit skin color |
| 9 | 32473327 | bottom | AX-115425415 | Av_SAv_Sat_Abs_AB_Medium | Chagné et al. (2016) | Percentage overcolor |
| 9 | 32473327 | bottom | AX-115425415 | Av_SAv_Sat_Abs_AB_Medium | Migicovsky et al. (2016) | Overcolor and overcolor intesity |
| 9 | 32473327 | bottom | AX-115425415 | Av_SAv_Sat_Abs_AB_Medium | Lee et al. (2017) | Skin chroma (color intensity) |
| 9 | 32473327 | bottom | AX-115425415 | Av_SAv_Sat_Abs_AB_Medium | Kumar et al. (2014) | Overcolor on zero to nine scale |
| 9 | 32473327 | bottom | AX-115425415 | Av_SAv_Sat_Abs_AB_Medium | Minamikawa et al. (2021) | Degree of skin coloration in percent (1-5 scale) |
| 9 | 32473327 | bottom | AX-115425415 | Av_SAv_Sat_Abs_AB_Medium | Moriya et al. (2017) | Overall color intensity |
| 9 | 32473327 | bottom | AX-115425415 | Av_SAv_Sat_Abs_AB_Medium | McClure et al. (2018) | % red blush covering the surface |
| 9 | 32473327 | bottom | AX-115425415 | Av_SAv_Sat_Abs_AB_Medium | Zheng et al. (2020) | Degree of fruit color cover |
| 9 | 32475582 | bottom | AX-115619261 | Av_SAv_Sat_Abs_AB_High | Costa (2015) | Overcolor % |
| 9 | 32475582 | bottom | AX-115619261 | Av_SAv_Sat_Abs_AB_High | Amyotte et al. (2017) | Presence or absence of red skin colour |
| 9 | 32475582 | bottom | AX-115619261 | Av_SAv_Sat_Abs_AB_High | Kunihisa et al. (2014) | “Depth” of the red coloration |
| 9 | 32475582 | bottom | AX-115619261 | Av_SAv_Sat_Abs_AB_High | Migicovsky et al. (2021) | Red versus green |
| 9 | 32475582 | bottom | AX-115619261 | Av_SAv_Sat_Abs_AB_High | Duan et al. (2017) | Six levels of fruit skin color |
| 9 | 32475582 | bottom | AX-115619261 | Av_SAv_Sat_Abs_AB_High | Chagné et al. (2016) | Percentage overcolor |
| 9 | 32475582 | bottom | AX-115619261 | Av_SAv_Sat_Abs_AB_High | Migicovsky et al. (2016) | Overcolor and overcolor intesity |
| 9 | 32475582 | bottom | AX-115619261 | Av_SAv_Sat_Abs_AB_High | Lee et al. (2017) | Skin chroma (color intensity) |
| 9 | 32475582 | bottom | AX-115619261 | Av_SAv_Sat_Abs_AB_High | Kumar et al. (2014) | Overcolor on zero to nine scale |
| 9 | 32475582 | bottom | AX-115619261 | Av_SAv_Sat_Abs_AB_High | Minamikawa et al. (2021) | Degree of skin coloration in percent (1-5 scale) |
| 9 | 32475582 | bottom | AX-115619261 | Av_SAv_Sat_Abs_AB_High | Moriya et al. (2017) | Overall color intensity |
| 9 | 32475582 | bottom | AX-115619261 | Av_SAv_Sat_Abs_AB_High | McClure et al. (2018) | % red blush covering the surface |
| 9 | 32475582 | bottom | AX-115619261 | Av_SAv_Sat_Abs_AB_High | Zheng et al. (2020) | Degree of fruit color cover |
| 9 | 33729410 | bottom | AX-115332892 | SD_SAv_Hue_Abs_AB_Yellow | Costa (2015) | Overcolor % |
| 9 | 33729410 | bottom | AX-115332892 | SD_SAv_Hue_Abs_AB_Yellow | Amyotte et al. (2017) | Presence or absence of red skin colour |
| 9 | 33729410 | bottom | AX-115332892 | SD_SAv_Hue_Abs_AB_Yellow | Kunihisa et al. (2014) | “Depth” of the red coloration |
| 9 | 33729410 | bottom | AX-115332892 | SD_SAv_Hue_Abs_AB_Yellow | Migicovsky et al. (2021) | Red versus green |
| 9 | 33729410 | bottom | AX-115332892 | SD_SAv_Hue_Abs_AB_Yellow | Duan et al. (2017) | Six levels of fruit skin color |
| 9 | 33729410 | bottom | AX-115332892 | SD_SAv_Hue_Abs_AB_Yellow | Chagné et al. (2016) | Percentage overcolor |
| 9 | 33729410 | bottom | AX-115332892 | SD_SAv_Hue_Abs_AB_Yellow | Migicovsky et al. (2016) | Overcolor and overcolor intesity |
| 9 | 33729410 | bottom | AX-115332892 | SD_SAv_Hue_Abs_AB_Yellow | Lee et al. (2017) | Skin chroma (color intensity) |
| 9 | 33729410 | bottom | AX-115332892 | SD_SAv_Hue_Abs_AB_Yellow | Kumar et al. (2014) | Overcolor on zero to nine scale |
| 9 | 33729410 | bottom | AX-115332892 | SD_SAv_Hue_Abs_AB_Yellow | Minamikawa et al. (2021) | Degree of skin coloration in percent (1-5 scale) |
| 9 | 33729410 | bottom | AX-115332892 | SD_SAv_Hue_Abs_AB_Yellow | Moriya et al. (2017) | Overall color intensity |
| 9 | 33729410 | bottom | AX-115332892 | SD_SAv_Hue_Abs_AB_Yellow | McClure et al. (2018) | % red blush covering the surface |
| 9 | 33729410 | bottom | AX-115332892 | SD_SAv_Hue_Abs_AB_Yellow | Zheng et al. (2020) | Degree of fruit color cover |
| 9 | 33744382 | bottom | AX-115505043 | PC1_color | Amyotte et al. (2017) | Presence or absence of red skin colour |
| 9 | 33744382 | bottom | AX-115505043 | PC1_color | Kunihisa et al. (2014) | “Depth” of the red coloration |
| 9 | 33744382 | bottom | AX-115505043 | PC1_color | Kumar et al. (2014) | Overcolor on zero to nine scale |
| 9 | 33744382 | bottom | AX-115505043 | PC1_color | Minamikawa et al. (2021) | Degree of skin coloration in percent (1-5 scale) |
| 9 | 33744382 | bottom | AX-115505043 | PC1_color | Moriya et al. (2017) | Overall color intensity |
| 9 | 33744382 | bottom | AX-115505043 | PC1_color | McClure et al. (2018) | % red blush covering the surface |
| 9 | 33744382 | bottom | AX-115505043 | PC1_color | Zheng et al. (2020) | Degree of fruit color cover |
| 9 | 33744382 | bottom | AX-115505043 | PC1_color | Costa (2015) | Overcolor % |
| 9 | 33744382 | bottom | AX-115505043 | PC1_color | Chagné et al. (2016) | Percentage overcolor |
| 9 | 33744382 | bottom | AX-115505043 | PC1_color | Migicovsky et al. (2016) | Overcolor and overcolor intesity |
| 9 | 33744382 | bottom | AX-115505043 | PC1_color | Migicovsky et al. (2021) | Red versus green |
| 9 | 33744382 | bottom | AX-115505043 | PC1_color | Duan et al. (2017) | Six levels of fruit skin color |
| 9 | 33744382 | bottom | AX-115505043 | PC1_color | Lee et al. (2017) | Skin chroma (color intensity) |
| 9 | 33799120 | bottom | AX-105213720 | PC4_color | Jung et al. (2022) | Red over color |
| 9 | 33799120 | bottom | AX-105213720 | SD_SAv_Hue_Abs_AB_Red | Jung et al. (2022) | Red over color |
| 9 | 33799120 | bottom | AX-105213720 | PC4_color | Jung et al. (2022) | Red over color |
| 9 | 33799120 | bottom | AX-105213720 | SD_SAv_Hue_Abs_AB_Red | Jung et al. (2022) | Red over color |
| 9 | 33799120 | bottom | AX-115332892 | SD_SAv_Hue_Abs_AB_Yellow | Jung et al. (2022) | Red over color |
| 9 | 33799120 | bottom | AX-115505043 | PC1_color | Jung et al. (2022) | Red over color |
| 9 | 33801013 | bottom | AX-105213720 | PC4_color | Moriya et al. (2017) | Overall color intensity |
| 9 | 33801013 | bottom | AX-105213720 | SD_SAv_Hue_Abs_AB_Red | Moriya et al. (2017) | Overall color intensity |
| 9 | 33801013 | bottom | AX-105213720 | PC4_color | McClure et al. (2018) | % red blush covering the surface |
| 9 | 33801013 | bottom | AX-105213720 | SD_SAv_Hue_Abs_AB_Red | McClure et al. (2018) | % red blush covering the surface |
| 9 | 33801013 | bottom | AX-105213720 | PC4_color | Zheng et al. (2020) | Degree of fruit color cover |
| 9 | 33801013 | bottom | AX-105213720 | SD_SAv_Hue_Abs_AB_Red | Zheng et al. (2020) | Degree of fruit color cover |
| 9 | 33801013 | bottom | AX-105213720 | PC4_color | Costa (2015) | Overcolor % |
| 9 | 33801013 | bottom | AX-105213720 | SD_SAv_Hue_Abs_AB_Red | Costa (2015) | Overcolor % |
| 9 | 33801013 | bottom | AX-105213720 | PC4_color | Amyotte et al. (2017) | Presence or absence of red skin colour |
| 9 | 33801013 | bottom | AX-105213720 | SD_SAv_Hue_Abs_AB_Red | Amyotte et al. (2017) | Presence or absence of red skin colour |
| 9 | 33801013 | bottom | AX-105213720 | PC4_color | Kunihisa et al. (2014) | “Depth” of the red coloration |
| 9 | 33801013 | bottom | AX-105213720 | SD_SAv_Hue_Abs_AB_Red | Kunihisa et al. (2014) | “Depth” of the red coloration |
| 9 | 33801013 | bottom | AX-105213720 | PC4_color | Migicovsky et al. (2021) | Red versus green |
| 9 | 33801013 | bottom | AX-105213720 | SD_SAv_Hue_Abs_AB_Red | Migicovsky et al. (2021) | Red versus green |
| 9 | 33801013 | bottom | AX-105213720 | PC4_color | Duan et al. (2017) | Six levels of fruit skin color |
| 9 | 33801013 | bottom | AX-105213720 | SD_SAv_Hue_Abs_AB_Red | Duan et al. (2017) | Six levels of fruit skin color |
| 9 | 33801013 | bottom | AX-105213720 | PC4_color | Chagné et al. (2016) | Percentage overcolor |
| 9 | 33801013 | bottom | AX-105213720 | SD_SAv_Hue_Abs_AB_Red | Chagné et al. (2016) | Percentage overcolor |
| 9 | 33801013 | bottom | AX-105213720 | PC4_color | Migicovsky et al. (2016) | Overcolor and overcolor intesity |
| 9 | 33801013 | bottom | AX-105213720 | SD_SAv_Hue_Abs_AB_Red | Migicovsky et al. (2016) | Overcolor and overcolor intesity |
| 9 | 33801013 | bottom | AX-105213720 | PC4_color | Lee et al. (2017) | Skin chroma (color intensity) |
| 9 | 33801013 | bottom | AX-105213720 | SD_SAv_Hue_Abs_AB_Red | Lee et al. (2017) | Skin chroma (color intensity) |
| 9 | 33801013 | bottom | AX-105213720 | PC4_color | Kumar et al. (2014) | Overcolor on zero to nine scale |
| 9 | 33801013 | bottom | AX-105213720 | SD_SAv_Hue_Abs_AB_Red | Kumar et al. (2014) | Overcolor on zero to nine scale |
| 9 | 33801013 | bottom | AX-105213720 | PC4_color | Minamikawa et al. (2021) | Degree of skin coloration in percent (1-5 scale) |
| 9 | 33801013 | bottom | AX-105213720 | SD_SAv_Hue_Abs_AB_Red | Minamikawa et al. (2021) | Degree of skin coloration in percent (1-5 scale) |
| 9 | 33801013 | bottom | AX-105213720 | PC4_color | Moriya et al. (2017) | Overall color intensity |
| 9 | 33801013 | bottom | AX-105213720 | SD_SAv_Hue_Abs_AB_Red | Moriya et al. (2017) | Overall color intensity |
| 9 | 33801013 | bottom | AX-105213720 | PC4_color | McClure et al. (2018) | % red blush covering the surface |
| 9 | 33801013 | bottom | AX-105213720 | SD_SAv_Hue_Abs_AB_Red | McClure et al. (2018) | % red blush covering the surface |
| 9 | 33801013 | bottom | AX-105213720 | PC4_color | Zheng et al. (2020) | Degree of fruit color cover |
| 9 | 33801013 | bottom | AX-105213720 | SD_SAv_Hue_Abs_AB_Red | Zheng et al. (2020) | Degree of fruit color cover |
| 9 | 33801013 | bottom | AX-105213720 | PC4_color | Costa (2015) | Overcolor % |
| 9 | 33801013 | bottom | AX-105213720 | SD_SAv_Hue_Abs_AB_Red | Costa (2015) | Overcolor % |
| 9 | 33801013 | bottom | AX-105213720 | PC4_color | Amyotte et al. (2017) | Presence or absence of red skin colour |
| 9 | 33801013 | bottom | AX-105213720 | SD_SAv_Hue_Abs_AB_Red | Amyotte et al. (2017) | Presence or absence of red skin colour |
| 9 | 33801013 | bottom | AX-105213720 | PC4_color | Kunihisa et al. (2014) | “Depth” of the red coloration |
| 9 | 33801013 | bottom | AX-105213720 | SD_SAv_Hue_Abs_AB_Red | Kunihisa et al. (2014) | “Depth” of the red coloration |
| 9 | 33801013 | bottom | AX-105213720 | PC4_color | Migicovsky et al. (2021) | Red versus green |
| 9 | 33801013 | bottom | AX-105213720 | SD_SAv_Hue_Abs_AB_Red | Migicovsky et al. (2021) | Red versus green |
| 9 | 33801013 | bottom | AX-105213720 | PC4_color | Duan et al. (2017) | Six levels of fruit skin color |
| 9 | 33801013 | bottom | AX-105213720 | SD_SAv_Hue_Abs_AB_Red | Duan et al. (2017) | Six levels of fruit skin color |
| 9 | 33801013 | bottom | AX-105213720 | PC4_color | Chagné et al. (2016) | Percentage overcolor |
| 9 | 33801013 | bottom | AX-105213720 | SD_SAv_Hue_Abs_AB_Red | Chagné et al. (2016) | Percentage overcolor |
| 9 | 33801013 | bottom | AX-105213720 | PC4_color | Migicovsky et al. (2016) | Overcolor and overcolor intesity |
| 9 | 33801013 | bottom | AX-105213720 | SD_SAv_Hue_Abs_AB_Red | Migicovsky et al. (2016) | Overcolor and overcolor intesity |

| Chr | Position | Segment | Marker | Feature | Publication | Trait in reference |
|---|---|---|---|---|---|---|
| 9 | 33801013 | bottom | AX-105213720 | PC4_color | Lee et al. (2017) | Skin chroma (color intensity) |
| 9 | 33801013 | bottom | AX-105213720 | SD_SAv_Hue_Abs_AB_Red | Lee et al. (2017) | Skin chroma (color intensity) |
| 9 | 33801013 | bottom | AX-105213720 | PC4_color | Kumar et al. (2014) | Overcolor on zero to nine scale |
| 9 | 33801013 | bottom | AX-105213720 | SD_SAv_Hue_Abs_AB_Red | Kumar et al. (2014) | Overcolor on zero to nine scale |
| 9 | 33801013 | bottom | AX-105213720 | PC4_color | Minamikawa et al. (2021) | Degree of skin coloration in percent (1-5 scale) |
| 9 | 33801013 | bottom | AX-105213720 | SD_SAv_Hue_Abs_AB_Red | Minamikawa et al. (2021) | Degree of skin coloration in percent (1-5 scale) |
| 9 | 33801013 | bottom | AX-105213720 | PC4_color | Jung et al. (2022) | Red over color |
| 9 | 33801013 | bottom | AX-105213720 | SD_SAv_Hue_Abs_AB_Red | Jung et al. (2022) | Red over color |
| 9 | 33801013 | bottom | AX-105213720 | PC4_color | Jung et al. (2022) | Red over color |
| 9 | 33801013 | bottom | AX-105213720 | SD_SAv_Hue_Abs_AB_Red | Jung et al. (2022) | Red over color |
| 9 | 33801013 | bottom | AX-105213720 | PC4_color | Jung et al. (2022) | Red over color |
| 9 | 33801013 | bottom | AX-105213720 | SD_SAv_Hue_Abs_AB_Red | Jung et al. (2022) | Red over color |
| 9 | 33801013 | bottom | AX-105213720 | PC4_color | Jung et al. (2022) | Red over color |
| 9 | 33801013 | bottom | AX-105213720 | SD_SAv_Hue_Abs_AB_Red | Jung et al. (2022) | Red over color |
| 9 | 33801013 | bottom | AX-105213720 | PC4_color | Jung et al. (2022) | Red over color |
| 9 | 33801013 | bottom | AX-105213720 | SD_SAv_Hue_Abs_AB_Red | Jung et al. (2022) | Red over color |
| 9 | 33801013 | bottom | AX-105213720 | PC4_color | Jung et al. (2022) | Red over color |
| 9 | 33801013 | bottom | AX-105213720 | SD_SAv_Hue_Abs_AB_Red | Jung et al. (2022) | Red over color |
| 9 | 33801013 | bottom | AX-105213720 | PC4_color | Jung et al. (2022) | Red over color |
| 9 | 33801013 | bottom | AX-105213720 | SD_SAv_Hue_Abs_AB_Red | Jung et al. (2022) | Red over color |
| 9 | 33801013 | bottom | AX-105213720 | PC4_color | Jung et al. (2022) | Red over color |
| 9 | 33801013 | bottom | AX-105213720 | SD_SAv_Hue_Abs_AB_Red | Jung et al. (2022) | Red over color |
| 9 | 33801013 | bottom | AX-105213720 | PC4_color | Jung et al. (2022) | Red over color |
| 9 | 33801013 | bottom | AX-105213720 | SD_SAv_Hue_Abs_AB_Red | Jung et al. (2022) | Red over color |
| 9 | 33801013 | bottom | AX-105213720 | PC4_color | Jung et al. (2022) | Red over color |
| 9 | 33801013 | bottom | AX-105213720 | SD_SAv_Hue_Abs_AB_Red | Jung et al. (2022) | Red over color |
| 9 | 33801013 | bottom | AX-105213720 | PC4_color | Jung et al. (2022) | Red over color |
| 9 | 33801013 | bottom | AX-105213720 | SD_SAv_Hue_Abs_AB_Red | Jung et al. (2022) | Red over color |
| 9 | 33801013 | bottom | AX-115332892 | SD_SAv_Hue_Abs_AB_Yellow | Jung et al. (2022) | Red over color |
| 9 | 33801013 | bottom | AX-115332892 | SD_SAv_Hue_Abs_AB_Yellow | Jung et al. (2022) | Red over color |
| 9 | 33801013 | bottom | AX-115332892 | SD_SAv_Hue_Abs_AB_Yellow | Jung et al. (2022) | Red over color |
| 9 | 33801013 | bottom | AX-115332892 | SD_SAv_Hue_Abs_AB_Yellow | Jung et al. (2022) | Red over color |
| 9 | 33801013 | bottom | AX-115332892 | SD_SAv_Hue_Abs_AB_Yellow | Jung et al. (2022) | Red over color |
| 9 | 33801013 | bottom | AX-115505043 | PC1_color | Jung et al. (2022) | Red over color |
| 9 | 33801013 | bottom | AX-115505043 | PC1_color | Jung et al. (2022) | Red over color |
| 9 | 33801013 | bottom | AX-115505043 | PC1_color | Jung et al. (2022) | Red over color |
| 9 | 33801013 | bottom | AX-115505043 | PC1_color | Jung et al. (2022) | Red over color |
| 9 | 33801013 | bottom | AX-115505043 | PC1_color | Jung et al. (2022) | Red over color |
| 10 | 2507193 | top | AX-115228698 | SD_SSD_Sat_Abs_AB_Low | Duan et al. (2017) | Six levels of fruit skin color |
| 10 | 2507193 | top | AX-115228698 | SD_SSD_Sat_Abs_AB_Low | Zheng et al. (2020) | Degree of fruit color cover |
| 16 | 9244325 | top | AX-115244827 | PC2_color | Jung et al. (2022) | Ground color |
| 17 | 3096111 | top | AX-115394842 | SD_SAv_Hue_Abs_AB_Yellow | Jung et al. (2022) | Russet cover |
| 17 | 3769755 | top | AX-115452523 | PC1_color | Jung et al. (2022) | Red over color |
| 17 | 3823904 | top | AX-115452523 | PC1_color | Jung et al. (2022) | Red over color |

Table S4: Novel significant associations with principal components (PCs) and uncorrelated features for shape/size and color identified in this study. These markers have no overlapping positions with QTL reported in previous studies.

| Category | Chr | Position | Segment | Marker | Trait | Major effect |
|---|---|---|---|---|---|---|
| Size/shape | 1 | 15580877 | center | AX-115444766 | Av_SSD_X_Abs_A_Q25 | large size |
| Size/shape | 1 | 15580877 | center | AX-115444766 | Av_SSD_Pol_Abs_B.L_SD | large size |
| Size/shape | 1 | 15580877 | center | AX-115444766 | Av_SSD_X_Abs_A_Q25 | large size |
| Size/shape | 1 | 15580877 | center | AX-115444766 | Av_SSD_Pol_Abs_B.L_SD | large size |
| Size/shape | 1 | 15868916 | center | AX-115625798 | Av_SSD_X_Abs_B_Q25 | large size |
| Size/shape | 1 | 15918343 | center | AX-115538412 | PC5_shape_size | conical shape |
| Size/shape | 1 | 17152361 | center | AX-115221494 | Av_SSD_X_Abs_A_Q25 | large size |
| Size/shape | 1 | 20696409 | center | AX-115341839 | Av_SAv_Pol_SD_A.LR_Min | large size |
| Size/shape | 2 | 7626325 | top | AX-115378613 | Av_SAv_Pol_SD_A.LR_Min | large size |
| Size/shape | 2 | 7872781 | top | AX-115655116 | Av_SSD_Pol_SD_B.LR_Q25 | large size |
| Size/shape | 2 | 7904059 | top | AX-115490525 | PC2_shape_size | conical shape |
| Size/shape | 2 | 12130252 | top | AX-115534322 | PC1_shape_size | large size |
| Size/shape | 4 | 30194434 | bottom | AX-115506351 | Av_SAv_Pol_SD_A.LR_Min | large size |
| Size/shape | 6 | 4761377 | top | AX-115319264 | Av_SSD_X_Abs_B_Q25 | large size |
| Size/shape | 6 | 8743643 | top | AX-115375194 | Av_SAv_XY_Sum_A_MaxR | large size |
| Size/shape | 6 | 10584742 | top | AX-115492426 | Av_SSD_Pol_SD_B.LR_Max | conical shape |
| Size/shape | 7 | 21705520 | center | AX-115254276 | PC3_shape_size | conical shape |
| Size/shape | 9 | 4698417 | top | AX-115442384 | Av_SSD_X_Abs_A_Q25 | large size |
| Size/shape | 9 | 5170431 | top | AX-115521252 | Av_SSD_X_Abs_B_Q25 | large size |
| Size/shape | 14 | 1478099 | top | AX-115628310 | PC1_shape_size | large size |
| Size/shape | 14 | 15862103 | center | AX-115363036 | Av_SSD_X_Abs_A_Q25 | large size |
| Size/shape | 14 | 21571993 | center | AX-115311704 | Av_SSD_Pol_Rat.Av_AB.LR_Min | large size |
| Color | 3 | 6282194 | top | AX-115321298 | PC1_color | |
| Color | 10 | 38213067 | bottom | AX-115182287 | PC1_color | |
| Color | 10 | 41138070 | bottom | AX-115536962 | SD_SSD_Sat_Abs_AB_Low | |

Table S5: Significant marker-feature associations with each principal component (PC) 1 to 5 for shape were assembled into haplotypes and divided into distinct groups using hierarchical clustering. The cluster number, the haplotype composition, the number of genotypes in each cluster (N) as well as the SNP markers constituting to each cluster are given.

See file Table_S5.csv

#### 2 Supplemental figures

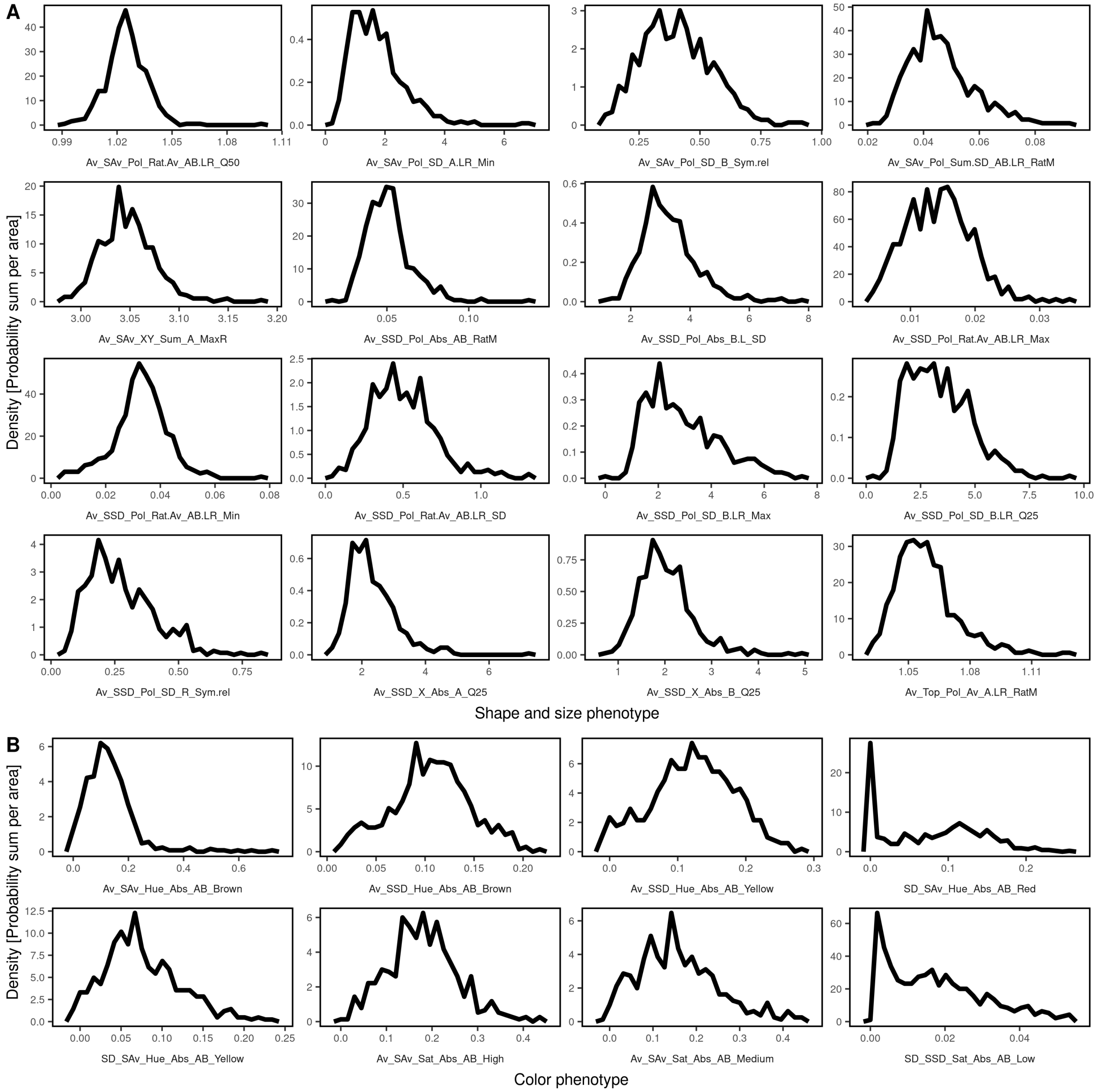


Figure S1: Kernel density estimate of the selected uncorrelated fruit features. A: Uncorrelated features for shape and size. B: Uncorrelated color features. The estimate is a smoothed histogram with the integral equal to one calculated using a Gaussian kernel and a binwidth of 1/30 of the data range. See Table 1 for feature abbreviations and description.

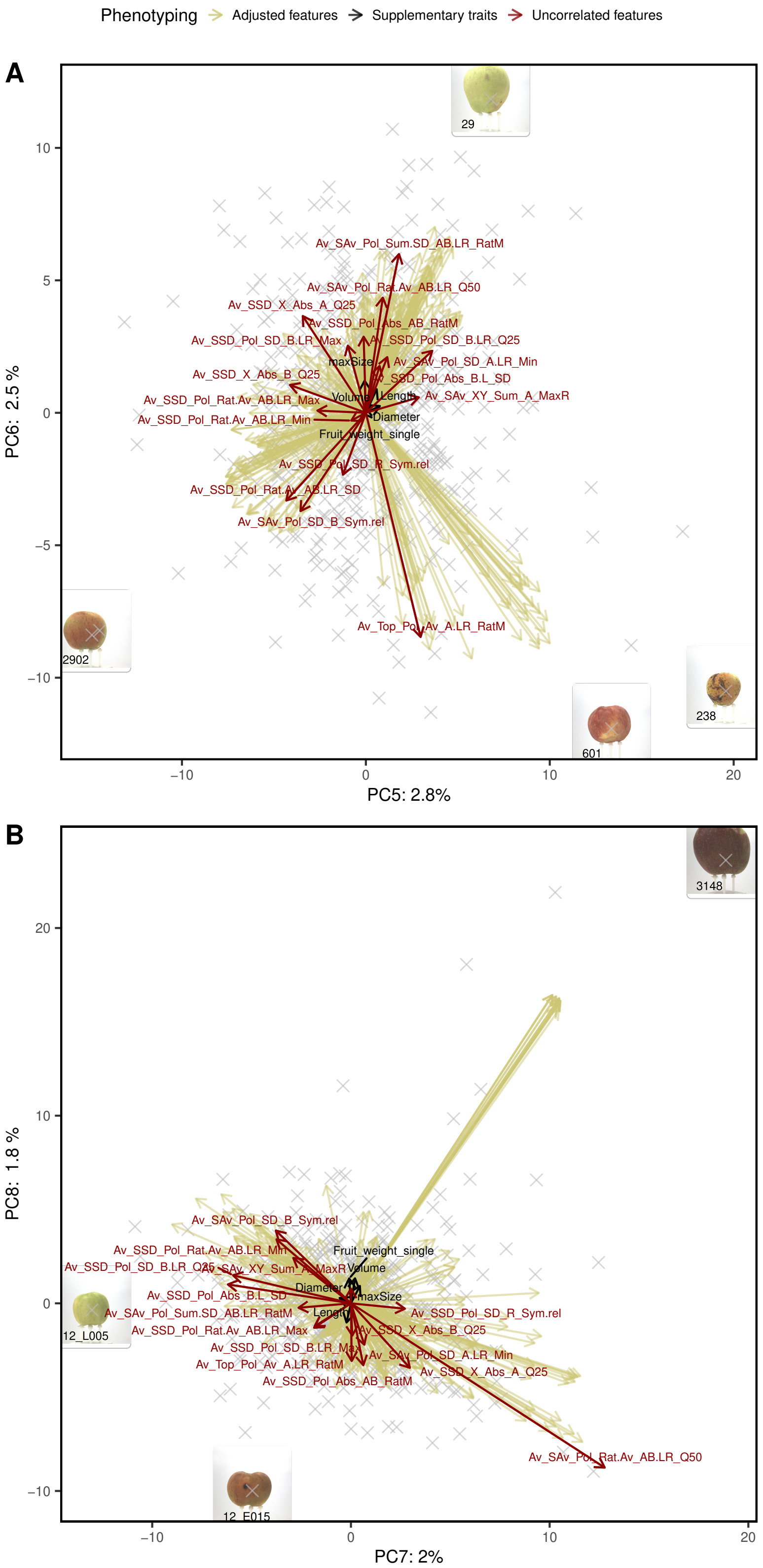


Figure S2: Principal component (PC) analysis of highly heritable features for fruit shape and size. Uncorrelated ($r < 0.75$) fruit shape and size features (red arrows) and supplementary features measured using the sorting machine (black arrows) were overlayed in a biplot. Images of extreme genotypes of the apple REFPOP were shown in boxes and labeled with their respective genotype codes. A: PC5 and PC6, B: PC7 and PC8.

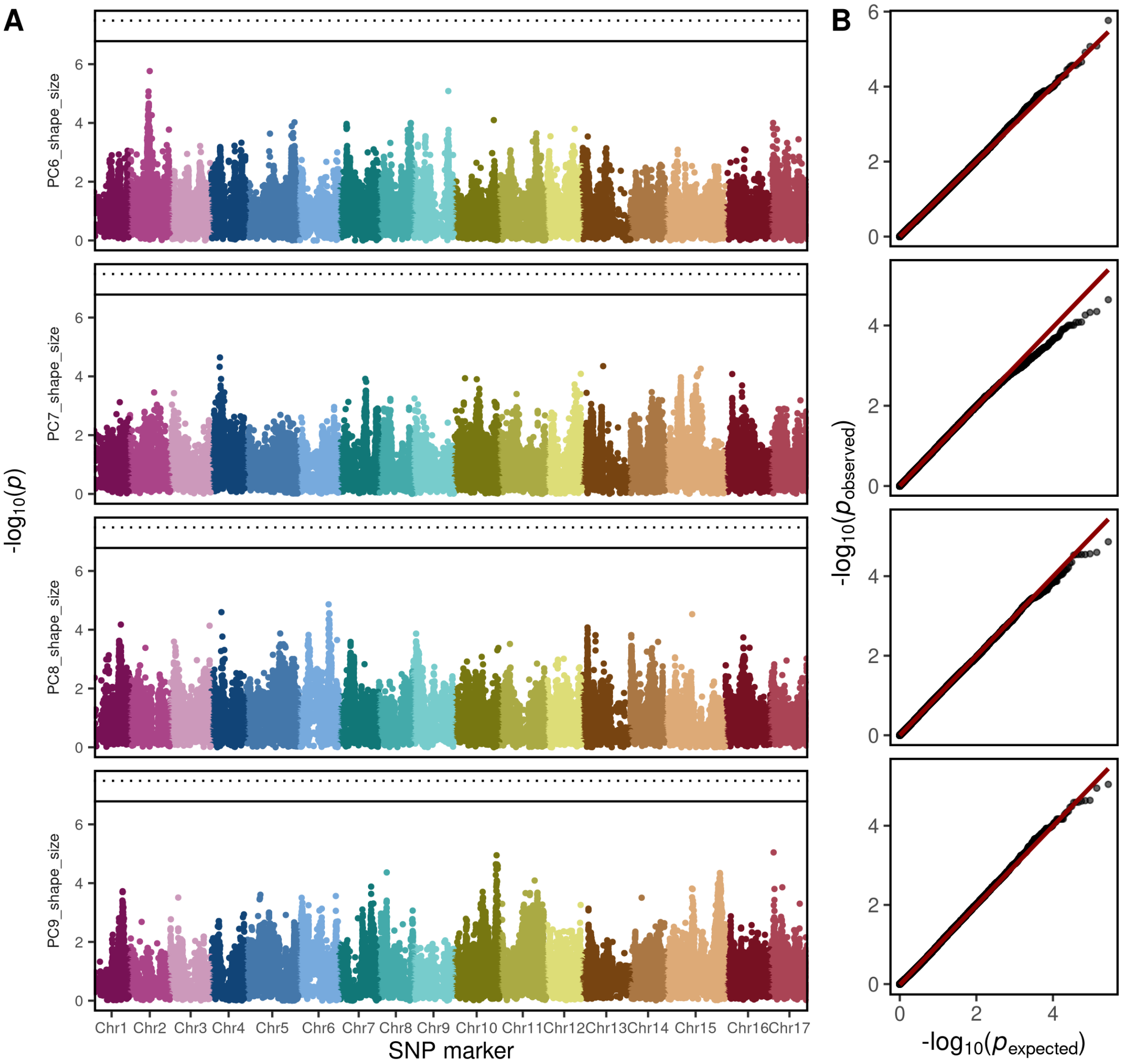


Figure S3: Results of the genome-wide association studies for the principal component (PC) 6 to 9 build from the highly heritable fruit shape and size features. A: Manhattan plots with the $-log_{10}(p)$ value for every SNP marker and PC. The SNP markers are displayed according to their physical position on the genome and chromosome (Chr). The Bonferroni-corrected significance level is indicated at 5% and 1% by the solid and dotted line, respectively. B: The expected and observed p-values for SNP markers and PCs. The 1:1 line for non-significant p-values is shown in red.

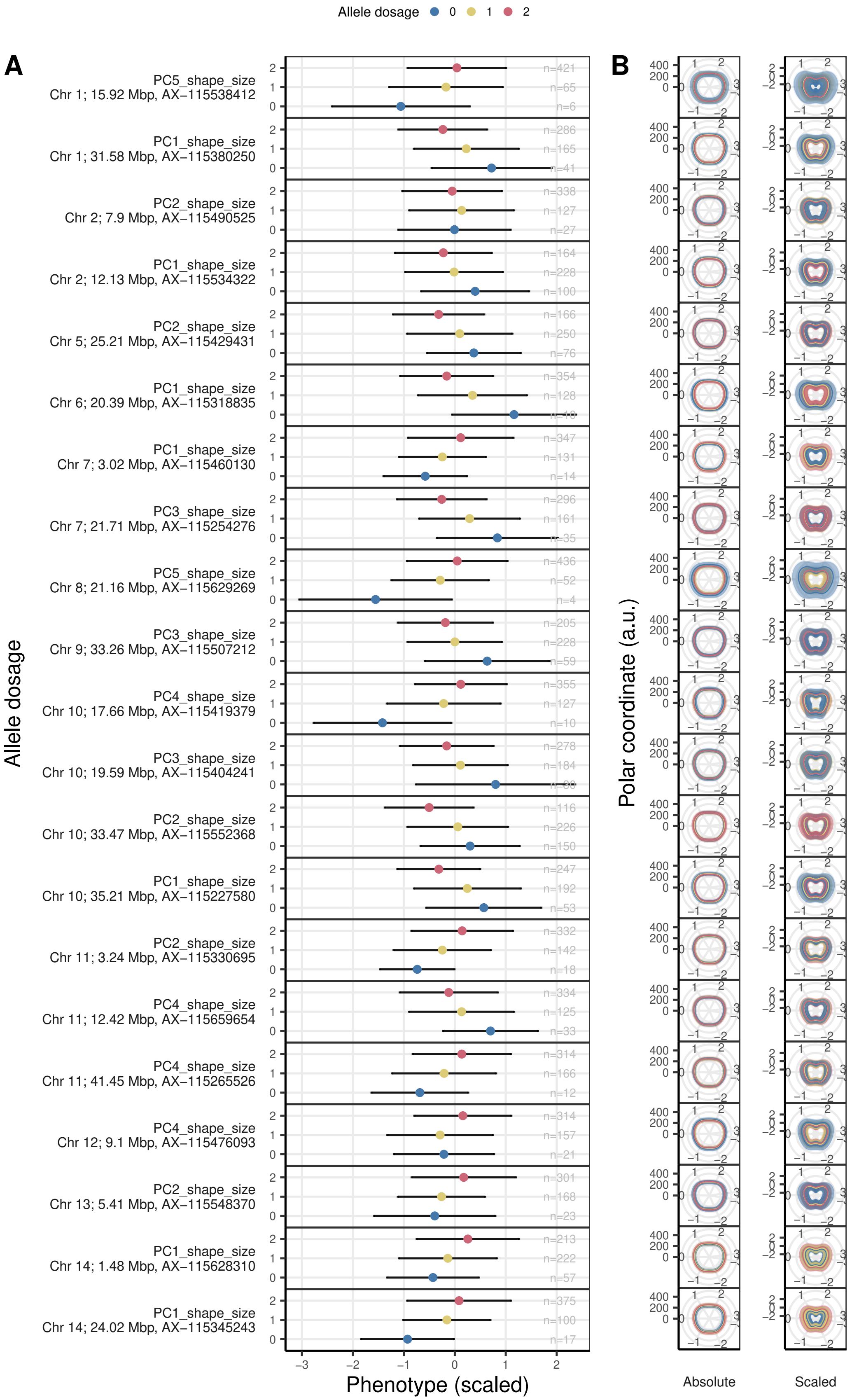


Figure S4: Effect of SNPs significantly associated with principal component (PC) 1 to 5 for fruit shape and size. A: Effect of the allele dosage (0, 1 or 2 alternative alleles) on the scaled values of PCs. B: The effect of each SNP on the averaged side-view fruit contour in absolute and relative values. The error bars show the standard deviation.

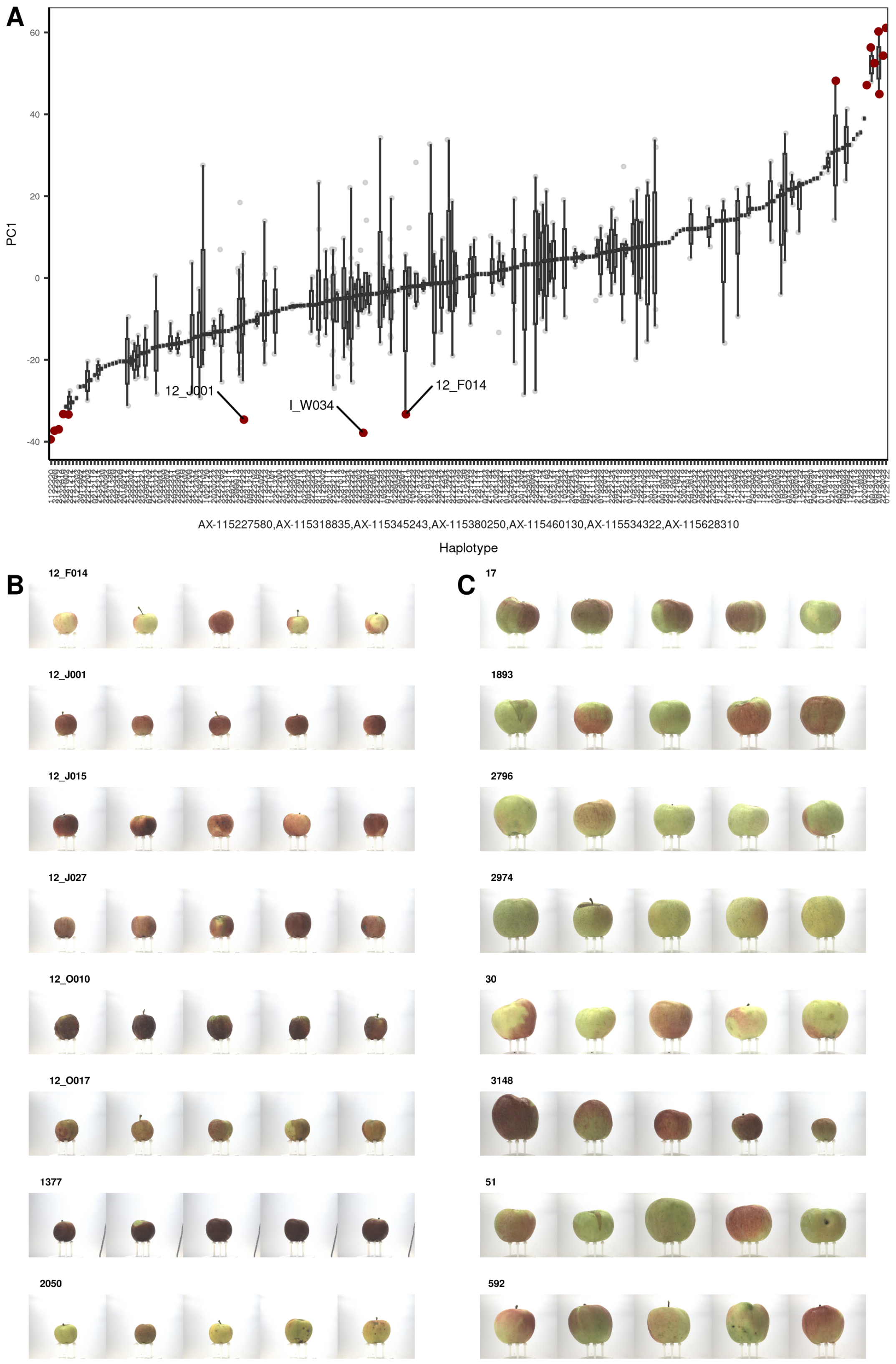


Figure S5: Combined effect of the SNPs identified for principal component 1 (PC1) on the size of apples. The SNPs were assembled into haplotypes. A: Boxplots of the effect of each haplotype on PC1. The haplotypes are labeled using combinations of allele dosage (0, 1 or 2 alternative alleles) of the SNPs. The eight most extreme phenotypes for low and high phenotypic trait values are depicted in red. Their fruit images with up to five replicates are shown in B: and C:, respectively.

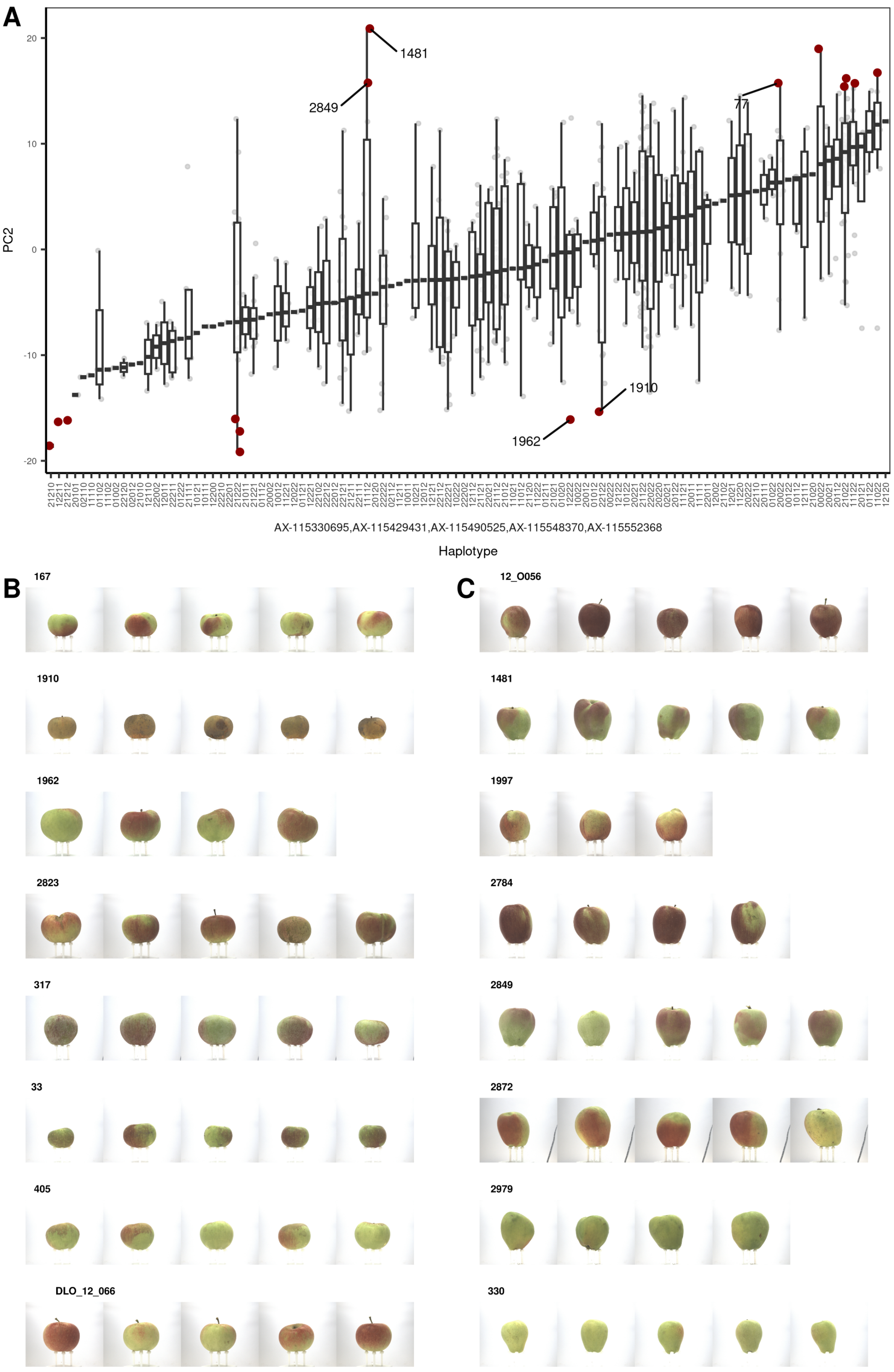


Figure S6: Combined effect of the SNPs identified for principal component 2 (PC2) on the shape of apples. The SNPs were assembled into haplotypes. A: Boxplots of the effect of each haplotype on PC1. The haplotypes are labeled using combinations of allele dosage (0, 1 or 2 alternative alleles) of the SNPs. The eight most extreme phenotypes for low and high phenotypic trait values are depicted in red. Their fruit images with up to five replicates are shown in B: and C:, respectively.

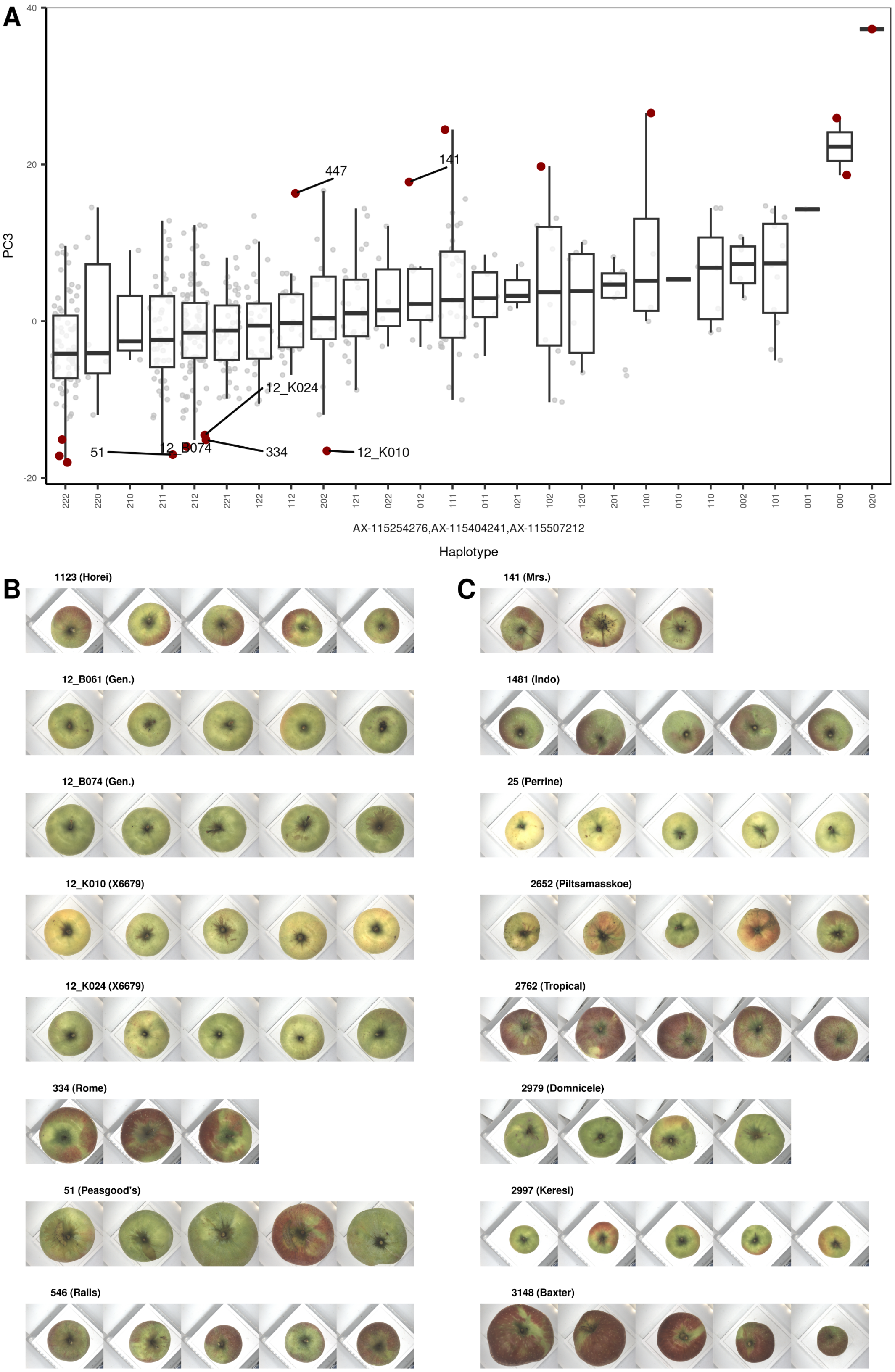


Figure S7: Combined effect of the SNPs identified for principal component 3 (PC3) on the shape of apples. The SNPs were assembled into haplotypes. A: Boxplots of the effect of each haplotype on PC1. The haplotypes are labeled using combinations of allele dosage (0, 1 or 2 alternative alleles) of the SNPs. The eight most extreme phenotypes for low and high phenotypic trait values are depicted in red. Their fruit images with up to five replicates are shown in B: and C:, respectively.

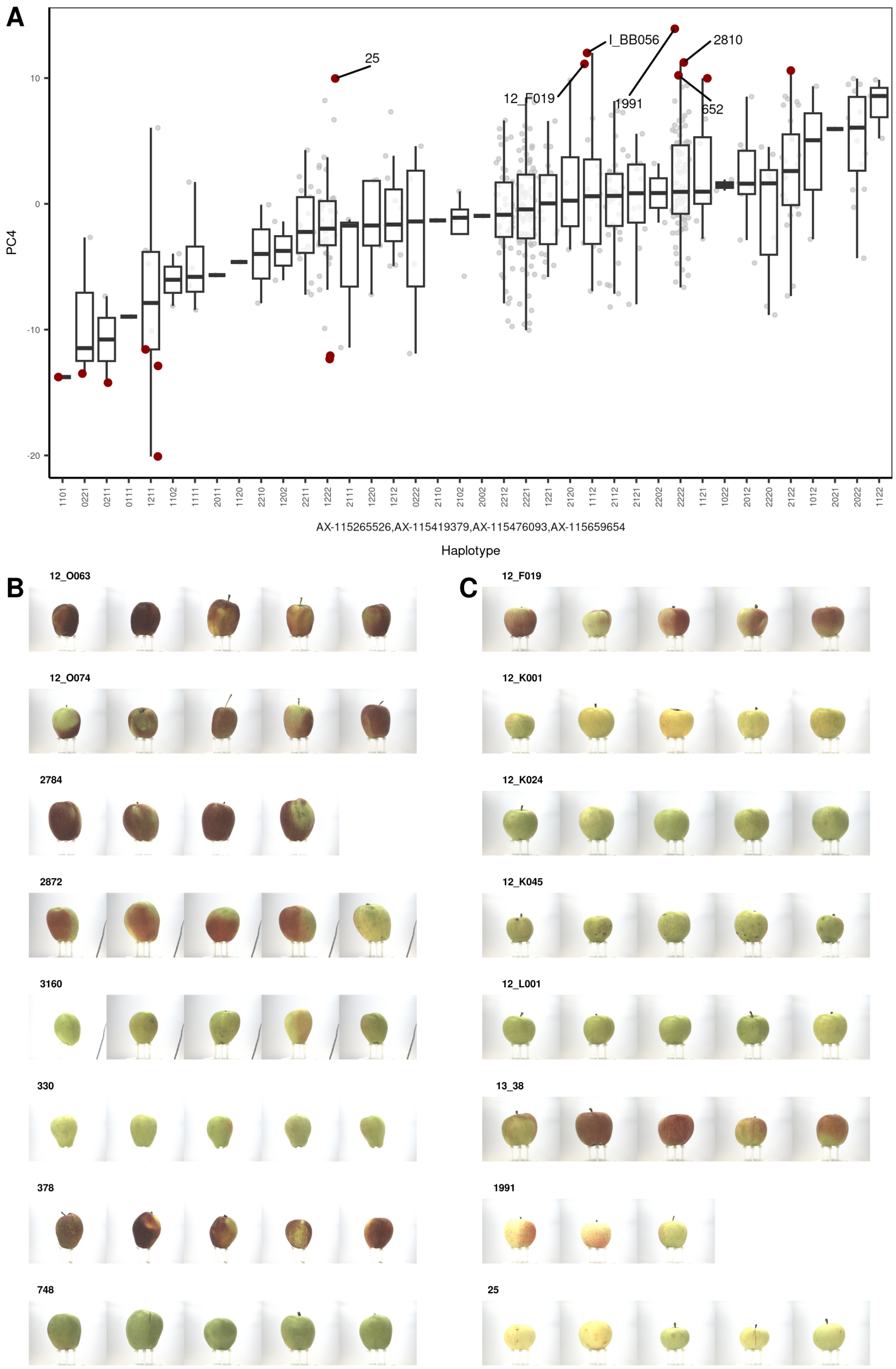


Figure S8: Combined effect of the SNPs identified for principal component 4 (PC4) on the shape of apples. The SNPs were assembled into haplotypes. A: Boxplots of the effect of each haplotype on PC1. The haplotypes are labeled using combinations of allele dosage (0, 1 or 2 alternative alleles) of the SNPs. The eight most extreme phenotypes for low and high phenotypic trait values are depicted in red. Their fruit images with up to five replicates are shown in B: and C:, respectively.

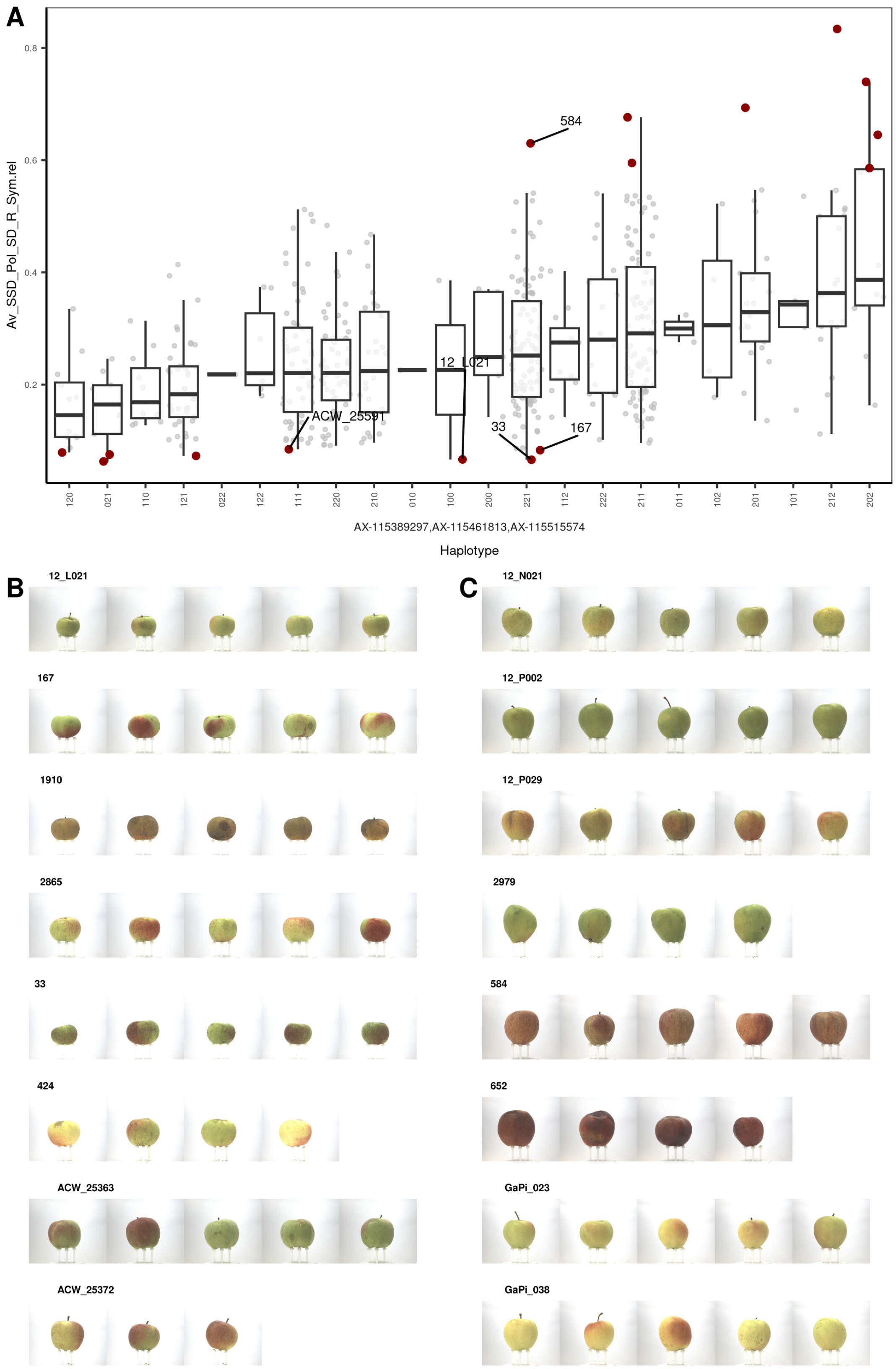


Figure S9: Effect of haplotypes build from SNPs AX-115389297, AX-115461813, and AX-115515574 that were significantly associated with the uncorrelated symmetry feature Av_SSD_Pol_SD_R_Sym.rel on the symmetry of apple fruits. A: Boxplots of the haplotype effects on the symmetry feature. The haplotypes are labeled using combinations of allele dosage (0, 1 or 2 alternative alleles) of the SNPs. The eight most extreme phenotypes for low and high phenotypic trait values are depicted in red. Their fruit images with up to five replicates are shown in B: and C:, respectively.

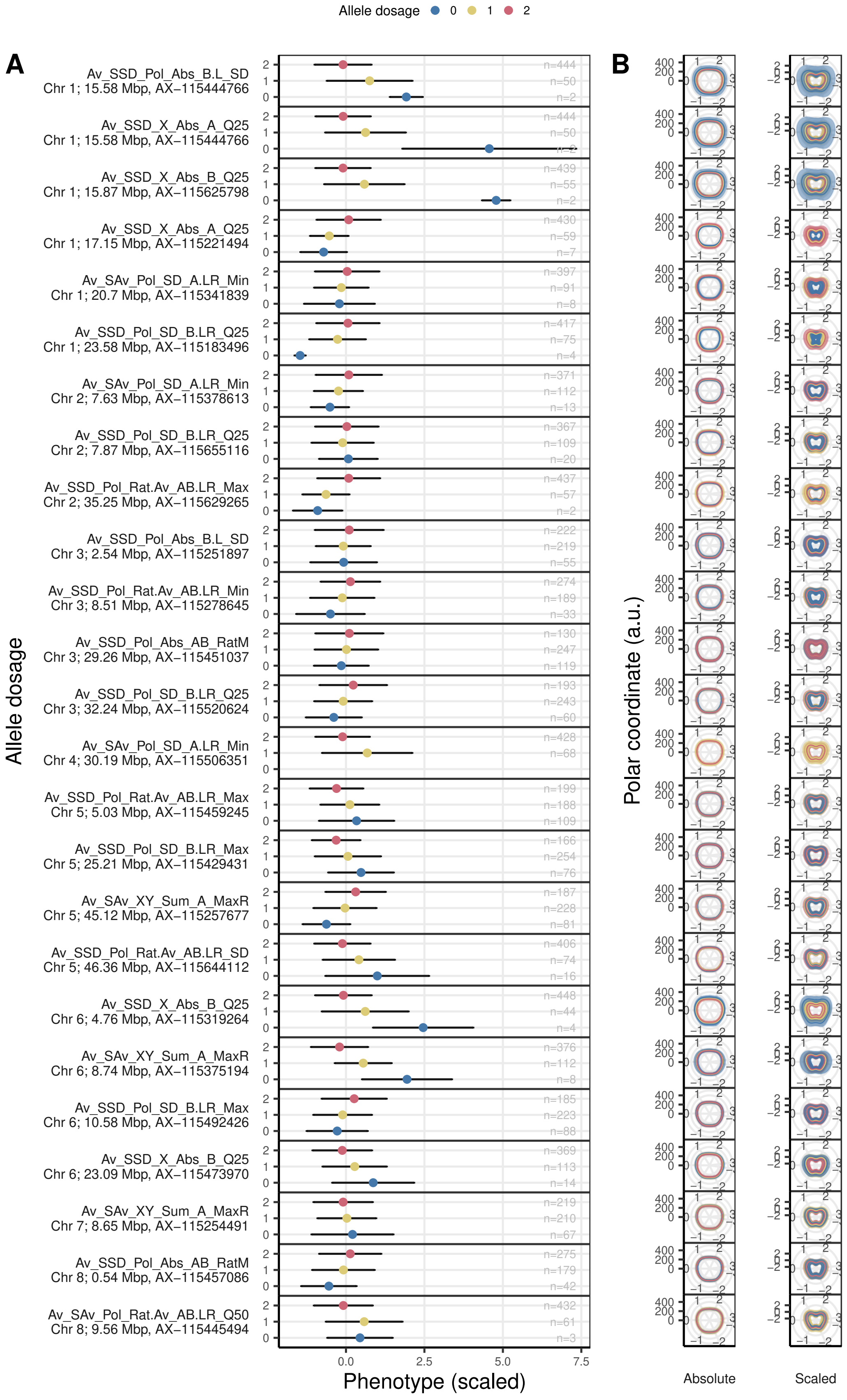


Figure continues on next page.

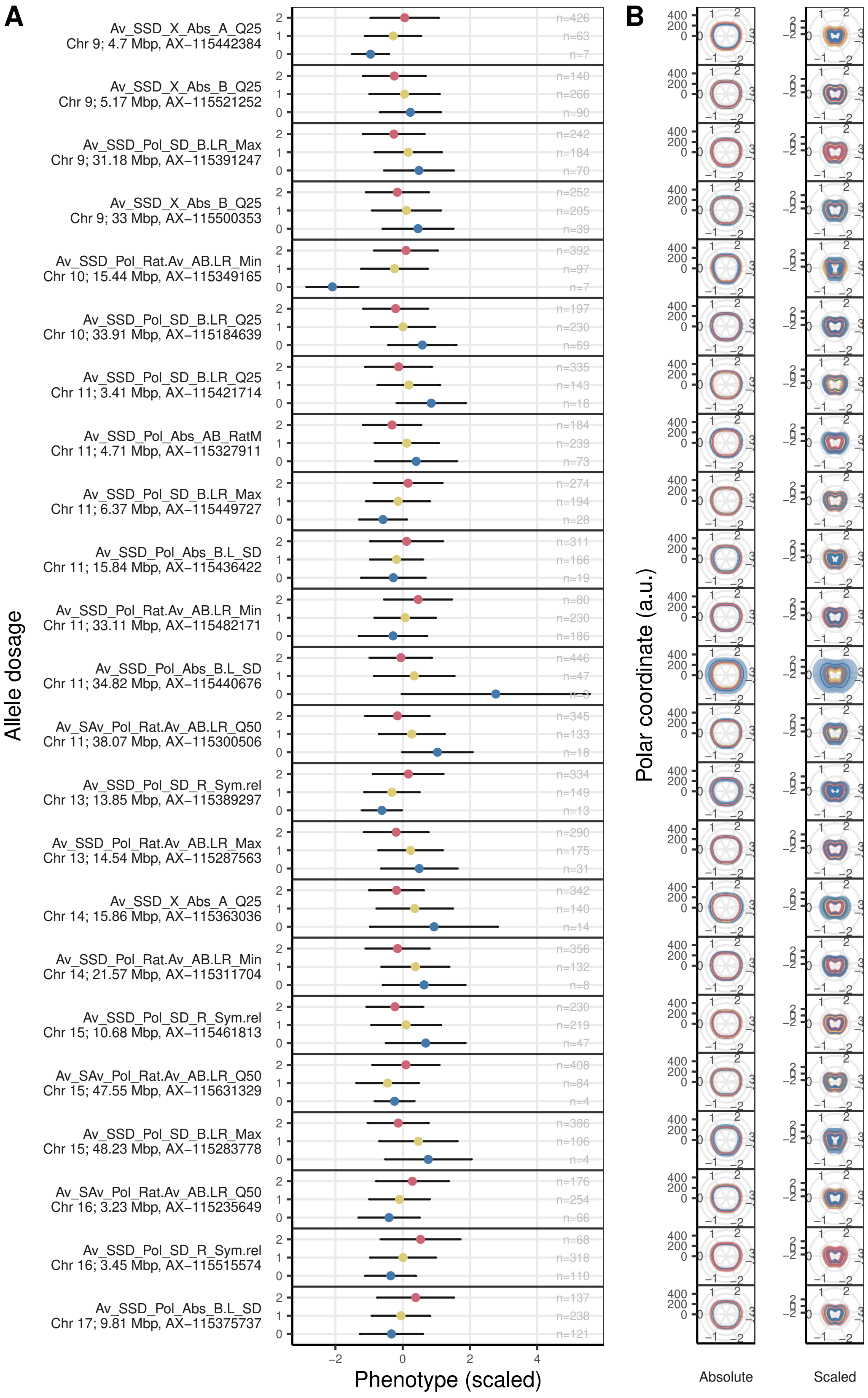


Figure S10: Effect of SNPs significantly associated with uncorrelated features for fruit shape and size. A: Effect of the allele dosage (0, 1 or 2 alternative alleles) on the scaled values of PCs. B: The effect of each SNP on the averaged side-view fruit contour in absolute and relative values. The error bars show the standard deviation. See previous page to view the the full figure.

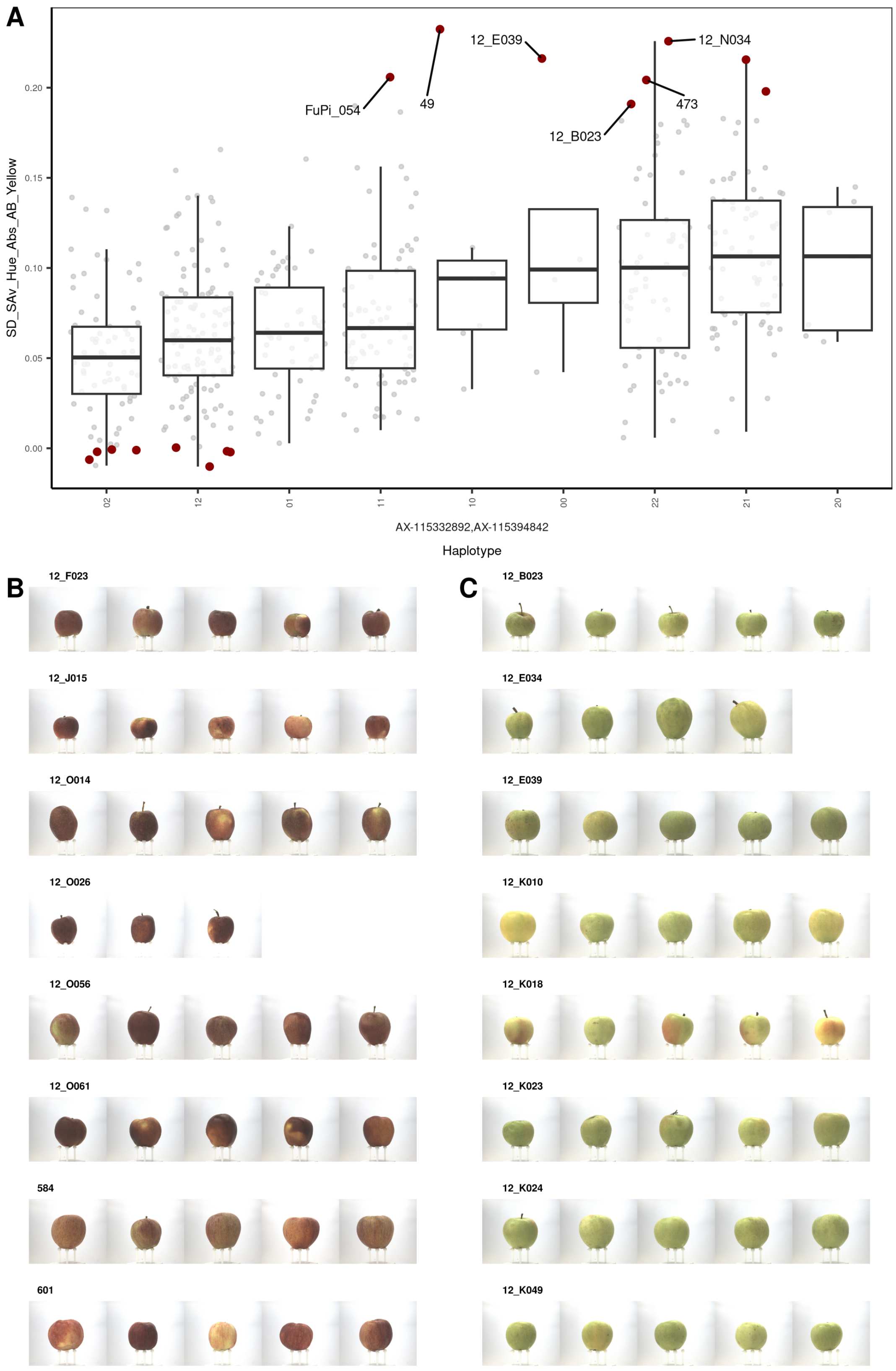


Figure S11: Effect of haplotypes build from SNPs that were significantly associated with the uncorrelated color feature SD_SAv_Hue_Abs_AB_Yellow on the color of apple fruits. A: Boxplots of the haplotype effects on the color feature. The haplotypes are labeled using combinations of allele dosage (0, 1 or 2 alternative alleles) of the SNPs. The eight most extreme phenotypes for low and high phenotypic trait values are depicted in red. Their fruit images with up to five replicates are shown in B: and C:, respectively.

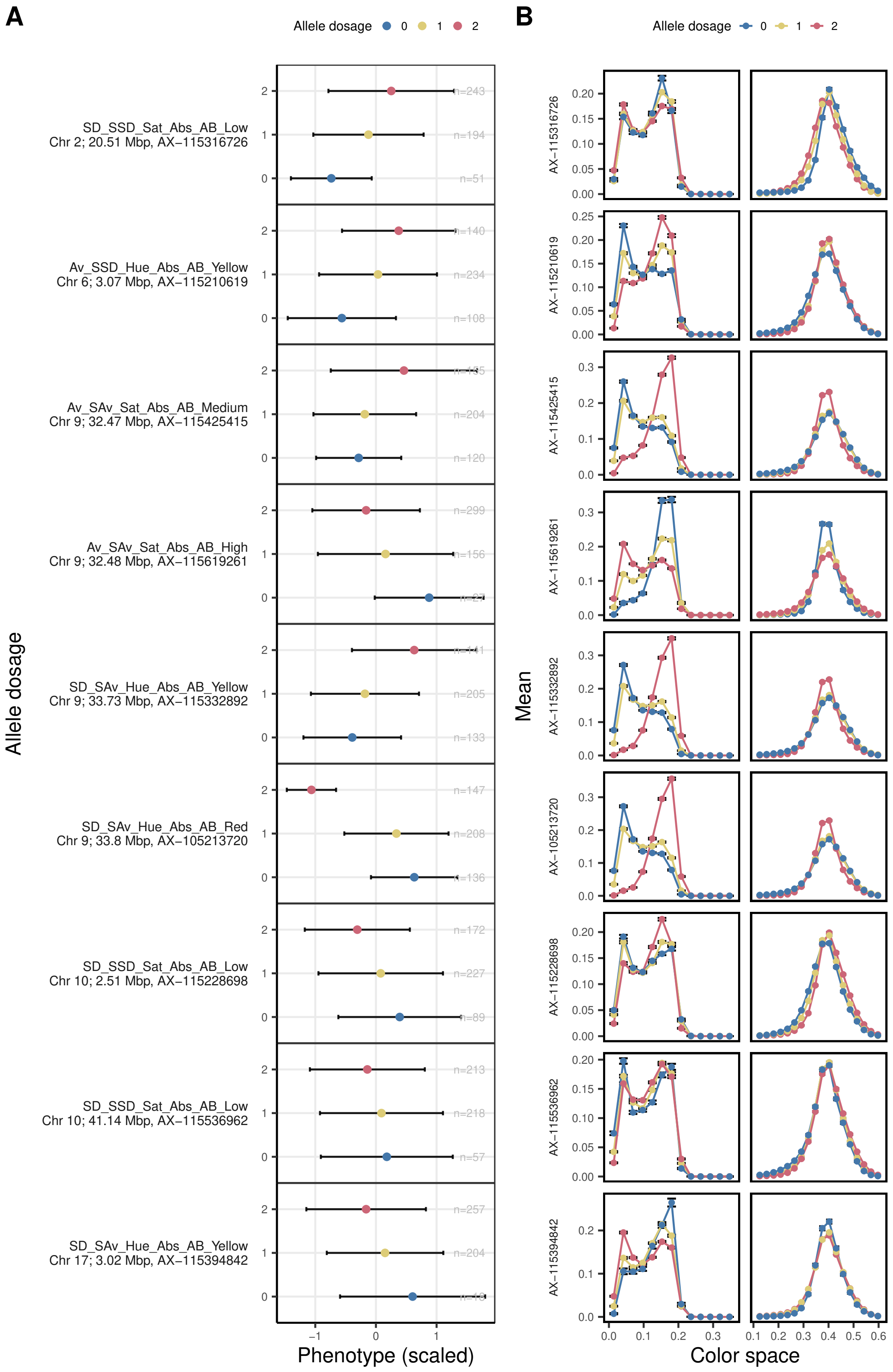


Figure S12: Allele dosage effect (0, 1 or 2 alternative alleles) on the hue and saturation color space for the uncorrelated color features. The error bars show the standard deviation of all genotypes per allele dosage. A: The position, the name of the SNP marker and the effect of the allele dosage is shown (0, 1 or 2 alternative alleles). The phenotypes were scaled for each feature to allow for a relative comparison. The error bar shows the standard deviation. B: The effect of SNPs on the hue and saturation color space. The error bars show the standard deviation of all genotypes per allele dosage. The Bonferroni-corrected significance level of the considered associations was 5%.
